# A MultiOmics time-resolved systems map of gene regulation in *Toxoplasma gondii*

**DOI:** 10.64898/2026.10.29.762658

**Authors:** Yihan Wu, Argenis A. Arriojas, Nachiket Shankar, Shalini Aggarwal, Jingjing Lou, Prarthana Prathap, Shreya Dua, Gretchen M. Diffendall, Jiahong Li, Pamodya Pamunuwa, Prince Oppong, Omar Graia, Lee Sarah Izhaki-Tavor, Eranthie Weerapana, Mélissa Léger-Abraham, Manoj T. Duraisingh, Kourosh Zarringhalam, Marc-Jan Gubbels

## Abstract

Understanding how *T. gondii* builds an invasion-competent cell has been limited by its rapid, asynchronous division cycle, incomplete gene function annotation, and studies that have examined one regulator or complex at a time. Here, we placed acute perturbation of 21 transcriptional and epigenetic factors, promoter occupancy, chromatin accessibility, and molecular validation on a common single-cell clock. The resulting consensus map links the lytic-cycle programs to candidate activating and repressive inputs, organizes genes into 108 co-regulated communities, and nominates regulatory partnerships that would not have emerged from any single data type. The work bridges three longstanding gaps: it resolves perturbation responses in cell-cycle time, distinguishes promoter occupancy from perturbation-linked regulation, and turns a large multidimensional dataset into an experimentally navigable map. We indeed confirmed known factor interactions such as MORC/HDAC3, and recovered combinations such as the Cdc5/SNIP1 and DMAP1/EPL1 as coregulatory nodes (that in other systems have been described to cooperate). Furthermore, we discovered a novel apicomplexan specific combination of AAF1/AP2XI-5 that provides mechanistic insights of the unique AP2 transcription factor family.

## 1 Introduction

Apicomplexan parasites have major impacts on human health, for example malaria is caused by *Plasmodium falciparum*, whereas *Toxoplasma gondii* causes opportunistic infections. Pathology is driven by repetitive lytic replication cycles that rapidly expand parasite biomass and destroy host tissues. These parasites share an obligate intracellular lifestyle and proliferate through unconventional budding modes, with each replicative cycle beginning and ending with an invasion-competent ‘zoite’ [1]. The rapidly proliferating *T. gondii* tachyzoite replicates through endodyogeny, assembling two daughters within the mother before host cell egress and reinvasion [2]. Repetition of this cycle drives tissue injury during acute infection and recrudescence [3,4]. This process is highly ordered but departs from familiar yeast and mammalian cell cycles in several ways. For example, DNA replication and mitosis progress in parallel with daughter budding and organelle biogenesis (**Fig. 1A**) [5,6]. The parasite must therefore build each component when it is needed, even as one cellular operation begins before another has finished.

**Figure 1.**
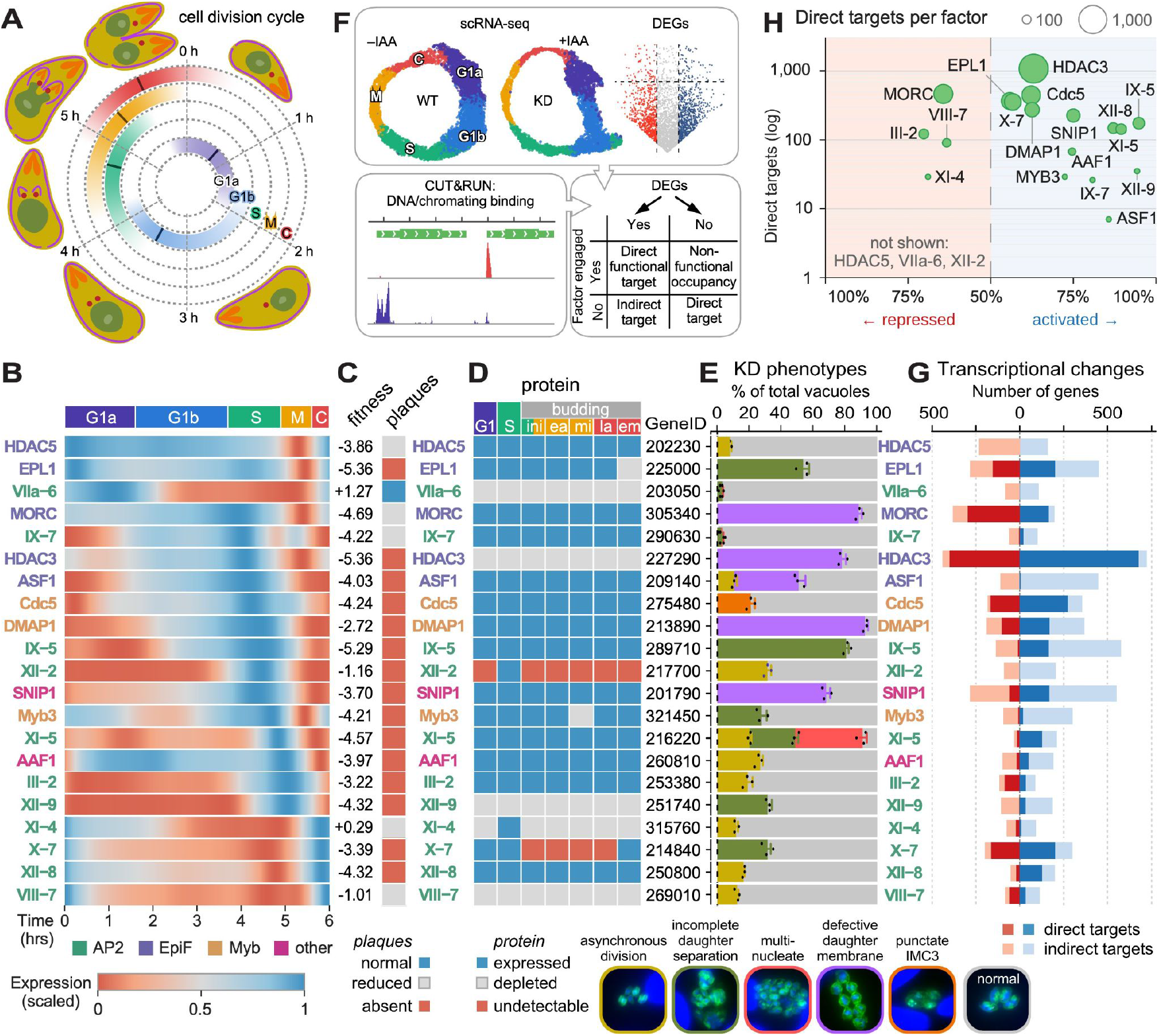
Approach and diversity of DNA binding and epigenetic factor genetic perturbation outcomes/phenotypes/response. **(A)** Schematic representation of the *T. gondii* cell division cycle. Cell cycle stage density derived from scRNA-seq data converted to pseudotime. Cell-cycle phase scores of wild-type (RH) cells along the tachyzoite cell cycle (0–6 h, clockwise from 0 h at the top): one ring per phase, G1a (inside) to C (outside), shaded from white to the phase color by the fitted phase-marker score (negative scores clipped, one shared maximum); the dark tick on each ring marks that phase’s peak. **(B)** Normalized mRNA expression profiles of the 21 factors selected for knock-down along the 6 hrs tachyzoite cell cycle. Rows are ordered by peak time and profiles are scaled per gene from 0 to 1 (low = red, high = blue); top bar marks cell-cycle phases. See **Fig. S20D** for absolute expression levels. **(C)** Plaque formation capacity for all cKD lines over 8 days growth in presence of IAA. ‘Reduced’ plaqueing captures both reduced plaque size and reduced number. See **Fig. S3** for the experimental data. Fitness scores are from [31]. **(D)** Protein expression profiles of the factors as assessed by immunofluorescence using the Ty epitope fusion (see **Fig. S4** for the experimental data from which the interpretations are derived). Budding progression defined as follows: ini: initiation; ea: early; mi: mid; la: late; em: emerging. GeneIDs represent the ToxoDB.org TGME49_ or TGGT1_ accession numbers. **(E)** Distribution of cell division and cytoskeleton defects scored across cKD mutants upon 6h factor depletion. Host cells were infected with the indicated parasite strains and incubated overnight, followed by treatment with IAA for 6 h. Immunofluorescence at the bottom represents the phenotypes scored by DAPI (marking nucleus) and IMC3 (marking the budding and mature cortical cytoskeleton) co-staining. Average of n=3+std 3 biological replicates plotted. 100 vacuoles counted in each replicate. Grey bar sections indicate the fraction in which “no defect was seen”. **(F)** Schematic representation of the basic genomics data collection and analysis approach. Each factor was genetically perturbed using mini-auxin inducible degron (mAID) fusion (genotype validations in **Fig. S2**). For each factor, single cell transcriptional data upon knock-down were collected as well as DNA association data by CUT&RUN under the permissive condition. **(G)** Integration of scRNA-seq data (treated as bulk) and CUT&RUN data for each factor, reporting the number of direct up- and down-regulated genes differentiated by direct (factor engagement of the gene seen by CUT&RUN) and indirect (factor not bound to gene). Transcriptional changes upon knock-down: genes differentially expressed (pooled pseudobulk knock-down vs. wild type; FDR < 0.05, |log2FC| > 1, expressed in > 5% of cells in either condition) that are down-regulated in the knock-down (activated targets; blue, right) or up-regulated (repressed targets; red, left), split into direct targets (CUT&RUN peak overlapping the TSS ± 1 kb, q < 0.075; dark) and indirect targets (light). We detect fewer than 25 direct targets for HDAC5, AP2VIIa-6 and AP2XII-2. **(H)** Direct targets per factor: direction balance vs. number of direct targets; circle area ∝ number of direct targets. Factors with fewer than 25 CUT&RUN targets are not shown.

Gene expression throughout this division cycle follows a circular, “just-in-time” principle. Landmark *P. falciparum* microarray studies first showed that genes rise and fall in a once-per-cycle order, a discovery that transformed parasite biology [7–9]. In *T. gondii*, transcriptome profiling revealed a comparable cascade linking biosynthesis, genome replication, daughter construction, and invasion [10]. Single-cell transcriptomic analysis further showed that asynchronous parasite populations span a continuous progression of transcriptional states rather than a small set of discrete stages [11]. Matched single-cell RNA and chromatin-accessibility atlases connected these expression waves to periodic promoter opening and identified coherent co-regulated gene sets [6]. Together,hese studies established when genes are expressed. How a compact repertoire of unconventional transcription factors and epigenetic regulators acts across the changing promoter landscape to coordinate the ordered gene-expression programs that drive the division cycle remains unresolved.

To coordinate gene-expression programs, apicomplexans rely mainly on ApiAP2 and Myb transcription factors. The ApiAP2s constitute the largest family and are key drivers of the cascading expression waves that propel lytic-cycle progression [5]. This lineage-expanded family was first identified amid an otherwise sparse collection of conventional eukaryotic DNA-binding domains in Apicomplexa [12]. Protein-binding microarray studies of *P. falciparum* ApiAP2 domains showed that they recognize defined DNA sequences, that orthologous domains in distant apicomplexans retain these preferences, and that individual proteins can bind more than one motif [13,14]. Functional studies in *T. gondii* have assigned several ApiAP2 factors to sequential or cooperative roles in lytic-cycle regulation. AP2IX-5 initiates the budding transcriptional program and activates AP2XII-9 [15], which subsequently controls aspects of daughter-cell construction [16], whereas AP2XI-5 cooperates with AP2X-5 to regulate invasion-organelle genes [17,18]. AP2XII-8 controls ribosomal protein genes required for progression through G1 [6,reviewed in 5,2]. Transcriptional regulation also extends beyond the lytic cycle. The Myb-like factor BFD1 drives tachyzoite-to-bradyzoite differentiation [4]. Single-cell analysis of brain-derived bradyzoites revealed an in vivo-specific cell cycle in which AP2XI-6 is required for encystation [19].

ApiAP2 factors also cooperate with epigenetic regulators. In *T. gondii*, MORC and HDAC3 act with ApiAP2 partners to repress sexual-stage programs and maintain merozoite genes in a poised state, whereas two distinct SWI/SNF remodeling complexes coordinate transcriptional timing across the cell cycle [20–24]. ApiAP2 proteins also associate with histone-modifying and reading machinery and with other ApiAP2s, providing direct precedents for combinatorial control [25,26]. Together with evidence for cooperative [18] and opposing, potentially competing ApiAP2 activities [27], these studies suggest that ApiAP2 factors operate within interconnected regulatory networks rather than as isolated regulators. How these networks are organized across the lytic cycle remains unresolved. Work in other eukaryotes provides a broader framework: recognizable motifs create potential binding sites, but factor occupancy and regulatory output also reflect chromatin accessibility, local motif arrangement, core-promoter properties, and interactions among factors [28–30]. Although studies in *T. gondii* have begun to connect transcriptional and chromatin regulation, most have examined one regulator, transition, or complex at a time, leaving their organization at network scale unclear.

Genome-scale resources have transformed the functional study of *T. gondii*. Genetic screens identify fitness-conferring genes, and spatial proteomics locates thousands of proteins within the parasite [31,32]. In *P. falciparum*, ChIP-seq profiling of 18 ApiAP2 factors across three bloodstage intervals, integrated with bulk RNA-seq from six ApiAP2 knockout lines, defined an occupancy-centered network focused on heterochromatin and variant-gene expression [33]. However, occupancy alone does not by itself identify responsive genes, and bulk measurements average over the division cycle, obscuring when a response occurs. *T. gondii* still lacks a systems-level map that couples promoter occupancy to time-resolved perturbational responses across multiple classes of transcriptional and epigenetic regulators. Combining promoter occupancy with a rapid transcriptional response to perturbation at a defined cell-cycle time provides convergent evidence for a promoter-linked regulatory association, although it does not establish the underlying molecular mechanism.

Here, we chart a broad regulatory landscape of the tachyzoite division cycle by strategically sampling 21 nuclear-localized factors, comprising 11 ApiAP2 proteins, three Myb-domain proteins, five chromatin regulators, and two additional nuclear proteins: Smad nuclear interacting protein 1 (SNIP1), which contains a forkhead-associated domain, and a previously uncharacterized protein we name AP2-associated factor 1 (AAF1). For each factor, we profiled single-cell transcriptomes after conditional depletion and mapped promoter occupancy by CUT&RUN (cleavage under targets and release using nuclease). We defined a regulatory association as promoter occupancy coupled to a time-matched expression response after depletion, with the response direction distinguishing activating from repressive associations. Using these associations, we assessed the robustness of network structure across a broad range of plausible parameter settings and, to limit dependence on any single analytical choice, retained associations recovered in the majority of networks. This majority-consensus framework yielded a time-resolved, promoter-linked gene regulatory network (GRN) of the tachyzoite cell division cycle.

The resulting GRN links successive division-cycle programs to factors associated with their activation or repression and places poorly characterized genes in functionally interpretable regulatory neighborhoods. Across the profiled factors, promoter occupancy was better predicted by promoter accessibility, promoter architecture, sequence features, and the surrounding factor environment than by a simple factor–motif code. These findings support a predictive opportunity-followed-by-selection model of promoter engagement without defining a causal recruitment hierarchy. Factor-centered analyses recovered known regulatory partnerships and nominated new ones; selected examples were validated through molecular biology and functional experiments.

Together, the breadth of these data and the temporal resolution of the single-cell perturbations provide an integrated view of the *T. gondii* division-cycle regulatory system. They reveal how genes are organized into functional modules, when those modules are deployed, which transcriptional and epigenetic regulators are associated with them, and where combinatorial factor relationships and candidate protein complexes emerge. By integrating timing, occupancy, perturbation, chromatin state, sequence, and molecular validation with analytical tools tailored to periodic biological processes, we transform this multidimensional dataset into an experimentally navigable regulatory map. The interactive companion application makes the map queryable by gene, regulator, community, and time, turning these data into a hypothesis-generation engine for investigating gene function, regulatory relationships, and protein localization. This study therefore provides both a systems-level view of the regulatory programs that accompany growth, daughter construction, and invasion and a practical framework for dissecting compact, evolutionarily divergent eukaryotic regulatory networks.

## 2 Results

### 2.1 Diverse sampling of the lytic cycle transcriptional network

Capturing the breadth of regulation across the lytic cycle requires sampling diverse DNA- and chromatin-binding factor classes and temporal expression patterns. With this goal, we selected 21 factors for conditional depletion guided by four considerations: (1) representation of both transcription factors and chromatin regulators; (2) periodic expression profiles with peaks distributed throughout the lytic cycle (**Fig. 1B** and **Fig. S1**); (3) fitness scores consistent with roles in lytic growth (19 of the 21 selected genes had negative scores in the genome-wide screen) (**Fig. 1C**) [31]. Nineteen of the 21 selected genes had negative fitness scores; and (4) a balance of established regulators and poorly characterized proteins, particularly those with predicted DNA-binding domains. Regarding the third criterion, we made an exception by including periodically expressed ApiAP2 factors that peak during cytokinesis regardless of fitness, so that progression into G1 was sufficiently represented [5,6]. Because ApiAP2 factors are key drivers of lytic-cycle regulation, we included 11 members of this family. The panel also included the less-studied Myb-domain proteins Myb3, DMAP1, and Cdc5, together with two additional nuclear proteins: SNIP1, which contains a forkhead-associated domain, and AA1, a previously uncharacterized protein with a predicted homing-endonuclease-like domain; AP2-associated factor 1 (AAF1) was named based on the ApiAP2 associations uncovered in this study. To extend our sampling to chromatin regulation, we selected five additional factors: MORC and HDAC3, which had been characterized previously [21], and HDAC5, enhancer of polycomb-like protein 1 (EPL1), and anti-silencing factor 1 (ASF1), which had not been functionally examined in *T. gondii*. Although grouped here with the Myb-domain proteins, DMAP1 should also be considered in the context of chromatin regulation because DMAP1 homologs have been linked to epigenetic complexes in other systems [34] (**Fig. S1**). The resulting panel therefore combined distinct predicted molecular functions with expression peaks distributed across the full 6 h tachyzoite division cycle, providing broad coverage of the regulatory space examined.

For each factor, we established parasite lines with indole-3-acetic (IAA) acid conditionally controlled alleles by simultaneously inserting an auxin-inducible degron and a Ty epitope tag (**Fig. 1F** and **Fig. S2**). We performed plaque assays to evaluate parasite growth, which, with few exceptions, aligned well with the reported lytic-cycle fitness scores, although the severity varied among lines (**Fig. 1C** and **Fig. S3**) [31]. Next, we tracked the spatiotemporal expression of each factor at the protein level by immunofluorescence. This confirmed that all factors localized to the nucleus. With the exception of ASF1, factors with strong periodic mRNA profiles showed corresponding changes in protein abundance across the lytic division cycle, whereas factors with less pronounced temporal variation in mRNA abundance generally remained detectable at the protein level throughout the cycle (**Fig. 1D** and **Fig. S4**). Next, we assessed the morphological consequences of depleting each factor for 6 h with IAA, corresponding to approximately one lytic cycle, using DAPI to mark nuclei and IMC3 to mark daughter buds and the cortical cytoskeleton of mature parasites. This interval was chosen to capture early disruptions of budding while limiting secondary defects that could accumulate after longer treatment. We observed defects with varying penetrance in daughter-scaffold assembly and integrity, nuclear organization, and coordination of division among parasites within the same vacuole (**Fig. 1E** and **Fig. S5**; **Table S1**). Phenotypes were scored in three biological replicates, with 100 vacuoles evaluated per replicate. Depletion of DMAP1, MORC, and HDAC3 produced some of the most penetrant phenotypes. Strong defects were not restricted to chromatin-associated factors, as depletion of AP2IX-5 and AP2XI-5 also caused frequent division abnormalities. The AP2IX-5 phenotype is consistent with its established role in initiating the budding transcriptional program [15]. Overall, the diversity of expression patterns and depletion phenotypes confirmed that the selected panel spans regulators acting across distinct stages and cellular processes of the lytic cycle.

To establish the experimental foundation for this time-resolved, promoter-linked GRN, we paired the gene-expression response to depletion of each factor with its genome-wide chromatin-association profile. We measured transcriptional responses by single-cell RNA sequencing (scRNA-seq) and mapped the chromatin association of each Ty-tagged factor by CUT&RUN under permissive conditions (**Fig. 1F**). These datasets provide complementary information: perturbation reveals the transcriptional consequences of factor loss, whereas CUT&RUN identifies promoters associated with that factor. For scRNA-seq, we used a previously optimized 2 h depletion (approximately 30% of the lytic cycle) to capture early transcriptional responses [6]. For this initial analysis, expression responses were considered across the complete cell population rather than resolved by inferred cell-cycle time. Genes that decreased after depletion were classified as activated targets of the factor, whereas genes that increased were classified as repressed targets. We operationally classified a response as direct when a CUT&RUN peak overlapped the annotated transcription start site ±1 kb and as indirect when no promoter occupancy was detected (**Fig. 1G**). These categories identify regulatory associations supported by both promoter occupancy and an expression response without specifying the underlying molecular mechanism. Because HDAC5, AP2VIIa-6, and AP2XII-2 were associated with fewer than 25 promoters, we did not classify them as predominantly activating or repressive in **Fig. 1H**. Their supported regulatory associations were nevertheless retained in the GRN, with each edge still requiring both promoter occupancy and an expression response after factor depletion.

Several chromatin regulators, including HDAC3, MORC, and EPL1, had broad sets of promoter-linked targets. However, target breadth did not track simply with factor class: Cdc5 also had many promoter-linked targets, whereas ASF1 had comparatively few. Most factors were associated with both activated and repressed genes, consistent with combinatorial regulation. Even so, the overall balance was activation-oriented for most factors; MORC, AP2III-2, AP2VIII-7, and AP2XI-4 instead had more repressed than activated promoter-linked targets (**Fig. 1H**). MORC’s repression-oriented profile is consistent with the established silencing function of the MORC– HDAC3 complex, which maintains developmental promoters in a poised state [21].

Together, collective perturbation of these 21 factors revealed broad diversity in mRNA and protein dynamics, growth and cell-division phenotypes, and the direction and breadth of promoter-linked transcriptional responses. This breadth establishes the experimental foundation for the time-resolved GRN and for resolving how individual regulators and their combinations organize gene expression across the tachyzoite lytic cycle.

### 2.2 A periodic single-cell clock resolves factor-specific disruption of the lytic cycle

The *T. gondii* tachyzoite cycle organizes endodyogeny through a just-in-time transcriptional cascade [10]. Individual ApiAP2 factors have been shown to control defined transitions within this program, including AP2IX-5, AP2XI-5, and AP2XII-8 [6,15,17]. To compare these and other regulators in one framework, however, we first needed to place asynchronous wild-type and knockdown parasites on a common cell-cycle scale.

One wild-type RH parent strain library and 21 factor-knockdown libraries were processed using the same quality-control criteria (Methods; **Fig. S6**; **Table S2**). This yielded 211,052 singlets, of which 198,284 could be assigned a cell-cycle time with an estimated error below 0.25 h. Common trajectory methods such as Slingshot and Monocle 2 are designed primarily for linear or branching transitions [35,36]. Lou et al. instead fitted a closed trajectory to cycling tachyzoites and scaled it to a 6 h cycle [6]. We extended this periodic approach beyond a two-dimensional representation by retaining transcriptomic components with strong cyclic signal and projecting each wild-type cell onto the resulting closed reference curve (Methods; **Fig. S7A-B**).

We assigned cell-cycle phases by transferring annotations from an independently characterized *T. gondii* tachyzoite single-cell atlas, in which phase assignments were validated against DNA content [11]. The transferred labels followed the expected progression from G1 through S phase, mitosis, and cytokinesis around the curve. We then mapped position along the curve onto a 6 h scale using cumulative wild-type cell density (Methods). Agreement between the transcriptome-derived ordering and the independently transferred phase labels established a biologically coherent reference clock while preserving continuous transitions between conventional cell-cycle stages (**Fig. 1A** and **Fig. S7C**). The clock therefore captured the order of the lytic program without imposing discrete phase boundaries on the cells.

Each knockdown library was aligned separately to the wild-type reference, and time was inferred from neighboring wild-type cells. To compare all libraries in a common expression space, we used single-cell variational inference (scVI), a probabilistic framework that integrates single-cell transcriptomes while accounting for library-to-library technical variation [37]. Wild-type cells spanned the complete cycle, and cells from all 21 knockdowns occupied the same broad continuum of transcriptional states. This shared coordinate system allowed perturbations to be compared at matched cell-cycle positions rather than conflating regulatory responses with differences in cell-cycle composition.

We next compared each knockdown with wild type across 24 overlapping 1 h windows. **Fig. 2A** summarizes two complementary measures: displacement from the wild-type trajectory at the same inferred time and the numbers of genes whose expression increased or decreased in each window. The profiles revealed factor-specific differences in both the magnitude and timing of transcriptional responses, indicating that depletion of different regulators perturbed distinct portions of the lytic-cycle program. Some perturbations produced concentrated departures over a restricted interval, whereas others affected a broader span of the cycle, revealing when each perturbation was most evident.

**Figure 2.**
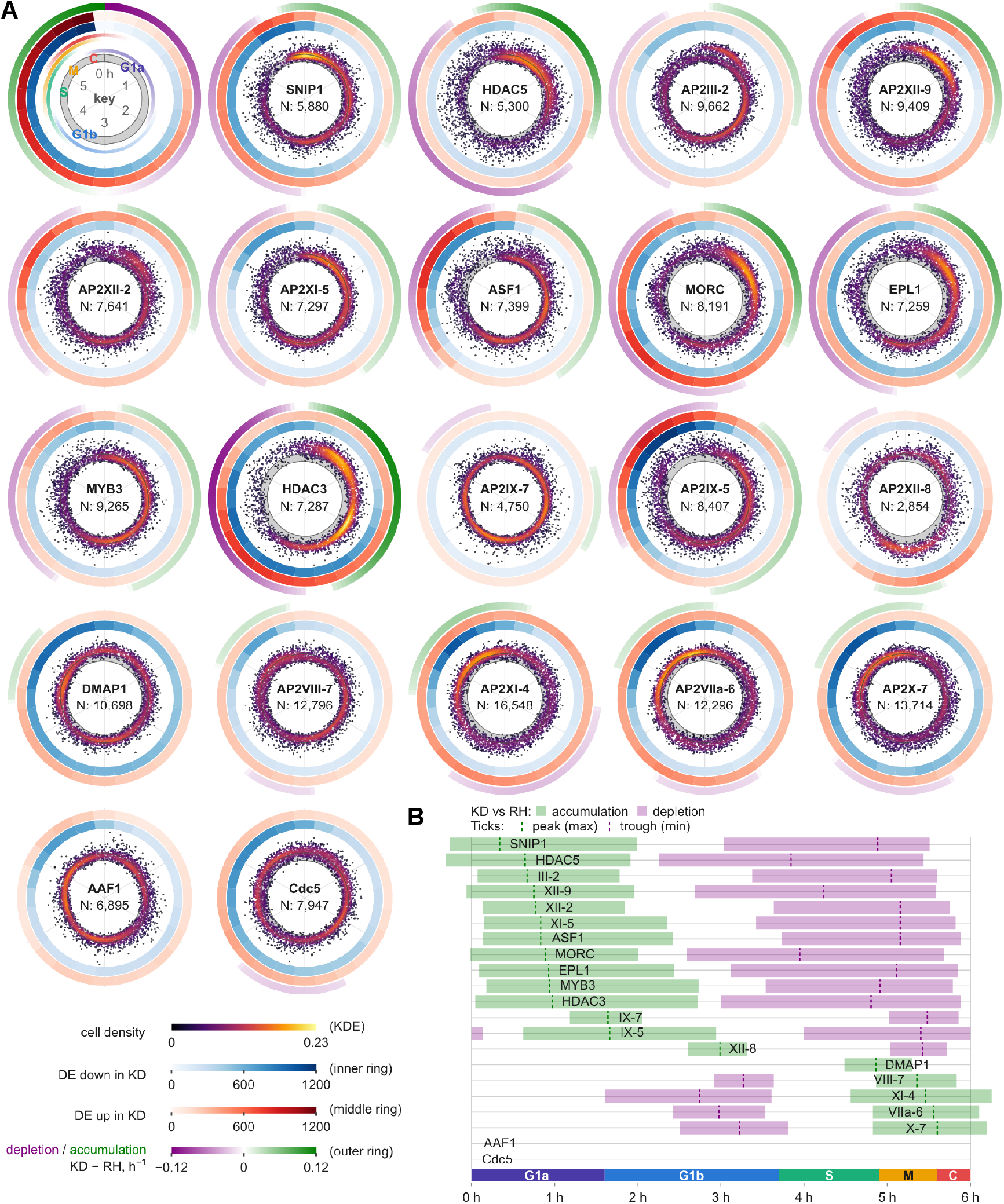
Single-cell RNA sequencing data for wild-type and factor knock-down experiments. **(A)** Transcriptomic deviation from RH along the cell cycle, resulting from knock-down of the indicated factors. Each sub-panel shows a polar-coordinates diagram with cell-cycle time as the angle and average distance to the RH dataset in the integrated shared scVI embedding as the radius. The gray band shows RH’s cell-to-cell spread around its reference trajectory. KD sample points are placed outward proportionally to their transcriptomic deviation from RH and colored by a kernel density estimate. The number of cells in each dataset is shown at the center. The number of differentially expressed genes along the cell cycle (KD vs. WT; per-interval FDR < 0.05, |log2FC| > 1, expressed in > 5% of cells in either condition) is shown as two circular heatmap bands (up- and down-regulated in KD; middle and inner band), one segment per time window (0 h at the top). The outermost band shows the accumulation (green) and depletion (purple) of KD cells along the cycle, as in (B), with color intensity proportional to the difference in cell-time density between KD and RH. The five colored bands inside the key sub-panel (top left) show interpolated wild-type marker-scores along the cycle, so that intervals where consecutive phases overlap appear as two simultaneously-lit bands. **(B)** Accumulation/depletion patterns of the parasite cells along the cell cycle. Three groups can be observed with increased numbers at the G1a/b and M/C phases, while XII-8 displaying accumulation right before S around the 3rd hour of the cycle. Ticks mark each profile’s peak (green) and trough (purple).

Factor depletion also altered the distribution of cells along the cycle (**Fig. 2B**). This analysis considers the complete transcriptomic response and does not separate genes into promoter-linked and indirect targets. Of the 21 knockdowns, 19 produced a detectable cell-density phenotype and could therefore be grouped; AAF1 and Cdc5 showed no detectable accumulation interval and were not assigned to an accumulation pattern. Among the remaining knockdowns, 13 accumulated in G1, five accumulated from late S phase through cytokinesis, and AP2XII-8 showed a distinct accumulation near 3 h, before the inferred onset of S phase. These results define three recurring arrest-like phenotypic patterns: accumulation in G1, accumulation from late S phase through cytokinesis, and the distinct pre-S-phase accumulation caused by AP2XII-8 depletion. The AP2XII-8 pattern agrees with its reported role in cell-cycle progression [6]. The convergence of otherwise distinct perturbations on a small number of accumulation patterns suggests that transcriptional disruptions become evident at a limited set of transition points in the lytic cycle. These profiles identify where cells become overrepresented after depletion but do not by themselves establish arrest or distinguish slower progression from selective loss or altered entry into or exit from a state.

Together, the periodic clock converts an asynchronous perturbation panel into a directly comparable, time-resolved framework. It separates shifts in cell-cycle composition from expression changes within the same inferred interval and provides the temporal coordinate needed to connect promoter occupancy, transcriptional response, and downstream regulatory programs. Rather than producing a uniform disruption, depletion of these regulators altered distinct intervals and transcript sets while frequently converging on recurring cell-cycle accumulation patterns, providing the temporal framework for the promoter-linked GRN that follows.

### 2.3 Integrated promoter occupancy and transcriptional responses define a time-resolved consensus regulatory map

To obtain an initial view of promoter occupancy across the factor panel, we used q < 0.075 as a reference peak threshold, one of the five thresholds subsequently evaluated in the network sweep, rather than a fixed choice for the final network. Across the 21 factors, this reference set contained 42,387 peaks. Assigning peaks to overlapping transcription-start-site ±1-kb promoter windows and retaining the strongest peak for each factor–gene pair yielded 48,615 promoter assignments across 6,888 genes. CUT&RUN signal was concentrated around transcription start sites for most factors, but promoter-association breadth varied by more than three orders of magnitude. Six factors, namely HDAC3, MORC, Cdc5, AP2X-7, AP2III-2, and AP2VIII-7, had broad profiles encompassing 3,519–6,613 assigned genes, eight had intermediate profiles spanning 1,130–2,664 genes, while seven were comparatively sparse, with 3–704 genes. At the extremes, HDAC3 was associated with 6,613 genes, whereas HDAC5 and AP2VIIa-6 were associated with only 3 and 6 genes, respectively (**Fig. 3A**).

**Figure 3.**
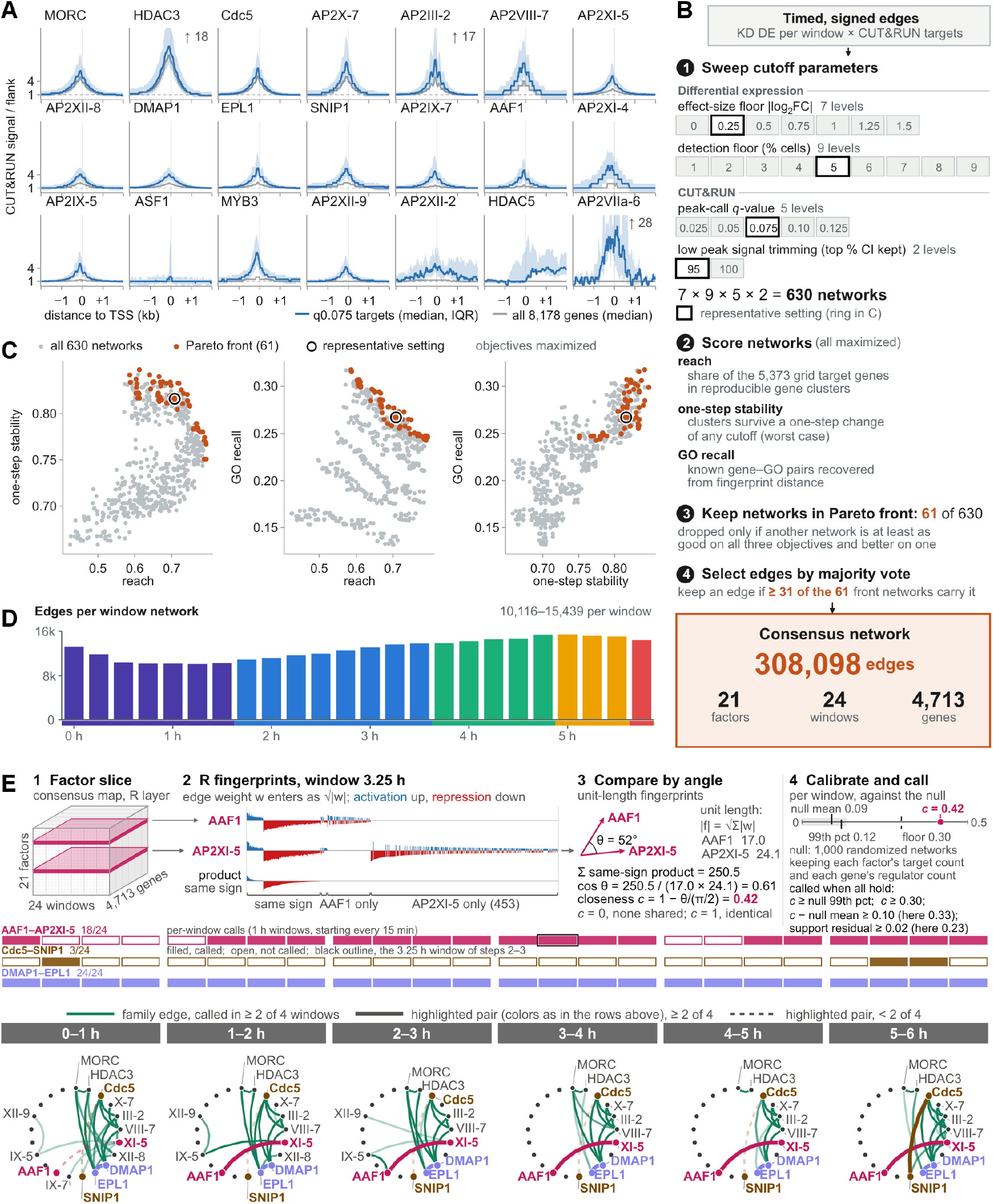
Construction of a stability-selected, time-resolved regulatory map. **(A)** CUT&RUN signal around transcription start sites (TSS ± 2 kb) for the 21 factors: median over each factor’s promoter-assigned target genes (colored line, peaks called at q < 0.075, band = interquartile range across genes), with the median over all 8,178 considered chromosome genes shown in gray behind it for comparison; both normalized to the flank signal (dashed line = 1). **(B)** Network construction and selection. Knock-down DE genes per time window and CUT&RUN promoter targets are combined into timed, signed edges; 630 networks are built from a cutoff parameter grid (one box per level; black outline, the representative setting) over differential expression (effect-size floor |log2FC|, detection floor in % cells) and CUT&RUN (peak-call q-value, low peak signal trimming as top % CI kept), and scored on three objectives; the Pareto front (61 networks) is retained, and the consensus network keeps the edges present in ≥ 31 of the 61 front networks (308,098 edges, 21 factors, 24 windows, 4,713 genes). **(C)** The 630 networks (gray) and the 61-member Pareto front (orange) in pairwise projections of the three objectives: reach, one-step stability, and GO recall (all maximized); black ring, the representative setting. **(D)** Edges per window: number of consensus-network edges in each of the 24 time windows, colored by cell-cycle phase. **(E)** Regulatory fingerprints and regulator-similarity networks (24 overlapping windows, labeled by their centers). 1, A factor’s regulatory fingerprint (R) is its slice of the consensus network: its signed edges to target genes in each window. 2, Activation and repression are kept apart, so only genes regulated in the same direction add to the product of two fingerprints. 3, Fingerprints are compared at unit length by the angle θ between them, counting only same-sign products; the angular closeness c = 1 − θ/(π/2) rescales θ to run from 0 to 1. 4, Each window’s closeness is calibrated against randomized networks (gray band, range of the null), and a window is called only when all four conditions shown hold. The support residual is the closeness left after a LO- ESS trend on the number of shared targets. The calls are a screen, not a multiplicity-corrected test. Rows, per-window calls for the three highlighted pairs. Below, regulator-similarity networks over six 1 h blocks (edge opacity, fraction of the block’s 4 windows called); nodes are placed clockwise by CUT&RUN peak count, and only factors with drawn edges are labeled. Edges indicate similar regulatory programs, not physical interaction. See **Movie S1** for an animated time progression of the network.

We then linked promoter association by each factor to the transcriptional response of the same gene following depletion of that factor in each of 24 overlapping 1-h windows. Genes that decreased after depletion received a positive, activation-associated edge, whereas genes that increased received a negative, repression-associated edge. Edge weights combined occupancy strength with response magnitude, preserving the regulator, target gene, regulatory direction, and cell-cycle time. Constructing these signed networks required choices about peak inclusion, expression-effect magnitude, expression detection, and the treatment of weak occupancy signals.

Because no single combination of these choices uniquely defines the network, we evaluated a systematic parameter grid comprising five peak-call thresholds, seven expression-effect thresholds, nine expression-detection thresholds, and two occupancy-trimming settings, producing 630 candidate networks (**Fig. 3B**). We evaluated each network using three complementary metrics (**Fig. 3C**). **Reach** measured how much of a fixed 5,373-gene universe was represented in gene communities; one-step **stability** measured how little community structure changed between adjacent parameter settings; and GO **recall** measured how well the resulting communities recovered established functional relationships. Together, these objectives distinguished networks that were stable because they were sparse, broad because they were permissive, or biologically coherent but limited in coverage.

Multiobjective optimization retained 61 non-dominated networks that together formed the *Pareto front*. A network lies on this front when no alternative performs at least as well on all three objectives and better on at least one. Rather than selecting a single threshold combination from this front, we retained an edge when it appeared in a majority of the 61 networks, requiring support from at least 31 (**Fig. 3B-C**). The resulting consensus contains 308,098 signed factor–gene– time edges connecting 21 factors to 4,713 genes across 24 windows and retains 87.7% of the 5,373 genes reached by at least one network in the parameter sweep. Network breadth varied across the cycle, from 10,116 edges at 1.25 h to 15,439 at 5.00 h (**Fig. 3D**). Four broadly acting regulators, HDAC3, MORC, AP2X-7, and Cdc5, collectively contributed 145,738 edges (47.3% of the consensus), whereas the remaining factors defined more focused programs. The consensus therefore preserves broad and time-restricted regulatory associations on the same quantitative scale without making the map depend on one arbitrary cutoff.

Because every edge retains its regulator, target, direction, and time, the consensus map can also be read from the perspective of each factor. At each window, we separated the factor’s positive and negative edge weights into activation- and repression-associated channels and normalized the resulting target-gene vector to define its regulatory fingerprint, R (**Fig. 3E**). We compared these fingerprints by normalized angular closeness, which emphasizes whether two regulatory programs point in the same direction rather than the absolute number of edges they contain. Because broadly acting factors have more opportunities to overlap, observed closeness was evaluated against randomized networks that preserved both the number of targets per factor and the number of regulators associated with each gene. This factor-similarity analysis used the regulatory layer alone. In each displayed 1-h block, an edge connects factors that met the predefined similarity rule in at least two of the four constituent windows, and opacity reports the fraction of those windows that passed.

The factor-centered view recovered the established MORC–HDAC3 relationship in all 24 windows and nominated three contrasting shared programs (**Fig. 3E** and **Movie S1**) [21]. DMAP1 and EPL1 were connected in all 24 windows and in all six displayed blocks, nominating a persistent chromatin-regulatory relationship consistent with homologs of both proteins occurring in NuA4/TIP60-family chromatin complexes [38,39]. AAF1 and AP2XI-5 shared a similarly persistent program in 18 of 24 windows, whereas Cdc5 and SNIP1 formed a restricted three-window episode spanning the end and beginning of the circular cycle. These are similarities between regulatory programs, not evidence of physical interaction or recruitment direction. They nevertheless provide specific hypotheses.

Together, integration across the Pareto front produced a parameter-robust, time-resolved, promoter-linked gene regulatory network that can be read by time, factor, or target gene. These complementary views establish the foundation for the factor-centered validation and gene-centered community analyses that follow.

### 2.4 The Cdc5-SNIP1 splicing complex regulates transcription

Our original rationale for including Smad Nuclear Interacting Protein 1 (SNIP1) in our collection was its reported modulation of cMyc TF activity in higher eukaryotes [40,41]. Since *T. gondii* does not encode a cMyc ortholog, we were curious about its role in this system. As shown by the temporal network in **Fig. 3E**, we observed a strong co-regulatory signature between Cdc5 and SNIP1 in the 5-6 h window, which is the late cytokinesis stage of the cell division cycle. This activity trails off to a more modest interaction into G1 at 0-1 h. Since Cdc5, which harbors two Myb domains, was recently reported to be a splicing factor in *T. gondii* [42], its association with SNIP1 initially appeared counterintuitive. However, SNIP1 has another function in the tripartite mRNA retention and splicing (RES) complex, which is conserved from plants to human [43,44], where it acts in splicing and nuclear mRNA retention. The temporal network therefore connected two proteins with independent links to RNA processing and suggested that they might coordinate transcription-associated chromatin occupancy with splicing.

Since SNIP1 function appears to be pluriform, we selected the *T. gondii* Cdc5/SNIP1 regulatory pair for detailed analysis. Cdc5 and SNIP1 displayed tightly paired mRNA-expression profiles that peaked at approximately 4.5 h, immediately before their strongest shared regulatory window (**Fig. 4A**). We next assessed their differential expression and chromatin associated footprints (**Fig. 4B**). This revealed a modest overlap of 75 differentially expressed genes, with Cdc5 and SNIP1 controlling an additional 78 and 256 genes, respectively. At the level of chromatin association, 99.3% of the SNIP1 footprint fell within the broader Cdc5 footprint (**Fig. 4B**), supporting coordinated promoter association. Gene Ontology analysis did not provide a strong signature for the Cdc5-specific or shared transcriptional responses, whereas the SNIP1-specific response was enriched for virulence-related genes involved in budding and apical secretory organelles (**Fig. 4C**). Thus, SNIP1 may regulate part of the transcriptional response independently of Cdc5, but its promoter association occurs almost entirely within the Cdc5-associated landscape.

**Figure 4.**
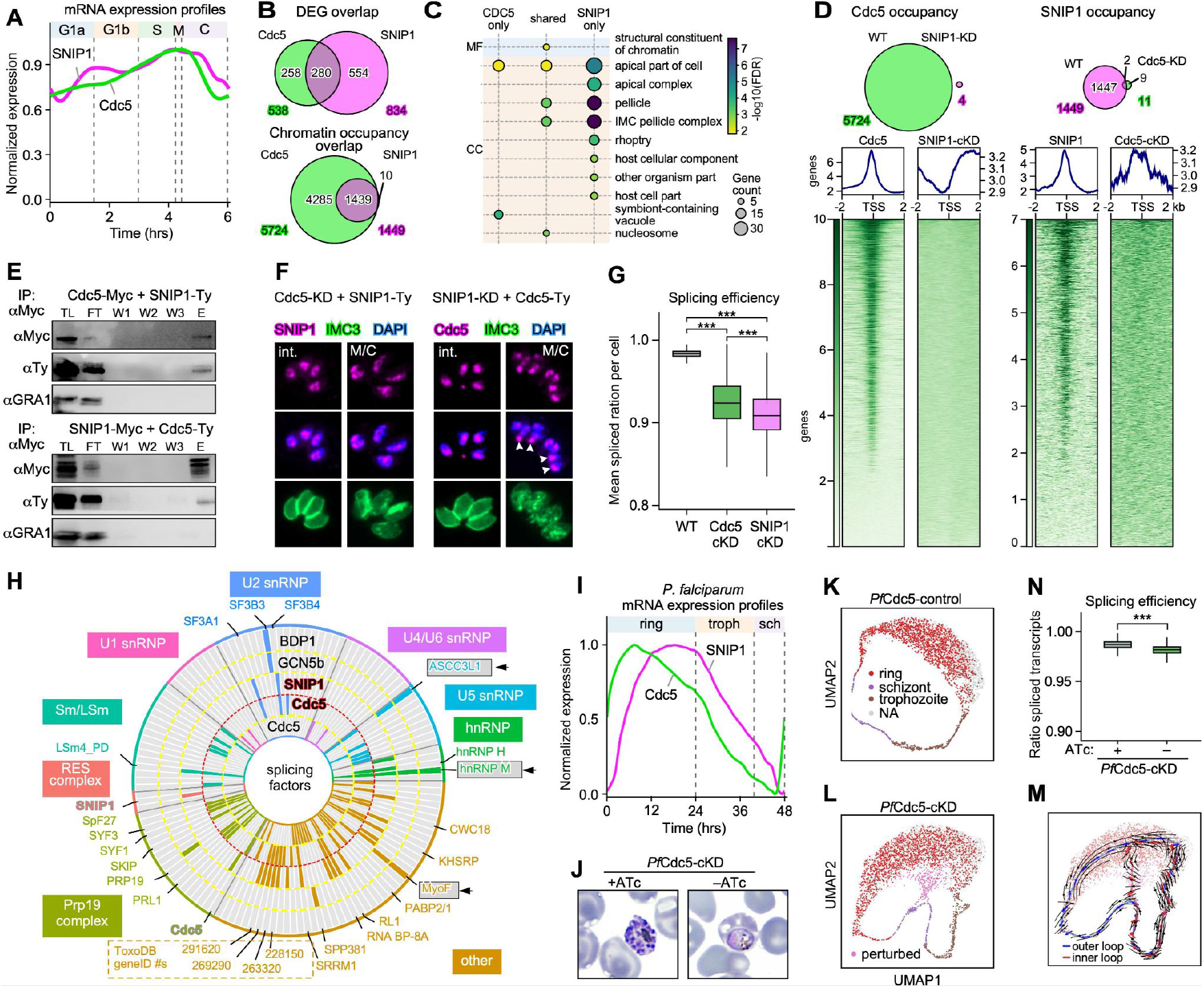
A gene regulatory complex comprising splicing factors Cdc5 and SNIP1. **(A)** Cdc5 and SNIP1 show coincidence expression peaks during the cell division cycle. RNA expression profiles of Cdc5 and SNIP1 across the tachyzoite cell division cycle in wild-type *T. gondii*. Both transcripts peak at the same stage, following the completion of mitosis. **(B)** Shared targets between Cdc5 and SNIP1 depleted parasites; coding and non-coding DEGs in scRNAseq data (top) and chromatin occupancy from CUT&RUN data (bottom). **(C)** GO terms of DEGs. CC: cellular component; MF: molecular function. For GO terms of chromatin occupancy see **Fig. S8G**. **(D)** The Cdc5-SNIP1 complex is required for DNA association; depletion of Cdc5 or SNIP1 results in reduced and altered DNA association of the reciprocal factor. Venn diagram represents peaks assigned to genes detected by CUT&RUN, whereas the bottom panels show a decrease/scatter in association in the region with 2 kb of transcription start sites (TSS). **(E)** Reciprocal co-IP assays of Cdc5 and SNIP1 demonstrate they reside in a complex.TL; total lysate; FT: flow through; W1, 2, 3: washes 1, 2, and 3; E: eluate. **(F)** IFAs of reciprocal depletion of Cdc5 (left) or SNIP1 (right). Depletion of either factor (24 hrs +ATc) causes IMC3 fragmentation and budding defects. Nuclear localization of either factor does not change, but the Cdc5 pattern in SNIP1-depleted mitotic nuclei is suggestive of reduced DNA association (marked by arrowheads). Int.: interphase. **(G)** Depletion of Cdc5 or SNIP1 causes an accumulation of incompletely spliced transcripts. See **Fig. S8** for splicing efficiencies of all factors in this study, and resolution of Cdc5 and SNIP1 reduction in splicing efficiency by intron number, cell cycle stage, or binding in the promoter region. *** *p*<0.0001 **(H)** Pull-down of Cdc5 and SNIP1 (donuts marked by red outlined text) represented in a donut plot reveal overlap in the spliceosome with episomal complexes containing GCN5b and BDP1. The ‘other’ category comprises both RNA binding proteins and other hits in the Cdc5 and SNIP1 pull downs. Published Cdc5 data (inner circle) from [42]; GCN5b pull-down data from [25] and [46]; BDP1 pull-down data from [26]. Factors bridging across spliceosome and epigenetic complexes are marked by arrowheads. See **Fig. S9** for a plot including all names. Boxed ‘others’ are shared between Cdc5 and SNIP1 pull-downs (TGGT1_269290 is annotated as Nup503). **(I)** Expression profiles of the *P. falciparum* Cdc5 (geneID: PF3D7_1033600) and SNIP1 (geneID: PF3D7_1307800) orthologs throughout the erythrocytic cycle show a peak in late ring stage. Schizonts are differentiated in early (e), mid (m) and late (l) stages. **(J) Depletion of PfCdc5 causes cell cycle arrest in *P. falciparum*.** Giemsa-stained thin blood smears of PfCdc5-cKD (pKD-PfCdc5-TetR-DOZI) parasites after 20 h of culture with (+ATc) or without (−ATc) anhydrotetracycline. Parasites were maintained in 20 nM ATc for two intraerythrocytic cycles, after which the culture was split into two equal parts. ATc was removed from one part by washing with fresh medium (−ATc, PfCdc5 depleted), while the other was kept in 20 nM ATc (+ATc, control). Control parasites progressed normally through the cell cycle, whereas PfCdc5-depleted parasites showed an aberrant morphology and arrested without progressing to the segmentation stage, indicating a strong requirement for PfCdc5 in cell cycle progression. **(K)** scRNA-seq UMAPs representing *P. falciparum* Cdc5 replete (top) and depleted (bottom) conditions through TetR aptamer fusion results in depletion of late schizont cells (arrows) and a change in early schizont cell position (arrowheads). See **Fig. S10** for phenotype validation. **(L)** Depletion of *P. falciparum* Cdc5 obstructs the parasite segmentation during early schizogony. **(M)** RNA velocity of PfCdc5-cKD cells, estimated from the ratio of unspliced to spliced transcripts and combined with a guided pseudotime, projected onto the cKD UMAP. The outer loop traces progression through the IDC stages and the inner loop leads into the perturbed cluster. **(N)** Depletion of *P. falciparum* Cdc5 reduces splicing efficiency. See **Fig. S10G-H** for resolution by number of introns and cell cycle stage. *** *p*<0.001.

To directly test this co-dependent chromatin-association model, we established conditional-depletion lines in which the reciprocal partner was endogenously tagged with a 10xMyc epitope tag (**Fig. S2K-N**). Following 2 h of depletion of one factor, we probed chromatin association of the reciprocal factor by CUT&RUN. Depletion of either factor almost completely disrupted the chromatin association of the other, indicating that stable promoter association requires both Cdc5 and SNIP1 (**Fig. 4D**). Because Cdc5 occupied a broader promoter set than SNIP1 in the unperturbed condition (**Fig. 4B**), SNIP1 may help establish or stabilize Cdc5 occupancy and then dissociate from (a subset of) binding sites. This model predicted an association between Cdc5 and SNIP1, which was validated by reciprocal co-immunoprecipitation from parasite lysates (**Fig. 4E**). We further evaluated their nuclear-localization patterns after reciprocal depletion by immunofluorescence. Loss of SNIP1 altered the distribution of Cdc5 relative to chromosomal DNA, with Cdc5 signal visible at the trailing end of mitotic nuclei where DAPI staining was depleted (**Fig. 4F**, arrowheads; see **Fig. S4** for SNIP1 and Cdc5 localization under unperturbed conditions). This imaging result is consistent with reduced Cdc5 association with chromatin after SNIP1 depletion. Together, these experiments support a Cdc5-SNIP1 complex whose stable chromatin association depends on both partners.

We also assessed whether Cdc5 and SNIP1 affect RNA-splicing efficiency by comparing the abundance of spliced and incompletely spliced transcripts. Depletion of either factor led to an approximately 7% reduction in splicing efficiency (**Fig. 4G**), with the severity of the defect increasing with intron number (**Fig. S8B**). Notably, none of the other factor depletions in our set caused a significant splicing defect, indicating that Cdc5 and SNIP1 act together in RNA processing (**Fig. S8A**). Because Cdc5 and SNIP1 are periodically expressed and their strongest shared regulatory activity occurred at 5-6 h, we asked whether splicing efficiency was affected in a stage-dependent manner. We did not detect a corresponding periodic effect (**Fig. S8C**), indicating that their function in splicing extends beyond the narrower temporal window of their strongest shared transcriptional activity. Promoter association by Cdc5 and SNIP1 also increased with gene intron number (**Fig. S8F**), linking the genes engaged by the pair to those with greater splicing demand. Together, these analyses establish a broad splicing contribution that accompanies their more temporally restricted regulatory relationship.

Given the established roles of Cdc5 and SNIP1 in pre-mRNA splicing, we expected their binding to occur preferentially at intron-containing genes. However, intron-distribution analysis showed that a substantial proportion of their CUT&RUN targets were intron-deprived. Only mRNA (protein-coding) genes were included in intron-distribution analyses. Intron-deprived genes accounted for 15.8% of Cdc5-unique targets (667 of 4,214), 18.6% of shared targets (254 of 1,364) and 11.1% of SNIP1-unique targets (1 of 9) (**Fig. S8E**; **Table S3**). Gene ontology analysis of the shared CUT&RUN targets of Cdc5 and SNIP1 showed enrichment for genes involved in budding and in apical secretory organelles. This is consistent with the phenotype observed in cells depleted of these factors (**Fig. S8G**).

We then examined the intron distribution of protein coding DEGs identified from scRNA-seq data. Intron-less genes made up 37.5% of Cdc5-unique DEGs (24 of 64), 17.2% of SNIP1-unique DEGs (42 of 244) and 12.3% of shared DEGs (8 of 65). Intron-rich genes (>4 introns) made up 34.4%, 42.2% and 47.7% of these groups, respectively (**Fig. S8D**). Enrichment analysis of the direct functional targets also identified preferred associations with both intron-rich and intron-deprived genes (**Fig. S8F**). These findings indicate that Cdc5 and SNIP1 associate with genes irrespective of their intron content, suggesting that their regulatory functions may extend beyond mRNA splicing to include roles in transcriptional regulation.

To define the molecular context of these transcriptional and splicing phenotypes, we performed pull-downs with both factors from total parasite lysates. Cdc5 recovered many components of the NTC/Prp19C spliceosomal subcomplex in which it resides [45], together with components of additional spliceosomal subcomplexes (**Fig. 4H**; **Fig. S9**; **Table S4**). Our data overlapped with recently published Cdc5 pull-down data [42], shown in the inner circle of **Fig. 4H**. However, SNIP1 was not detected by mass spectrometry in the Cdc5 pull-down, and the reciprocal SNIP1 pull-down likewise did not recover Cdc5. The two datasets nevertheless overlapped in components of the U2 snRNP and hnRNP subcomplexes (**Fig. 4H**). Both baits also recovered proteins with RNA-binding domains (contained in ‘other’ category) as well as a set of hypothetical proteins that by hyperLOPIT compartmental proteomics were detected in the nucleus. Surprisingly, SNIP1 does not reveal the other two members of the RES complex, which are encoded in the *T. gondii* genome (Bud13: TGGT1_264130; Snu17: TGGT1_263595), suggesting that this interaction may be transient or that SNIP1 participates in a distinct complex.

We also detected several proteins previously reported in pull-down experiments using the *T. gondii* chromatin-associated factors GCN5b [25,46] and BDP1 [26] as baits, including ASCC3L1, hnRNP-M, and MyoF (**Fig. 4H**). Those studies noted the otherwise puzzling presence of spliceosomal components. Their recurrence in the Cdc5 and SNIP1 pull-down data suggests that this overlap is not coincidental and identifies a potential molecular connection between spliceosomal and chromatin-associated regulatory proteins.

To test the dual transcription-associated and splicing function of Cdc5 in another apicomplexan lineage, we turned to *P. falciparum*, which has substantially fewer introns than *T. gondii*. Intron-deprived genes make up 19.3% of protein-coding genes in *T. gondii ME49* (1,568 of 8,140) while up to 44.5% in *P. falciparum 3D7* (2,369 of 5,318). The Cdc5 ortholog in *P. falciparum* is known as CEF1 (PF3D7_1033600), although we use PfCdc5 here for consistency [47]. PfCdc5 has a TN-seq mutagenesis index score (MIS) of 0.12 in *P. falciparum* [48], indicating low tolerance for disruption and supporting an essential role during the intraerythrocytic developmental cycle (IDC). Its *P. knowlesi* ortholog (PKNH_0618400, pre-mRNA-splicing factor CEF1, putative) is similarly intolerant of disruption in a genome-wide transposon mutagenesis screen, with an MIS of 0.176 [49]. Both *Pf*Cdc5 and the *P. falciparum* ortholog of SNIP1 (PF3D7_1307800) peak in expression at the ring stage and are least expressed in schizonts (**Fig. 4I**). To study PfCdc5 function, a TetR-DOZI mutant was generated for PfCdc5 (**Fig. S10A-B**). To establish the ATc dose required for knockdown and the number of cycles needed for the phenotype to manifest, the ATc concentration was titered out (**Fig. S10C**). 20 nM represents the lowest ATc concentration that sustains normal growth, and consequently the lowest concentration from which ATc withdrawal generates a cKD within a single cell cycle. By Giemsa staining, this resulted in a late trophozoite/early schizont arrested state where merozoite segmentation was never observed (**Fig. 4J**).

scRNAseq of the PfCdc5 control shows the complete IDC UMAP with rings, trophozoites and schizonts (**Fig. 4K**), whereas the cKD shows a unique cell population alongside the IDC stages, here labelled as “perturbed cells” (**Fig. 4K-M** and **Fig. S10D-F**). Unlike its *T. gondii* counterpart, the perturbed cells in PfCdc5-cKD form a new cluster, resulting from arrested parasites at early schizogony. Comparing this cluster to rings and to schizonts within the cell cycle gave 23 and 73 DEGs, respectively, within the cKD culture. To determine how these perturbed cells arise, we estimated RNA velocity from the ratio of unspliced to spliced transcripts in each cell (velocity package). We combined it with a guided pseudotime and projected the result onto the cKD UMAP (**Fig. 4M**). The velocity streams followed two routes. The outer loop traced progression through the IDC stages, and the inner loop led into the perturbed cluster. This indicates that perturbed cells originate from parasites progressing through the cycle rather than forming a separate population, and the direction of RNA maturation agrees with this pseudotime path. Consistent with the effect of Cdc5 depletion on transcript processing, the overall ratio of spliced transcripts was significantly lower in the cKD than in the control (**Fig. 4N**; p < 0.001). Of these, 7 out of 23 and 33 out of 73 genes were intron-deprived in perturbed versus rings and perturbed versus schizonts, respectively (**Table S5** and **Table S6**). This indicates a role for PfCdc5 in the transcriptional regulation of intron-deprived genes, although further investigation is required to validate this hypothesis. GO enrichment of perturbed cells versus rings showed enrichment of nucleosome assembly among the upregulated genes. For perturbed cells versus schizonts, the downregulated genes were enriched for secretory organelles and the upregulated genes for protein trafficking machinery, consistent with the survival of the cKD within the RBCs (**Table S5** and **Table S6**).

Of the 505 protein-coding *T. gondii* DEGs, 412 were intron-containing (126 up, 286 down) and 93 intron-deprived (49 up, 44 down). Intron-deprived genes were enriched among up-regulated DEGs (28.0% vs 15.5% of detected genes; p = 1.2 × 10⁻⁵) but not among down-regulated DEGs (13.3%; p = 0.88). Only 7 of the 49 up-regulated intron-deprived DEGs (14%) had a *P. falciparum* ortholog, compared with 23 of 44 down-regulated intron-deprived (52%) and 222 of 412 intron-containing DEGs (54%). Compared with *P. falciparum* Cdc5-perturbed vs schizont DEGs, ortholog groups down-regulated in both species were enriched (70 observed vs 45.8 expected; p = 1.9 × 10⁻⁶; 25 one-to-one), whereas those up-regulated in both were not (11 vs 7.9; p = 0.15). The 25 one-to-one groups include Tg histone H4, ASF1 family protein, thymidylate kinase, RON6, APR2 and AAP7 (the last two with uncharacterized Pf orthologs), proteasome subunit beta type-3 and 26S protease regulatory subunit 4. Intron status was often not shared: 52 of the 113 intron-containing Tg DEGs with a Pf schizont-DEG partner had at least one intron-deprived partner. Pairs intron-deprived in both species and regulated in the same direction were limited to Tg histones H4, H2Ba, H2Bb, H3 and H2A1 and an EF hand domain-containing protein (all down), and an RNA recognition motif-containing protein and exosome component RRP43 (both up).

The cKD shows a significant reduction in splicing efficiency as compared to the control parasite culture (**Fig. 4N**). Examining splicing efficiency across genes indicated that the splicing efficiency is inversely proportional to the number of introns, up to an intron number greater than 4, after which there was no significant difference compared to the control (**Fig. S10G**). Splicing efficiency also appears to be reduced in the cKD across the IDC stages of the parasite (**Fig. S10H**). Together, these findings indicate that PfCdc5 functions in both splicing and transcriptional regulation, as observed for its *T. gondii* counterpart, suggesting that Cdc5 and its dual function are evolutionarily conserved.

Taken together, our broad sampling of putative DNA-binding proteins led to the insight that Cdc5 and SNIP1 co-regulate a set of genes late in the cell-division cycle. In line with documented roles for these factors, both also contribute to mRNA splicing. Beyond splicing, Cdc5 and SNIP1 associate with chromatin in a co-dependent manner, although the current data do not establish whether this dependence is direct or mediated through additional proteins. The splicing function of SNIP1 is conserved from Apicomplexa to plants, yeast, and mammals, whereas its transcription-associated role through c-Myc in mammals appears to be a lineage-specific adaptation. The intersection of Cdc5 and SNIP1 with both spliceosomal and chromatin-associated proteins provides a route by which transcription and RNA processing can be coordinated.

### 2.5 Regulatory fingerprints organize the lytic-cycle cascade into functional gene communities

To compare target genes in a common framework, we read the consensus factor-by-time-by-gene regulatory tensor by gene across the complete cycle (**Fig. 5A**). A gene slice preserves which factors are associated with that gene, in which direction, and when. We separated activation- and repression-associated edges to define each gene’s regulatory fingerprint (**R**), then considered this regulatory view alongside its matched wild-type expression (**E**) and promoter-accessibility (**A**) trajectories. Each view was converted separately to a normalized angular distance, and the three distances were averaged to define the primary fused R/E/A distance. R alone and the combined R/E distance were retained as sensitivity analyses. Angular distance emphasizes shared patterns rather than the overall signal magnitude that can dominate Euclidean distance. Thus, R identifies genes with similar regulatory inputs, whereas E and A ask whether those relationships are reinforced by when a gene is expressed and when its promoter is accessible.

**Figure 5.**
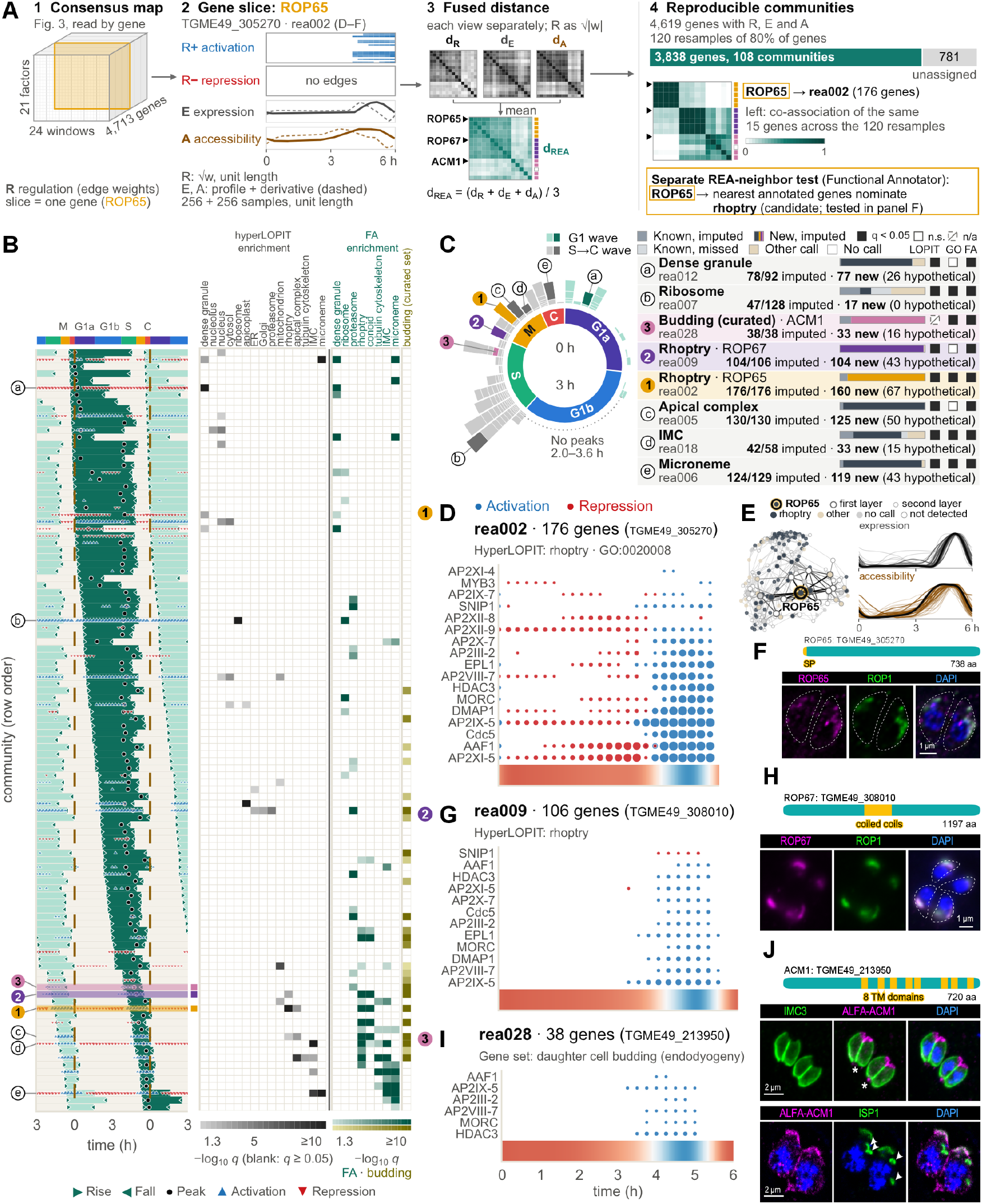
Regulatory fingerprints resolve a timed transcriptional cascade and nominate gene functions. **(A)** Gene fingerprints and gene communities. 1, The consensus map of Fig. 3 read by gene: each gene’s fingerprint is its slice of regulatory edges (R). 2, Each gene is also described by its wild-type expression (E) and accessibility (A) profiles, shown for ROP65. 3, Each view gives a separate distance between genes (angle over its maximum: π/2 for R, π for E and A), and the three are averaged into the fused distance d_REA_. Shown for 15 genes: the tracked proteins of panels F, H, and J, each followed by its four most frequently co-clustered genes (ties broken by smaller d_REA_); each heatmap is scaled separately, darker = closer; side bars, community of each gene (colors as in B–J; gray, another community; open, unassigned). 4, Ensemble clustering over gene resamples commits genes to reproducible communities and leaves the rest unassigned; heatmap, the fraction of resamples drawing both genes in which they cluster together, same genes and order. **(B)** Expression timing of the 108 communities (rows) over two repeated 6-h cycles: dark block, on-interval (half-amplitude rise to fall), repeated in light every 6 h; rows ordered by rise time, starting with the first community peaking after 0 h. Activation and repression markers, ≥ 70% of significant regulators (fdr < 10^−10^) agree (size, best fdr). Right, enrichment of hyperLOPIT compartments and of Functional Annotator (FA) calls (members whose top-3 calls include the term, against the 4,619 annotated genes; one-sided hypergeometric test, BH over all 108 × 9 cells) for the eight compartments with a matching GO cellular-component term (conoid, a part of the apical complex, for the apical column) and, last, for the curated daughter-cell budding set. The calls come from the same network that built the communities, so this shows where they concentrate, not independent validation. **(C)** Expression peaks (left; one block per community in its 0.125 h peak bin, length ∝ genes; teal = G1 wave, 20 communities, 608 genes; gray = S→C wave, 88 communities, 3,230 genes; dark or link color = badged) and the hyperLOPIT make-up of the badged communities (right; calls at probability ≥ 0.99; fold over all 4,619 genes); lettered and numbered badges mark the communities in (B) and (D, G, I). (D, G, and I) Directional regulatory profiles for rea002, rea009, and rea028. At each time window, one-sided Fisher tests identify factors whose activating (blue) or repressive (red) consensus edges are enriched in the community (BH q < 0.05; circle size, −log10 q). The heat strip below each plot shows mean community expression on the shared time axis. Headings give community size and the top displayed annotation from GO, hyperLOPIT, or the curated daughter-budding gene set. **(E)** (right) expression and accessibility profiles for community rea002, and (left) the community genes and their 4 closest neighbors given by their fingerprint distances. **(F)** Ty-tagged ROP65 (TGME49_305270), with a predicted signal peptide, co-stained with ROP1. **(H)** Ty-tagged ROP67 (TGME49_308010), with predicted coiled-coil regions, co-stained with ROP1. **(J)** ALFA-tagged ACM1 (TGME49_213950), with eight predicted transmembrane domains, costained with IMC3 (upper row) or ISP1 (lower row). DAPI marks DNA. SP, signal peptide; TM, transmembrane domain.

We used this gene-centered geometry to identify groups that persisted under resampling [50]. Of the 4,713 target genes in the consensus map, 4,619 had complete R, E, and A profiles and entered the analysis. Across 120 resamples containing 80% of these genes, 3,838 genes were assigned to 108 reproducible communities (**Fig. 5A-B**). Community sizes ranged from 10 to 181 genes, with a median of 19; 781 genes remained unassigned. This organization reduces a dense factor-gene-time network to temporally coordinated modules that can be examined by timing, function, and regulatory input.

The communities formed a coordinated transcriptional cascade across the 6-h cycle (**Fig. 5B-C**). Twenty communities containing 608 genes peaked during the first 2 h, followed by a gap in community peak times from 2 to 3.5 h; the remaining 88 communities, containing 3,230 genes, peaked from late G1 through S phase, mitosis, and cytokinesis. The first wave was dominated by growth and biosynthetic programs, whereas the second assembled daughter-cell and invasion machinery, consistent with successive waves of lytic-cycle transcription and the modular construction of apicomplexan daughters [1,10]. The map therefore resolves the just-in-time program into an ordered transition from early-cycle maintenance to late-cycle morphogenesis and invasion. Its resolution is greatest across S phase, mitosis, and cytokinesis, when DNA replication, daughter budding, and apical-organelle biogenesis proceed concurrently. Rather than constituting one broad late program, these processes are partitioned among successive modules with distinct regulatory inputs.

Co-regulated genes also provide functional context by association, an important advantage in *T. gondii*, where many lineage-specific proteins remain poorly annotated. Separately from community detection, we developed a ‘Functional Annotator’ that used the related R/E/A neighborhood to ask which annotated gene sets lie unusually close to a query and whether that association is unusually specific compared with genes having similar regulatory support (**Fig. 5A**; Section “Community identity: Functional Annotator”). We integrated these predictions with GO terms and high-confidence hyperLOPIT assignments to interpret the communities (**Fig. 5B-C**). The resulting modules were not simple organelle bins: individual communities contained genes assigned to several cellular functions, whereas genes associated with the same organelle were distributed across multiple communities. This organization argues against a simple one-organelle/one-community model. Instead, different components of an organelle can be embedded in distinct temporal and regulatory programs, while one community can coordinate several cellular processes that must proceed together during endodyogeny. To examine this organization in a concrete biological setting, we focused on two rhoptry-associated communities and one daughter-budding community.

We next tested, at each cell-cycle window, whether activation- or repression-associated edges from each factor were overrepresented among the genes in a community. These directional tests connected community dynamics to distinct combinations of promoter-linked regulatory associations (**Fig. 5D**, **Fig. 5G**, **Fig. 5I**). In the 176-gene ROP65-containing rhoptry program, rea002, 17 of 21 factors were enriched in at least one window. Repressive associations predominated before the program rose, whereas activating associations began at 3.5 h and expanded across a broad factor cohort as community expression increased and peaked (**Fig. 5D**). The expression and accessibility trajectories and local regulatory neighborhood of this program are shown in **Fig. 5E**. The 106-gene TGME49_308010 rhoptry program, rea009, followed a different route to a similar late-cycle output: 11 factors contributed activating associations from 3.25 to 5.5 h, whereas only two contributed sparse repressive associations (**Fig. 5G**). The 38-gene ACM1 daughter-budding program, rea028, received activating associations from six factors between 3.25 and 5.0 h without a corresponding repressive phase (**Fig. 5I**). Thus, late-cycle programs are not governed by one universal switch. The two rhoptry programs reach a related organellar outcome through different balances of repression and activation, whereas the budding program is dominated by coordinated activating associations.

These communities also guided experimental follow-up through three complementary selection routes (**Fig. 5C-J**). ROP65 was prioritized from rea002 because both its regulatory neighborhood and its high-confidence hyperLOPIT assignment linked it to rhoptry biology. Endogenous tagging confirmed its localization with the rhoptry marker ROP1 (**Fig. 5F**), consistent with the recent independent annotation of this protein as ROP65 [51]. The rhoptry-associated rea009 community prioritized TGME49_308010, for which no functional data were available. The tagged protein showed an apical punctate pattern in parasites co-stained for ROP1 (**Fig. 5H**), and we named it ROP67. We used a different query to investigate division-stage biology that is underrepresented in existing system-wide localization datasets. Notably, hyperLOPIT data are derived from non-dividing extracellular parasites [32], whereas proximity-labeling surveys cannot probe inside membranes [52–54]. Hence, we first identified communities containing a curated set of daughter-budding genes (**Table S7**) and then prioritized poorly or uncharacterized proteins with predicted transmembrane domains and strong negative fitness scores. This route nominated TGME49_213950, which has eight predicted transmembrane domains and a fitness score of -3.91. The tagged protein localized to the apical cap of mature mother parasites but was not detected in budding daughters (**Fig. 5J**). We therefore named it Apical Cap Membrane protein 1 (ACM1). These examples show how the map can recover established organelle biology while also nominating proteins missed by static or annotation-dependent approaches.

Together, gene-centered regulatory fingerprints transform the consensus network into three connected outputs: a temporal atlas of cellular programs, functional hypotheses for uncharacterized genes, and a window-resolved map of candidate regulatory inputs. The communities reveal that a common cellular destination, including rhoptry biogenesis, can be reached through distinct regulatory trajectories, while individual modules coordinate genes spanning several cellular processes. By connecting when a program is deployed, which promoter-linked factors accompany its rise or repression, and which uncharacterized genes share its neighborhood, the map turns the lytic-cycle cascade into a set of experimentally addressable biological modules.

### 2.6 Regulatory opportunity and factor combinations shape ApiAP2 promoter engagement

ApiAP2 DNA-binding domains recognize defined sequences, and protein-binding microarrays established sequence preferences for this family in *Plasmodium* spp. [13,14]. Individual *T. gondii* ApiAP2s likewise engage biologically important promoter motifs, while studies of AP2XI-5 and AP2X-5 show that factor partnerships can shape promoter occupancy [17,18]. Yet a motif match alone cannot explain why one promoter is occupied while another promoter carrying the same sequence is not, or why several factors repeatedly converge on the same promoter. Across plants and animals, chromatin state, co-bound factors, and partner-dependent motif syntax help resolve this same gap between biochemical sequence recognition and genomic occupancy [28,29,55,56]. We therefore asked how sequence, promoter state, and the local factor environment combine to distinguish the promoters engaged by each factor.

We first built a systematic motif map from CUT&RUN peak sequences using two complementary discovery arms (**Fig. 6A** and **Fig. S11A**). One arm followed the conventional sequence-centered approach: on seven prespecified discovery chromosomes, each factor’s peaks at functional target promoters were compared with dinucleotide-preserving shuffles of the same sequences, yielding 265 position-weight matrices (PWMs). A second arm used all 14 chromosomes to compare summit-centered windows with promoter windows matched for accessibility, GC content, co-binding burden, and position, retaining 17 high confidence PWMs. Before merging similar PWMs, we added four prespecified motif anchors so that an established or hypothesis-driven sequence could enter the same promoter-wide predictive test even when its exact reference sequence was not recovered de novo. The anchors were the experimentally validated AP2XI-5 motif GCTAGC [17]; TGCATGCA, an instance of the *T. gondii* ribosomal-protein promoter element TRP-2 that was subsequently linked to AP2XII-8 [6,57]; an AP2XI-4-associated GT/CA repeat nominated by our preliminary analysis (GTGTGTGT); and TATAWAWR as a general core-promoter comparator. The resulting 286 candidate PWMs were grouped by sequence similarity, and representatives were selected using occupancy statistics from all 14 chromosomes, yielding a shared set of 72 non-redundant motifs (**Fig. S11B**). Sixteen of 1,080 factor–motif combinations showed summit association: they were enriched and centered under a factor’s CUT&RUN summits, and 12 reproduced in a separate chromosome-half check. Thus, motif discovery defined both a common sequence representation for predictive modeling and factor–motif associations that could be distinguished from generic promoter composition.

**Figure 6.**
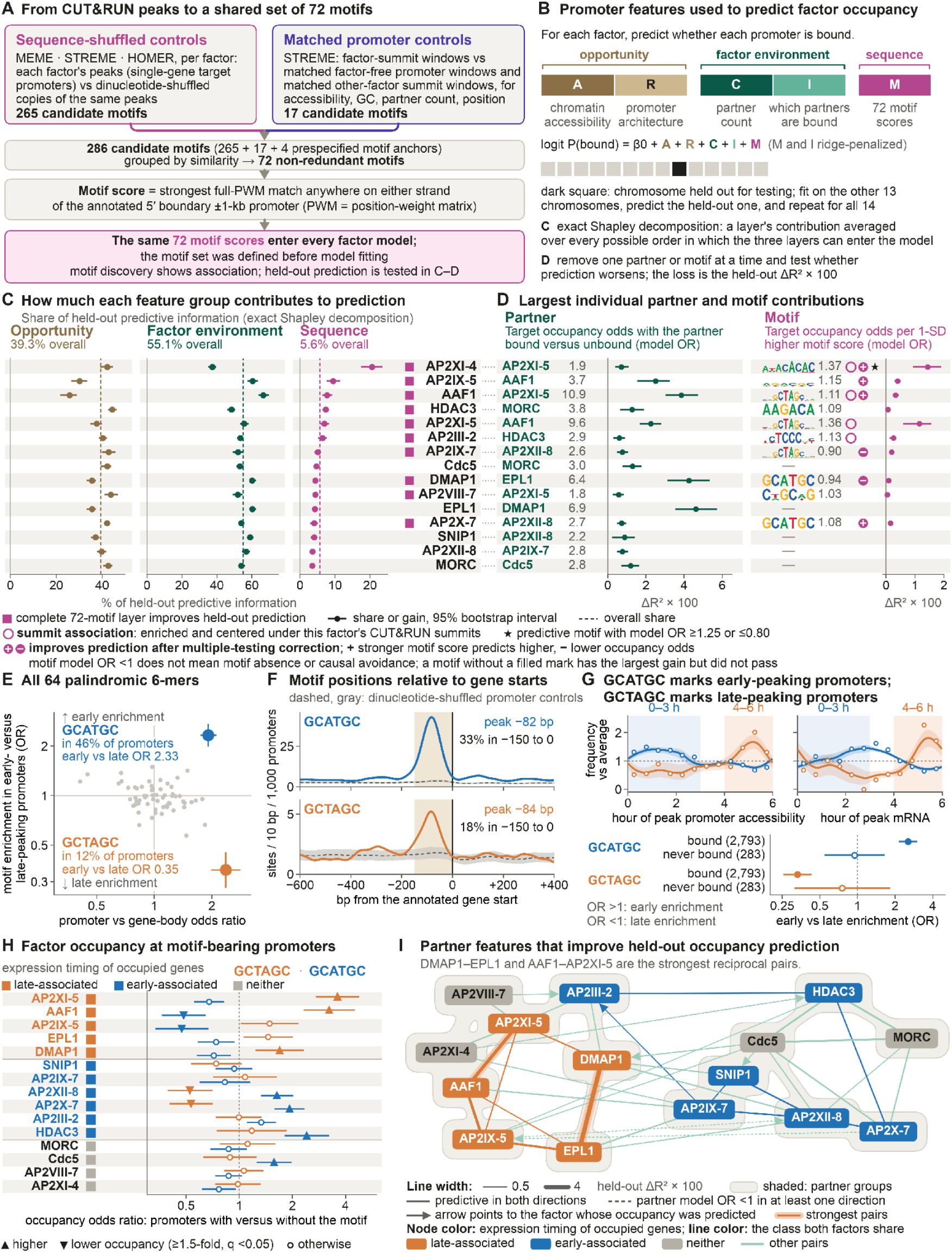
Promoter opportunity, the factor environment, and sequence predict promoter occupancy. **(A)** Two complementary analyses generated candidate position-weight matrices (PWMs) from CUT&RUN peak sequences. The sequence-shuffled arm compared full peak intervals at singlegene promoters of functional targets with dinucleotide-preserving shuffles of the same sequences, yielding 265 candidate motifs from seven prespecified discovery chromosomes. The matched-promoter arm compared 101-bp factor-summit windows with factor-free promoter windows and other-factor summit windows matched for accessibility, GC content, partner count, and promoter position, retaining 17 candidate motifs from all 14 chromosomes. Adding four pre-specified motif anchors (GCTAGC, TGCATGCA, TATAWAWR, and GTGTGTGT)produced 286 candidate PWMs. Similarity grouping and removal of redundancy yielded 72 motifs. Each promoter received a continuous score for each motif, defined as the strongest full-PWM match on either strand of the annotated 5′ boundary ±1-kb promoter. The same 72 motif scores entered every factor model. Because motif discovery and representative selection used chromosome-wide information overall, chromosome holdout tests predictive performance, not motif discovery. **(B)** Ridge-logistic models predicted consensus CUT&RUN occupancy separately for 15 factors across 8,116 promoters. The predictors comprised promoter opportunity—chromatin accessibility (**A**) and promoter architecture (**R**)—the factor environment—other-factor count (**C**) and the identities of the other bound factors beyond count (**I**)—and sequence (**M**), represented by the 72 continuous motif scores. **M** and **I** were ridge-penalized. Each model was fitted on 13 chromosomes and evaluated on the omitted chromosome, repeating the procedure until all 14 chromosomes had served as the test set. All transformations and penalty choices were learned without the held-out chromosome. **(C)** Exact Shapley decomposition allocated the complete model’s held-out predictive information among promoter opportunity, factor environment, and sequence by averaging the information added by each layer over every possible order of inclusion. Points show factor-specific shares; horizontal bars show descriptive 95% chromosome-bootstrap intervals. Dashed lines show the across-factor shares: 39.3% for promoter opportunity, 55.1% for factor environment, and 5.6% for sequence. Magenta squares mark the 10 factors for which the complete 72-motif layer improved held-out prediction under the exploratory layer-level BH *q* ≤0.10 rule. Factor names at the right edge align with the rows in (D), and dotted guides connect each factor to its leading partner and motif. Thin white bands separate promoter opportunity, factor environment, and sequence. Shapley allocation measures predictive information, not causal effects. **(D)** Individual partner and motif contributions were evaluated by removing one feature from the complete model and refitting under the same modeling rule. The horizontal axis shows the resulting loss in held-out McFadden R², multiplied by 100; McFadden R² measures held-out log-likelihood improvement over a prevalence-only prediction. Bars show descriptive 95% chromosome-bootstrap intervals. The partner column shows, for each target, the partner with the largest held-out gain among those that improved prediction after multiple-testing correction and met the model-OR criterion. Its partner model odds ratio (OR) compares modeled target-occupancy odds when that partner is bound versus unbound, conditional on the remaining features. For each factor whose complete sequence layer passed the layer-level test, the motif column shows the motif with the largest held-out gain, whether or not that motif passed the individual-feature test. Its motif model OR is the multiplicative change in modeled occupancy odds per one fitting-set standard-deviation increase in motif score, conditional on the remaining features. A thin white band separates the partner and motif subpanels. Open magenta rings mark summit associations: motifs independently enriched and centered under that factor’s CUT&RUN summits and reproduced across chromosome halves. Filled magenta marks denote motifs that improved held-out prediction after multiple-testing correction; white **+** or **−** indicates a motif model OR above or below one. Stars additionally mark predictive motifs with model OR ≥1.25 or ≤0.80. A motif without a filled mark is the largest-gain motif for that row but did not pass the individual predictive test. A motif model OR below one does not imply motif absence or causal avoidance. **(E)** All 64 reverse-complement-palindromic 6-mers were evaluated in 4,288 single-gene promoter cores. The horizontal axis is the GC-stratified Mantel–Haenszel OR comparing motif occurrence in these promoters with equal-length gene-body windows from 3,892 genes. The vertical axis is the accessibility- and GC-stratified OR comparing promoters with early (0–3 h; *n* = 1,353) versus late (4–6 h; *n* = 1,723) accessibility peaks. OR >1 indicates early enrichment and OR <1 late enrichment. Bars show 95% confidence intervals. GCATGC (blue) occurred in 46.1% of promoters and had promoter-versus-gene-body OR 1.92 and early-versus-late OR 2.33; GCTAGC (orange) occurred in 12.3% and had corresponding ORs of 2.37 and 0.35. **(F)** Positions of GCATGC (blue) and GCTAGC (orange) in strand-oriented annotated 5′ boundary ±1-kb promoters. Curves show Gaussian-smoothed motif starts per 10 bp per 1,000 promoters (SD = 20 bp). Dashed gray lines and bands show the median and 95% range from 1,000 promoter-wise dinucleotide-preserving shuffles. Beige shading marks −150 to 0 bp. GCATGC peaked at −82 bp, with 32.6% of its sites in the shaded interval; GCTAGC peaked at −84 bp, with 18.3% in that interval. Both positional enrichments had empirical *P* = 0.001 and BH *q* = 0.008. **(G)** GCATGC (blue) and GCTAGC (orange) frequency across inferred accessibility-peak and mRNA-peak times. Open points show half-hour-bin frequencies relative to the average among promoters with a defined peak; curves are descriptive circular smooths and bands are 95% chromosome-bootstrap intervals. The lower plot compares early-versus-late accessibility-peak enrichment among promoters with at least one profiled-factor binding call (*n* = 2,793) and promoters with no such call (*n* = 283). Bars show 95% confidence intervals. GCATGC marked early-peaking occupied promoters (OR 2.53), whereas GCTAGC marked late-peaking occupied promoters (OR 0.33); neither motif showed a resolved early–late bias among promoters without a profiled-factor binding call. **(H)** Factor occupancy at promoters containing GCTAGC (orange) or GCATGC (blue). For each factor–motif pair, a Mantel–Haenszel OR compares occupancy at promoters with versus without the motif within accessibility and other-factor-count strata; bars show 95% confidence intervals. Upward and downward triangles mark ≥1.5-fold higher or lower occupancy odds, respectively, at BH *q* <0.05 across 30 tests; open circles mark all other comparisons. Factor colors describe the expression timing of occupied genes: late-associated, early-associated, or neither. These labels do not specify when a factor binds or acts. This panel uses the independent single-gene-promoter *q* < 0.075 peak calls from the motif-timing analysis rather than the consensus-bound response used in panels B–D. **(I)** Partner features that improved held-out occupancy prediction among the 15 modeled factors. The panel shows 71 directional calls connecting these factors; seven additional calls involving MYB3, AP2XII-9, or ASF1 remained in the 17-partner testing families but are not displayed. The 71 displayed calls collapsed to 39 factor pairs: 32 were predictive in both modeled directions and seven in one direction. Undirected lines denote reciprocal calls; an arrow points toward the factor whose occupancy was predicted for a one-direction call. Line width represents the mean held-out ΔR² ×100 across the displayed direction or directions. Dashed lines mark the two pairs with a partner model OR below one in at least one direction. Node color gives the expression timing of occupied genes. Lines are orange for the seven late–late pairs, which account for 44.9% of the summed displayed gain; blue for the seven early–early pairs, which account for 13.6%; and light green for the remaining 25 pairs, which account for 41.5%. Gray denotes the node class neither; it is not a line class. Neutral background shading shows two descriptive partner groups derived from the gain matrix, not tested biological modules, and carries no timing meaning. Orange halos identify the two strongest reciprocal pairs by mean held-out gain, DMAP1– EPL1 and AAF1–AP2XI-5. Partner associations do not establish simultaneous occupancy, physical interaction, recruitment direction, or regulatory causality. See also **Fig. S11** and **Fig. S12**.

The resulting map was many-to-many: similar PWMs recovered from several factor-specific peak sets often collapsed into the same motif cluster, while individual factor peak sets contributed PWMs to several clusters (**Fig. S11B**). It therefore describes a recurrent promoter-sequence vocabulary rather than a set of factor-exclusive recognition motifs.

To test the predictive power of these features, we fitted a logistic-regression occupancy model for each factor that combined promoter opportunity (chromatin accessibility and promoter architecture) with the number and identities of other bound factors and the shared 72-motif sequence representation (**Fig. 6B** and **Fig. S12A**). The same fixed set of 72 motif features was used for every target factor and every chromosome fold. Each PWM was scanned across both strands of every annotated 5′ boundary ±1-kb promoter, retaining the strongest match per promoter. Within each fit, motif scores were centered and scaled using only the training promoters before prediction on the held-out chromosome. An independent sequence-similarity audit defined a conservative exclusion set of 62 genes linked to promoter homology across chromosomes, including all 55 genes with a direct close match on another chromosome. We excluded this set before model fitting so that a held-out chromosome could not be tested against a near-duplicate promoter already seen during training, leaving 8,116 of 8,178 promoters in the primary analysis. Binding was reconstructed from the CUT&RUN component of the 61 selected network models by retaining their peak and promoter-assignment criteria while removing the knockdown-response requirement. A factor–gene pair was considered consensus-bound when it passed in at least 31 of the 61 models.

For each of 15 factors, the complete occupancy model used all three biological layers to assign an occupancy probability to each promoter. It was fitted on 13 chromosomes and used to predict the omitted chromosome, repeating the procedure until every chromosome had served as the test set (**Fig. 6B**). A prevalence-only reference model instead assigned every promoter the factor’s binding frequency in the training chromosomes. The complete models achieved held-out McFadden R² values of 0.26–0.77; here, held-out McFadden R² measures how much promoter-specific prediction improved over that reference on promoters the model had not seen. We next quantified how well these conditional models ranked bound promoters on omitted chromosomes (**Fig. S13**). Across the 15 factors, complete-model AUROC ranged from 0.849 to 0.987, compared with 0.794–0.974 for ranking promoters by other-factor count (**C**) alone and 0.816–0.973 for accessibility alone; the paired AUROC difference from **C** was 0.003–0.056 (median, 0.014), with chromosome-bootstrap intervals above zero for all 15 factors. Precision ranged from 0.500 to 0.975 (median, 0.816), compared with 0.432–0.965 (median, 0.773) for **C** alone. Adding motifs last produced a median AUROC difference of 0.0008, with intervals above zero for seven factors—all among the 10 factors passing the exploratory log-likelihood layer test (**Fig. S12A**)— whereas the motif-free model exceeded **C** with an interval above zero for all 15; because the 72-motif set was selected using all chromosomes, performance estimates containing **M** may be optimistic.

We then asked two successive questions. First, we used the models fitted with every combination of the three biological layers to divide the held-out predictive information among promoter opportunity, factor environment, and sequence (**Fig. 6C**). Second, we removed one partner or one motif at a time from the complete model, while retaining the remaining features, to pinpoint the partner identities and motif specificities that contributed beyond them (**Fig. 6D**).

Exact Shapley decomposition averaged each biological layer’s added held-out information across every possible order of inclusion (**Fig. 6C**). The surrounding factor environment accounted for 55.1% of the mean complete-model value, promoter opportunity for 39.3%, and sequence for 5.6%. Consistent with this decomposition, removing all motif scores reduced held-out prediction by a mean of 0.00502 bits per promoter, with a positive aggregate sequence contribution on all 14 chromosomes. At the final exploratory Benjamini–Hochberg (BH) q ≤ 0.10 reporting threshold, sequence added information for 10 of 15 factors. Removing the factor environment reduced prediction for all 15 factors, with a mean gain 15.3-fold larger than the sequence gain. Thus, promoter state and the surrounding factor ensemble carried most of the generalizable information about occupancy, while sequence made a smaller, consistently positive contribution concentrated in selected factors.

AP2III-2, HDAC3, and AP2X-7 showed broadly similar relative contributions from promoter opportunity, factor environment, and sequence, but their depletion phenotypes differed. AP2III-2 and HDAC3 produced G1-centered cell accumulation, whereas AP2X-7 accumulated around M/C with a profile extending into G1 (**Fig. 2B**). Thus, a similar balance of predictive layers can characterize factors whose loss disrupts different intervals of the lytic cycle.

We further sought to pinpoint the partner identities and motif features that carried this factor-specific information (**Fig. 6D** and **Fig. S12B**). Removing one partner at a time identified 78 partner features that improved held-out prediction after multiple-testing correction.

The corresponding motif analysis separated summit association from conditional predictive specificity. Eleven factor–motif pairs improved held-out prediction after multiple-testing correction, and AP2XI-4–C23:AYACACAC was the only one that also met the prespecified model odds ratio (OR) threshold (gain, 0.00875 bits per promoter; motif model OR, 1.365 per one-training-set-standard-deviation increase in motif score; **Fig. 6D** and **Fig. S12B**). AP2XI-5– C01:GCTAGC also passed the prespecified internal recovery check. These results show that summit-associated motifs can carry predictive information while demonstrating that sequence association and a unique contribution beyond promoter context are not equivalent.

Predictive sequence relationships were not uniformly positive. Four of the 11 predictive factor– motif pairs had motif model ORs below one, namely AP2IX-7–GCTAGC, AP2IX-5–GCATGC, AP2IX-5–RTGTRCATGCA, and DMAP1–GCATGC, meaning that, after the remaining promoter and factor-context features were held constant, a stronger motif score predicted lower occupancy odds (**Fig. S12B**). These are inverse conditional associations, not evidence that the motifs are absent or actively avoided. The same motif could carry opposite conditional associations for different factors: GCATGC was positive for AP2X-7 but inverse for AP2IX-5 and DMAP1, whereas GCTAGC was positive for AAF1 but inverse for AP2IX-7. Thus, a shared promoter motif need not carry the same predictive information in every factor context.

The smaller average sequence contribution is not evidence that DNA sequence is dispensable. It shows that scores for individual PWMs add comparatively little *marginal* prediction after accessibility, architecture, and factor context are known. Monomeric motifs also do not represent the full cis-regulatory sequence space. Partner-dependent spacing, orientation, and composite motifs can create binding preferences not captured by either factor’s isolated PWM, as demonstrated experimentally in plant heterodimers and in a large human DNA-guided interaction map [29,56]. The *T. gondii* data therefore place motif specificity within a richer promoter context rather than replacing sequence with a sequence-independent model.

AAF1–AP2XI-5 illustrates how partner and sequence information can converge without being equivalent. Removing C01:GCTAGC reduced held-out prediction more strongly for AP2XI-5 than for AAF1 (0.01051 versus 0.00192 bits per promoter), and the corresponding motif model OR was larger for AP2XI-5 (1.36 versus 1.11; **Fig. 6D**). Thus, GCTAGC is associated with both occupancy maps, but the sequence-linked signal is stronger for AP2XI-5, whereas AAF1 occupancy is captured more strongly by their shared factor environment. This asymmetry does not imply that AAF1 directly recognizes the motif.

We next asked whether short promoter motifs marked broader regulatory programs independently of assignment to one factor. Screening all 64 reverse-complement palindromic 6-mers highlighted GCATGC and GCTAGC, two related motifs with opposing temporal associations (**Fig. 6E-G**). GCATGC occurred in 46.1% of promoters and GCTAGC in 12.3%; both were enriched relative to an unpaired, GC-stratified set of gene-body windows (OR, 1.92 and 2.37, respectively) and peaked at nearly the same promoter position, 82–84 bp upstream of the annotated gene start. The position and prevalence of GCATGC closely recapitulate the promoter element identified by high-resolution mapping of *T. gondii* transcription starts [58], while GCTAGC corresponds to the established AP2XI-5 element [17].

Despite their similar position, the two motifs marked opposing phases of the lytic-cycle program. GCATGC was enriched at promoters with early accessibility peaks (early-versus-late OR, 2.33), whereas GCTAGC marked promoters with late peaks (OR, 0.35). The corresponding frequency shifts were visible across both accessibility-peak and mRNA-peak time. They were also concentrated in the occupied promoter landscape: among promoters bound by at least one profiled factor, the early-versus-late ORs were 2.53 for GCATGC and 0.33 for GCTAGC, compared with 0.95 and 0.75 among never-bound promoters. These related motifs therefore define opposing temporal biases rather than interchangeable promoter signals. GCATGC extends the early promoter program associated with AP2XII-8 and ribosomal-protein regulation, whereas GCTAGC connects AP2XI-5 to a broader late program of daughter construction and invasion [6].

To relate factor occupancy to the lytic cycle, we classified each factor once from the wild-type mRNA peak times of genes at its occupied promoters. A factor was late-associated when its occupied promoters were enriched among genes peaking at 4–6 h and not among genes peaking at 0–3 h; early-associated was the converse, and the remaining factors met neither rule. These labels describe the expression timing of occupied genes, not when the factor itself binds. Factor occupancy followed the same temporal sequence logic after controlling for accessibility and total factor burden (**Fig. 6H**). GCTAGC-positive promoters were preferentially occupied by the late-associated factors AP2XI-5, AAF1, and DMAP1 and showed reduced occupancy by AP2X-7 and AP2XII-8. GCATGC-positive promoters showed the reciprocal organization: they were preferentially occupied by AP2XII-8, AP2X-7, HDAC3, and Cdc5 and showed reduced occupancy by AAF1 and AP2IX-5. Sequence-associated promoter classes therefore aligned with distinct groups of regulators even after comparing promoters with similar openness and overall occupancy.

The Cdc5–SNIP1 pair illustrates why promoter-level motif association and factor-specific predictive information must be distinguished. After promoters were stratified by accessibility and the number of other bound factors, GCATGC occurred at 49.4% of Cdc5-bound promoters and 40.5% of unbound promoters (OR, 1.58; 95% CI, 1.25–1.98; BH q = 2.3 × 10⁻⁴; **Fig. 6H**). SNIP1 showed no significant association with either GCATGC or the related GCTAGC motif. In the leave-one-chromosome-out analysis, however, removing GCATGC produced only a small change in Cdc5 prediction and did not significantly reduce held-out performance after correction for the 72 motif tests within Cdc5 (gain, 0.00017 bits per promoter; BH q = 0.97; **Fig. S12B**). The aggregate sequence-layer estimate was likewise unresolved for Cdc5 (gain, 0.00104 bits per promoter; BH q = 0.117; **Fig. 6C**). Thus, GCATGC marks a promoter context associated with Cdc5 occupancy but did not independently improve prediction after the broader promoter and factor context was modeled. Evidence for the Cdc5–SNIP1 partnership instead comes from their coordinated regulatory and occupancy profiles and the experimental tests described above, rather than from a shared factor-specific motif.

**Fig. 6I** displays the 71 calls connecting factors within the 15-member modeled cohort; these collapsed to 39 factor pairs, with 32 supported in both modeled directions and seven in one direction, including two pairs with an inverse directional association (partner model OR <1). The analysis recovered the established HDAC3–MORC complex and the reciprocal AAF1–AP2XI-5 and DMAP1–EPL1 relationships. DMAP1 and EPL1 were each other’s leading partners and formed the strongest reciprocal pair by mean held-out gain. Direction in this analysis indicates which factor was predicted and which partner was omitted; it does not assign recruitment order or regulatory causality.

The partner map also connected the occupancy model to the time-resolved network (**Fig. 3E**, **Fig. 6I**, and **Fig. S12B**). The reciprocal AAF1–AP2XI-5 relationship was retained across most temporal-network windows, whereas Cdc5–SNIP1 was concentrated around the end and beginning of the circular cycle. DMAP1–EPL1 was retained in all 24 windows, supporting its convergence across complementary occupancy and time-resolved views of the regulatory map. These agreements nominate persistent and stage-restricted regulatory partnerships for direct experimental tests without requiring the occupancy model to assign physical interaction.

More broadly, the predictive-partner map brought these relationships together without assigning descriptive groups as tested biological modules (**Fig. 6I**). Line color marked the timing class shared by each pair: seven late–late pairs accounted for 44.9% of the summed displayed held-out ΔMcFadden R², seven early–early pairs for 13.6%, and the remaining 25 pairs for 41.5%. The map recovered persistent links among late-associated factors, including DMAP1–EPL1 and AAF1–AP2XI-5, and a partner group containing the early-associated factors AP2X-7, AP2XII-8, HDAC3, and SNIP1 together with Cdc5 and MORC, which were classified as neither early- nor late-associated. It therefore identifies factor combinations whose occupancy profiles jointly improve prediction, while the motif analyses indicate which of those relationships also coincide with shared or factor-linked sequence signals.

Together, these analyses support an **opportunity-followed-by-selection** model of promoter engagement. Accessibility and architecture define a predictive opportunity landscape, and the surrounding factor environment carries most of the information that distinguishes occupied promoters within that landscape. Sequence adds a smaller layer that was positive in aggregate across all 14 held-out chromosomes and concentrated in selected factors, motifs, and temporally organized promoter classes. “Followed by” describes the organization of predictive information, not a demonstrated causal or temporal recruitment hierarchy.

The central conclusion is therefore not that motifs are unimportant, but that a shared motif vocabulary is not itself a targeting code. Promoter state and the local factor ensemble provide the dominant generalizable context, while selected sequence features and conditional partner associations refine factor-specific occupancy. This layered map converts a complex co-occupancy landscape into explicit, testable hypotheses about which promoter features and factor combinations contribute to target selection.

### 2.7 AP2 associated factor AAF1 is part of the AP2X-5-AP2XI-5 transcriptional complex

One of the strongest connections in the GRN is between AP2XI-5 and AAF1 (**Fig. 3E** and **Fig. 6I**). We assigned the name AP2-associated factor 1 (AAF1) to this previously uncharacterized gene, based on the work presented here. Earlier pull-downs using AP2XI-5 as bait identified AP2X-5 and a hypothetical protein that was not pursued at the time, which corresponds to AAF1 [18]. The proposed model was that the highly cyclical expression profile of AP2X-5 controls the activity and timing of AP2XI-5, which has a more constitutive expression profile. AAF1 likewise has a relatively flat expression profile, with a peak at the mitosis–cytokinesis transition together with AP2XI-5 and AP2X-5 (**Fig. 7A**). Bulk RNA-seq after AP2XI-5 depletion previously highlighted differentially expressed genes controlling the budding scaffold and apical secretory organelles [18]. Here, we observed a similar signature by scRNA-seq, that overlapped with AAF1 response (**Fig. 7B**, **Fig. S14A**). Moreover, the DNA of 93.7% of AAF1-associated genes were also occupied by AP2XI-5, and the shared targets were enriched for apical-organelle and innermembrane-complex associated functions (**Fig. 7B**, **Fig. S14B**; **Table S8**). We validated this observation directly with CUT&RUN experiments of reciprocally depleted factors. Depletion of either AAF1 or AP2XI-5 nearly eliminated detectable DNA association of the reciprocal factor (**Fig. 7C**). Hence, AAF1 and AP2XI-5 are both required for DNA association.

**Figure 7.**
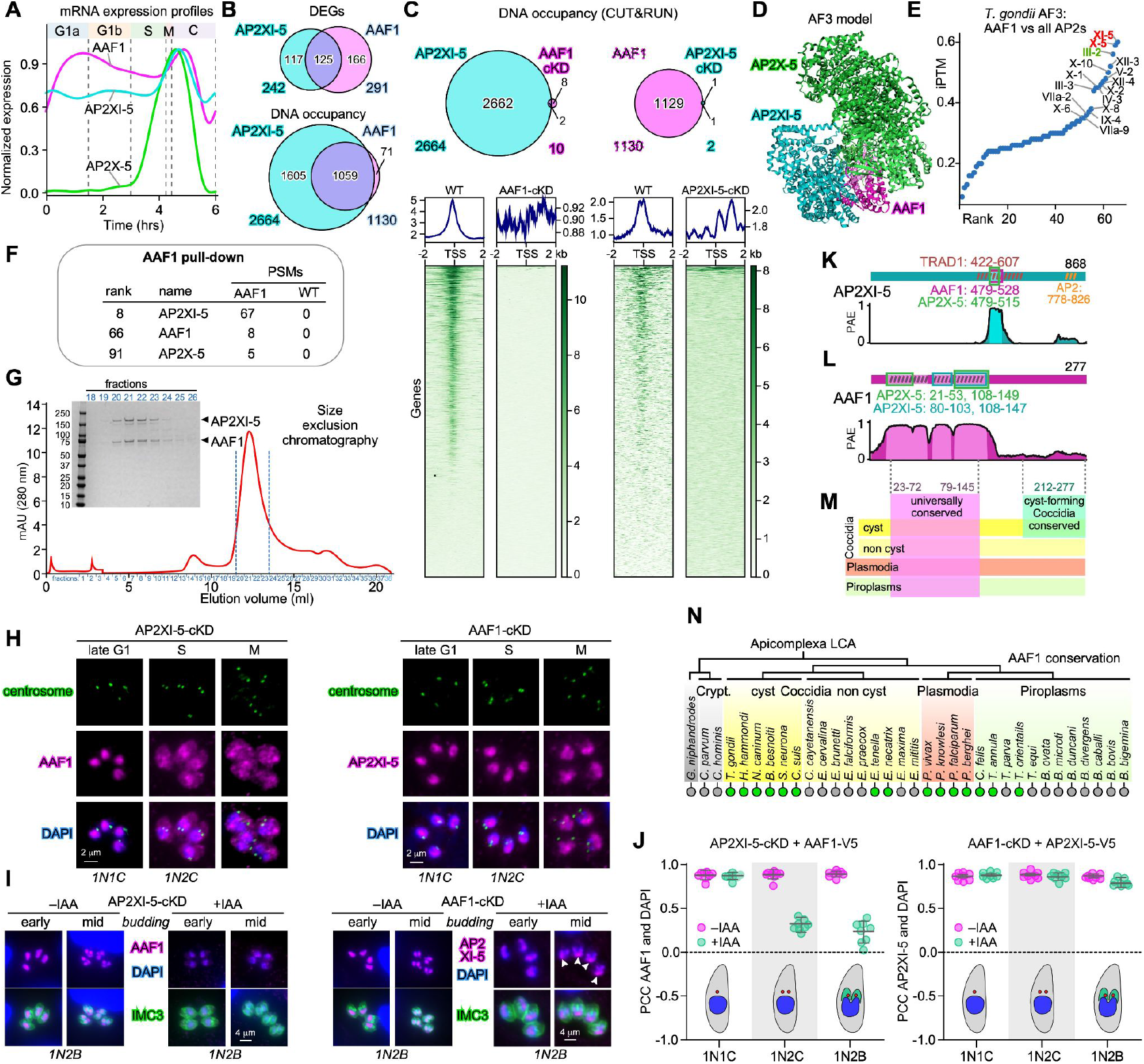
Identification of an AAF1/AP2X-5/AP2XI-5 transcriptional regulatory complex. **(A)** Expression profiles show that AAF1 and AP2XI-5 are constitutively expressed, whereas AP2X-5 expression peaks exclusively around completion of M-phase. Pseudo time wild-type scRNA-seq data. **(B)** Number of coding and non-coding DEGs and genes with shared chromatin occupancy between AP2XI-5 and AAF1 (CUT&RUN data). **(C)** Chromatin occupancy upon reciprocal depletion of AAF1 or AP2XI-5 causes loss of the partner’s DNA binding. Venn diagrams show overlap in genes whereas bottom panels show the occupation density of the targets around the TSS in replete (left; WT) and deplete (right: cKD) conditions or the partner. **(D)** AlphaFold-3 ribbon model of the tripartite complex shows AAF1’s structure conferring effect on the AP2 factors. See **Movie S2** for a rotation of the model. **(E)** AF3 iPTM score of all TgAP2s in complex with AAF1. Other AF3 metrics can be found in **Fig. S14**. **(F)** Summary of complex relevant AAF1 pull-down data. The average of two biological replicates, with three technical replicates each, is listed. The wild type control is the RHΔ*Ku80* parasite strain. PSM: peptide spectrum match. All data in **Fig. S15**. **(G)** Co-expressed and co-affinity purified recombinant AP2XI-5 and AAF1 co-elute on size exclusion chromatography; insert is a CBB stained gel of column fractions, with the peak fractions #19-24 containing both proteins. This demonstrates physical binding of the two proteins. **(H)** AP2XI-5-cKD and AAF1-cKD wherein the reciprocal partner was tagged with an epitope tag to track partner localization through the cell division cycle. Centrosome and DAPI co-staining marks the cell cycle stage as indicated at the top. Parasites were allowed to invade and replicate overnight followed by 6 h IAA treatment. 1N1C: 1 nucleus, 1 centrosome (interphase; G1a/b); 1N2C: 1 nucleus, 2 centrosomes (early S-phase). **(I)** AP2XI-5-cKD and AAF1-cKD herein the reciprocal partner was tagged as marked in panel H. IMC3 (cortical cytoskeleton budding marker) and DAPI co-staining marks the daughter budding stage as indicated at the top. Arrowheads mark AP2XI-5 accumulation at the trailing end of the splitting nucleus, where there is very little DNA as seen by DAPI. I 1N2B: 1 nucleus, 2 buds. All single channel panels provided in **Fig. S17**. **(J)** Pearson’s Correlation Coefficient of DAPI and magenta signals in the conditions indicated. **(K and L)** Mapping the AAF1-AP2XI-5 interaction domain reveals overlap with the previously mapped transactivation domain 1 (TRAD1), conserved in TgAP2XI-5 and orthologous PfAP2-O [60]. Light shading marks regions with predicted high-confidence partner interactions (minimum inter-chain PAE ≤ 6 Å). Colored outlines indicate partner-specific contact regions, with residue ranges labeled. Diagonal hatching denotes annotated or conserved domains, including AP2 (orange) and TRAD1 (brown). Gaps ≤2 residues are bridged for visualization only; residue ranges remain exact. Residue scales are independent. **(M)** AAF1 domain conservation. AAF1 orthologs are present in all cyst forming Coccidia and Plasmodia, but are much spottier outside these two groups. All orthologs share a conserved AP2XI-5 interacting domain. The cyst forming Coccidia shares a conserved AAF1 sub-domain (ccc-region). Magenta: universally conserved AP2 interaction domain; seafoam: region conserved exclusively in the cyst forming Coccidia. **(N)** AAF1 orthologs in Apicomplexa detected by BLASTP search of vEUPathDB using TgAAF1 as bait; green = present; grey = absent.

We next used AlphaFold 3 (AF3) to model putative interactions in this tripartite complex, which revealed a surprisingly extended folding of all three factors (**Fig. 7D** and **Movie S2**). Because AlphaFold 3 can introduce spurious structural order in intrinsically disordered regions, we interpret the detailed geometry of the predicted ApiAP2 structures cautiously [59]. Notably, pairwise interactions between AAF1 and each AP2s produced stronger AF3 scores than the tripartite model (**Fig. S14C**). We experimentally tested the direct interaction between AAF1 and AP2XI-5 using a recombinantly expressed protein pair. Indeed, the two factors migrate as a single complex through a size exclusion chromatography column (**Fig. 7G**), establishing direct physical association and independently supporting the interaction predicted by AF3 by *in vitro* interaction data.

To test whether AAF1 associates indiscriminately with ApiAP2 factors, we obtained AF3 models for every *T. gondii* AP2 and AAF1 pair (**Table S9**). AP2XI-5 and AP2X-5 were the highest-ranking partners across the AF3 metrics examined (**Fig. 7E** and **Fig. S14D-E**). AP2III-2 also consistently scored high with AAF1, which is one of the 21 factors we studied. However, we did not detect a functional relationship between AP2III-2 and AAF1 either in the temporal network or in motif analyses. To directly test whether such interaction might occur in parasites, we performed an AAF1 pull-down experiment, which did detect AP2XI-5 and AP2X-5, but no other AP2 factor (**Fig. 7F** and **Fig. S15**; **Table S10**). Thus, the experimental data support selective association of AAF1 with AP2XI-5 and AP2X-5 in tachyzoites, but not AP2III-2. Mapping the AF3 modeled interaction domains in detail, identified that the single region in AP2XI-5 interfacing AAF1 and AP2X-5 was previously functionally mapped as transactivation domain 1 (TRAD1) in deletion studies [60] (**Fig. 7K**). It is therefore likely that the transactivation capacity of TRAD1 works through AAF1 and/or AP2X-5. From the AAF1 perspective, the interaction interfaces with AP2X-5 and AP2XI-5 overlapped in the central region, but each also included a distinct interaction region in the N-terminal half (**Fig. 7L**, **Fig. S16**).

To obtain dynamic insights at the cell biological level of the AAF1-AP2XI-5 relationship, we performed IFA experiments on parasite lines encoding reciprocally tagged genes with one of the pairs allowing conditional expression. Depletion of AP2XI-5 resulted in a loss of AAF1 signal and a partial redistribution of AAF1 to the cytoplasm (**Fig. 7H-J**; **Fig. S17**). AP2XI-5 remained nuclear regardless of AAF1’s presence, but in late mitotic nuclei it was restricted to the trailing part of the nucleus, where DNA was depleted (arrowheads in **Fig. 7I**). Importantly, in AAF1 replete parasites AP2XI-5 always overlaps with the DNA (compare the ‘mid budding’ panels in **Fig. 7I**). To further pinpoint the timing of AAF1’s shift to the cytoplasm in AP2XI-5 depleted parasites, we co-stained with a centrosome marker to allow differentiation between interphase nuclei (1 centrosome per nucleus) and S-phase nuclei (2 centrosomes per nucleus) (**Fig. 7H**). This revealed that the cytoplasmic shift started at the onset of S-phase (**Fig. 7J**), coinciding with the start of AP2X-5 expression.

The IFA data showed that the DNA association of AAF1 and AP2XI-5 is reciprocally dependent on the presence of the partner. Overall, our data support a model wherein the co-expressed AAF1/AP2XI-5 complex is needed for DNA association, and that nuclear retention of AAF1 requires AP2XI-5 presence during S-phase, when AP2X-5 is expressed. The timing is consistent with competition or partner exchange between AP2XI-5 and AP2X-5 for AAF1. Together with the AAF1 pull-down, the published AP2XI-5 pull-down, and the AF3 interface models, these findings support a tripartite regulatory system in which AAF1 links AP2XI-5 with the periodically expressed AP2X-5.

Lastly, we evaluated the evolutionary footprint of AAF1 across available apicomplexan genomes. This showed that the N-terminal half containing the predicted ApiAP2-interaction surfaces was the most strongly conserved region of AAF1, whereas the C-terminal half is conserved in the close tissue-cyst forming relatives of *T. gondii* (**Fig. 7L-M**). Notably, AP2XI-5 has a functional ortholog in *P. falciparum*, AP2-O, which harbors a conserved TRAD1 domain [60]. AF3 modeling between PfAAF1 and all *P. falciparum* ApiAP2 factors ranked AP2-O as the strongest canidate partner, supporting co-evolution of the AAF1-ApiAP2 pair (**Fig. S14F-G**). What is puzzling however, is that AAF1 orthologs were not universally present across Apicomplexa (**Fig. 7N**). They were absent from the sampled chromerids, gregarines, and cryptosporidia, reflective of the expansion of the ApiAP2 family in coccidian and hemosporidian lineages [61]. But then within these lineages AAF1 was repeatedly lost. Among Coccidia, AAF1 was more consistently retained in species whose sexual and asexual cycles require different hosts, suggesting that it may expand regulatory diversity from a limited set of ApiAP2 factors. Piroplasms, which have a vertebrate and invertebrate vector host, did also lose AAF1, however, which may reflect their overall streamlined genomes and developmental programs. So the most parsimonious model is one wherein AAF1 expands transcriptional versatility of the available AP2 repertoire.

The motif analysis provided a complementary sequence-level view. GCTAGC, originally identified for AP2XI-5 by ChIP-on-chip, was enriched in both the AP2XI-5 and AAF1 occupancy maps (**Fig. 6D**) [17]. In the leave-one-chromosome-out predictive analysis, however, GCTAGC remained independently informative for AP2XI-5 after the surrounding factor context was added, but not for AAF1. Thus, AP2XI-5 carries the stronger factor-linked sequence signal, whereas AAF1 occupancy is explained more strongly by the shared promoter environment. This result reinforces the extensive overlap in the genes engaged by AP2XI-5 and AAF1 without implying that AAF1 directly recognizes the motif.

Overall, AAF1 is a previously uncharacterized factor that co-evolved with selected ApiAP2 transcription factors following ApiAP2 family expansion in Coccidia and Hemosporidia. The overlap between the predicted AAF1-interaction surface and the functionally defined AP2XI-5 TRAD1 region suggests that AAF1 may contribute to the transactivation capacity of the complex. Our future work will test this hypothesis. Together, the physical, localization, and chromatin-ssociation data identify AAF1 as a shared partner that stabilizes AP2XI-5 promoter engagement and links it to the cell-cycle-restricted AP2X-5 program.

## 3 Discussion

Landmark bulk and single-cell studies established the ordered, just-in-time transcriptional cascade of apicomplexan replication [6–8,10,11]. Work on individual ApiAP2 factors, including AP2IX-5, AP2XI-5 and AP2XII-8, further showed that successive regulators control daughter budding, invasion-organelle biogenesis, and ribosomal-protein programs [6,15,17,18]. Our map agrees with this cascading logic but departs from a simple relay in which one factor controls one phase or one regulon. At the individual-gene level, most profiled factors were associated with both activation and repression in different contexts, and genes with similar expression timing separated into communities with distinct regulator combinations. These communities provide a regulatory architecture for coordinating DNA replication, mitosis, daughter assembly, and organelle biogenesis as overlapping processes. However, organelle specific gene repertoires span multiple regulatory communities, which provided the modular flexibility to assemble developmental stage-specific organelle protein compositions [1,5]. To illustrate the biological rationale, consider the AMA protein family. AMA proteins are secreted from the micronemes and are required to establish the moving junction facilitating host cell invasion. Tachyzoites use AMA1 but in bradyzoites deploy a combination of AMA2 and AMA4 [62], whereas AMA3 dominates in sporozoites [63]. Hence, stage specific organelle proteomes require customized gene community combinations, resulting in a mosaic of organelle communities.

The map also revises how ApiAP2 target gene selection should be framed. Protein-binding studies established that ApiAP2 DNA-binding domains recognize defined sequences, and individual *T. gondii* factors engage biologically important promoter motifs [13,14,17,18]. Those findings establish sequence preference, but do not establish that a motif alone specifies which promoters are occupied *in vivo*. Across the factors modeled here, accessibility and promoter architecture described regulatory opportunity, while the surrounding factor environment and selected sequence features refined target choice. Sequence remained informative for particular factors and promoters, but no universal one-factor/one-motif code explained occupancy. This opportunity-followed-by-selection model agrees with broader eukaryotic evidence that chromatin state and combinatorial factor binding bridge biochemical sequence recognition and genomic occupancy [28,29,55,56]. It also extends the cooperative and competing ApiAP2 relationships described in *T. gondii* [18,27]. The central question is therefore no longer simply which motif a factor can bind, but which promoter states and factor combinations allow that sequence information to select a target. This is a predictive organization, not yet a causal recruitment hierarchy.

Several relationships illustrate how the map can move from regulatory architecture toward mechanism. Recovery of the established MORC-HDAC3 relationship provided an internal biological anchor [21]. Cdc5 and SNIP1 associated biochemically, each lost most detectable promoter occupancy after depletion of the other, and both contributed to pre-mRNA splicing, supporting a shared complex while leaving its proposed transcriptional role to be tested. Notably, the Cdc5/SNIP1 association was not part of our starting hypothesis as we included SNIP1 on basis of its cMyc co-factor function. Hence, our approach can make new functional connections at multiple levels of gene expression control, illustrated here by an mRNA splicing complex.

The second regulatory relationship we uncovered is between AAF1 and AP2XI-5, which cooperate additionally with AP2X-5 to regulate invasion-organelle genes. Our observations support a lineage-specific regulatory assembly. We show that AAF1 is required for AP2XI-5’s transactivation function and vice versa, and that AP2X-5 previously shown to associate with AP2XI-5 affects the regulatory capacity of this tripartite complex. Against the backdrop that ApiAP2 proteins are typically large with ∼2000 aa with the 50-60 aa AP2 domain the only reconizable feature in an otherwise predicted disordered sequence. Against this backdrop, our finding provides a first insight in how transactivation function could be relayed to the basal RNA polymerase II machinery. Most notably, the overlap between the predicted AAF1-AP2 interaction surface and the functionally defined AP2XI-5 TRAD1 region suggests that AAF1 may convey the transactivation capacity of the complex. Whether this discovery of a potentially auxiliary AP2 transactivation factor is a blueprint for how other ApiAP2s convey their function remain open. Either way, the AlphaFold3-modeled AAF1-ApiAP2 interactions do reveal that specific auxiliary factors could impose structure on ApiAP2 factors that may be required for their function.

An additional candidate complex is suggested by the DMAP1-EPL1 relationship. We selected these factors agnostically on the basis of their Myb and polycomb-like DNA binding domains, respectively. Their strong association throughout the time-resolved and DNA motif networks led us to the NuA4 complex which contains both factors [64]. The NuA4 complex is a histone acetyl transferase (HAT) complex that shares many subunits with the SAGA complex. *T. gondii*‘s SAGA complex was recently resolved [65]. Notably, the GCN5b component interacts with two different AP2 complexes (AP2XII-4/AP2VIIa-5 and AP2IX-7/AP2X-8 [25]) through apicomplexan specific components rather than the canonical Tra1 protein that interacts with transcription factors in higher eukaryotes [64]. Since in higher eukaryotes Tra1 is the shared factor between SAGA and NuA4 complexes, it is an interesting question on how this will look in *T. gondii*. None of the ApiAP2 factors we profiles interacted functionally with EPL1/DMAP1. Hence, this is as an example of a new hypothesis generated by the established network.

Overall, by placing perturbation, occupancy, chromatin state, sequence, and molecular validation on one cell-cycle clock, this work converts the long-recognized just-in-time cascade from an expression pattern into an experimentally navigable regulatory architecture. The map shows how functional modules are deployed, identifies the factors and candidate molecular assemblies associated with their control, and makes the remaining causal questions explicit. Together with the interactive resource, it transforms the multidimensional data into a hypothesis-generation engine for discovering gene function and dissecting how transcription factors, chromatin regulators, and promoter context cooperate as an apicomplexan parasite builds the next invasive cell.

## 4 Methods

### Key resources table

Refer to **Table S11**, **Table S12**, **Table S13**, **Table S14** and **Table S15** for information on key resources

### Resource availability

#### Materials availability

Materials as in **Table S11** and **Table S12** available upon request.

#### Data and code availability

The data can be explored in the ToxoHub interactive application at https://umbibio.math.umb.edu/toxohub/v4/.

### Experimental model details

#### Parasites and host cell culture

All parasite strains used for this study are listed in **Table S11**. *T. gondii* tachyzoite cultures were maintained in human telomerase reverse transcriptase (hTERT)-immortalized human foreskin fibroblasts (HFFs) as previously described [66]. Selection for stable parasite transfectants was performed with 1 µM pyrimethamine and stable lines were cloned by limiting dilution. Protein knock-downs using the minimal Auxin-Inducible Degron (mAID) system were performed with 500 µM 3-indoleacetic acid (IAA; Sigma-Aldrich) in 100% ethanol.

### Method details

### Experimental methods

#### Generation of transgenic strains

All oligonucleotides and parasite strains and used in this study are listed in **Table S11**, **Table S12** and **Table S14**. *T. gondii* tachyzoites were propagated in human foreskin fibroblasts (HFFs) immortalized by human telomerase reverse transcriptase (hTERT), as previously described [66].

For conditional protein depletion, parasite strains carrying either N- or C-terminal minimal auxin-inducible degron (mAID) tags were generated using CRISPR-Cas9-mediated genome editing. Stable transgenic parasites were selected with 1 μM pyrimethamine and subsequently cloned by limiting dilution. Degradation of mAID-tagged proteins was induced by treatment with 500 μM indole-3-acetic acid (IAA; Sigma-Aldrich), prepared in absolute ethanol. Treatment durations varied among experiments and are specified in the corresponding figure legends.

To establish TetO7-SAG4 conditional knockdown strains, the endogenous promoters of target genes were replaced with the anhydrotetracycline (ATc)-regulated TetO7-SAG4 promoter using CRISPR-Cas9-mediated genome editing, together with the introduction of an N-terminal Ty tag.

For C-terminal tagging, a 10×Myc epitope was introduced at the endogenous loci of target genes using CRISPR-Cas9-mediated genome editing and the pLIC-SM-Myc construct. Stable transgenic parasites were selected with 20 μM chloramphenicol and cloned by limiting dilution.

For N-terminal epitope tagging, the HXGPRT-T2A-5×V5, HXGPRT-T2A-3×Myc, and HXGPRT-T2A-ALFA constructs were used to introduce 5×V5, 3×Myc, and ALFA tags, respectively, at the endogenous loci of target genes through CRISPR-Cas9-mediated genome editing [53,67]. Stable transgenic parasites were selected with 25 μg/mL mycophenolic acid (MPA) supplemented with 50 μg/mL xanthine and subsequently cloned by limiting dilution.

For *P. falciparum*, the conditional KD construct, pKD-PfCdc5-TetR-DOZI, was generated using the method and plasmid described in Rajaram et al. 2020 [68]. Two days before transfection, a high-parasitemia culture (≥5%) was pellet down (500 × g, 5 min) and resuspended in 5 mL of 5% sorbitol and incubated for 15 min to synchronize parasites at the ring stage. Cells were pelleted again (500 × g, 5 min), washed three times with RPMI 1640, resuspended in complete RPMI, and plated in a 30 mL dish at 2% haematocrit (HCT) and incubated at 37 °C. One day before transfection, the medium was replaced with fresh medium to obtain a high-parasitemia and placed in a shaking incubator at 37°C.

On the day of transfection, parasitemia was assessed by blood smear and was ∼10% ring stages. DNA mixtures were prepared by adding 18 µL of Supplement and 82 µL of P3 solution (Lonza P3 Primary Cell 4D-Nucleofection Kit, V4XP-3024) to 50 µg of sterile DNA. The culture was pellet down in a 50 mL tube (500 × g, 8 min), and the Lonza Amaxa X unit and FP158 protocol for P3 primary cells was used. After removing the supernatant, 70 µL of infected RBCs (iRBCs) was taken slowly from the bottom of the tube, mixed with the DNA/P3/Supplement solution, and mix was transferred into Amaxa Nucleocuvette for transfection. The transfected contents were immediately transferred to a petri dish containing 10 mL of warm complete RPMI and 300 µL of fresh human RBCs, then incubated at 37°C. One day post-transfection, a blood smear was taken to assess lysis, the medium was changed, and drug selection was started in a shaking incubator. From day 3 onward, the medium was changed daily for 3 days with a small amount of blood (0.5% HCT) added, then every 2–3 days. Parasites were split 1:2 about every 7 days to keep the blood fresh and emerged about 3 weeks after transfection. The mutant culture was maintained at 500 nM ATc, and conditional knockdown (cKD) was induced by washing the cells thrice to remove the ATc from the culture.

#### Immunofluorescence microscopy

Tachyzoites were cultured overnight in confluent human foreskin fibroblast (HFF) monolayers grown on glass coverslips in 6-well plates. Infected cells were fixed with 4% paraformaldehyde (PFA), permeabilized with 0.25% Triton X-100 for 30 min, and blocked with 0.3% bovine serum albumin (BSA) in 1× phosphate-buffered saline (PBS) for 30 min. Primary antibodies used in this study are listed in **Table S13**. Alexa Fluor 594-conjugated anti-mouse and Alexa Fluor 488-conjugated anti-rabbit secondary antibodies were diluted 1:400 in blocking buffer and incubated for 1 h at room temperature. Following secondary antibody incubation, samples were washed three times with 1× PBS for 5 min each, with 4′,6-diamidino-2-phenylindole (DAPI) included in the first wash to stain DNA. Fluorescence images were acquired using a DeltaVision Ultra microscope and processed using Fiji (ImageJ) software. For high-resolution phenotypic analysis of double-tagged parasite strains, images were acquired using a Zeiss Axio Imager Z2 microscope equipped with an EXFO LED illumination system. Phenotypic quantification was performed by examining at least 100 parasitophorous vacuoles (PVs) per experiment across three independent biological replicates. Colocalization analysis was performed using the Coloc 2 plugin in Fiji (ImageJ). Regions of interest (ROIs) were manually defined for individual parasites based on the inner membrane complex (IMC) fluorescence channel. The same ROIs were applied to the DAPI and magenta fluorescence channels for colocalization analysis. Pearson’s correlation coefficients (PCCs) were calculated from pixel intensity correlations within each ROI using the unthresholded Pearson’s R values reported by Coloc 2.

#### Single-cell RNA sequencing (scRNA-seq)

For *T. gondii*, intracellular RHΔKu80 tachyzoites were harvested at 27 and 30 h post-infection (hpi). For auxin-inducible degradation experiments, infected human foreskin fibroblast (HFF) monolayers were treated with indole-3-acetic acid (IAA) for 2 h before parasite harvesting. Infected monolayers were washed three times with ice-cold phosphate-buffered saline (PBS), and parasites were released by cell scraping, mechanical disruption through a 26-gauge needle, and filtration through a 3-µm filter to remove host-cell debris. Parasites were maintained on ice throughout sample preparation. For each sample, 5 × 10⁶ parasites were collected by centrifugation at 1,000 × g for 15 min at 4°C, washed twice with ice-cold Hanks’ balanced salt solution (HBSS) containing 0.04% bovine serum albumin (BSA), and resuspended at 1 × 10⁶ parasites/mL in the same buffer. Equal numbers of parasites from the 27 and 30 hpi samples were pooled before library preparation.

For *P. falciparum*, NF54 parasites maintained in 20 nM anhydrotetracycline (ATc) were divided into two equal cultures. ATc was removed from one culture by three washes to induce conditional knockdown (cKD), while the control culture was maintained in 20 nM ATc. Both cultures were incubated for 20 h at 37°C, resulting in parasitemia levels of 3% (+ATc) and 2% (−ATc), respectively.

Single-cell 3′ gene expression libraries for both species were generated using the Chromium Next GEM Single Cell 3′ Reagent Kit v3.1 (Dual Index, 10x Genomics) according to the manufacturer’s instructions, with approximately 16,500 cells loaded per channel to target a recovery of 10,000 cells.

For *T. gondii*, sequencing data were processed and analyzed as described in Section “Single-cell RNA-seq data processing”. For *P. falciparum*, empty droplets and doublets were removed as described in Section “Velocity along pseudotime (*Pf*Cdc5)”. Filtered cells were clustered and assigned to intraerythrocytic developmental cycle (IDC) stages based on Malaria Cell Atlas annotations [69]. The cKD-specific population, designated “perturbed cells,” was compared separately with ring-stage and schizont-stage populations to identify differentially expressed genes. These genes were expanded according to the number of annotated introns in the *P. falciparum* genome. Gene Ontology (GO) enrichment analysis was performed separately for upregulated and downregulated genes from each comparison.

#### Single-cell multiome (scMultiomics)

Tachyzoites were harvested as described for scRNA-seq above. For nuclei isolation, parasite pellets were gently resuspended in 100 μL of ice-cold ATAC Lysis Buffer (Active Motif) freshly supplemented with 1 mM dithiothreitol (DTT) and 1 U/μL RNase inhibitor. The lysates were subsequently diluted with 1 mL of ice-cold wash buffer containing 10 mM Tris-HCl (pH 7.4), 10 mM NaCl, 3 mM MgCl₂, 1% bovine serum albumin (BSA), 0.1% Tween 20, 1 mM DTT, and 1 U/μL RNase inhibitor. Nuclei were collected by centrifugation at 1,500 × g for 10 min at 4 °C using a swinging-bucket rotor and washed twice more under the same conditions. The resulting nuclei pellets were resuspended in 20 μL of chilled diluted nuclei buffer containing 1× Nuclei Buffer (10x Genomics), 1 mM DTT, and 1 U/μL RNase inhibitor. Nuclei concentrations were determined using a hemocytometer and adjusted to 4,000–4,600 nuclei/μL. Nuclei isolated from parasites harvested at 27 and 30 hpi were pooled for subsequent multiome library preparation. For each library, 3.5 μL of nuclei suspension at 4,600 nuclei/μL (or 4.025 μL at 4,000 nuclei/μL) was allocated for the transposition reaction according to the 10x Genomics Multiome protocol (CG000338, Rev F), targeting a recovery of 10,000 nuclei per library.

#### CUT&RUN

CUT&RUN was performed using the CUTANA™ ChIC/CUT&RUN Kit Version 4 (EpiCypher; User Manual Version 4.1), with modifications for *T. gondii*. Intracellular RH tachyzoites were cultured in confluent human foreskin fibroblast (HFF) monolayers in two T175 flasks per sample and harvested approximately 30–36 h post-infection, before extensive host-cell lysis. Infected monolayers were washed once with ice-cold phosphate-buffered saline (PBS), scraped into 35 mL of ice-cold Ed1 medium per flask, and mechanically disrupted by passage through a 26-gauge needle. Parasites were purified by filtration through a 3 µm filter to remove host-cell debris and counted using a hemocytometer. Approximately 2 × 10^8^ parasites were collected per CUT&RUN reaction by centrifugation at 1,000 × g for 15 min at 4°C.

Purified parasites were immobilized on activated concanavalin A (ConA) magnetic beads and permeabilized using digitonin-containing buffer according to the manufacturer’s protocol. Samples were incubated with primary antibodies overnight at 4°C. Anti-Ty (BB2), anti-V5, or anti-Myc antibodies (2.0 µg per reaction) were used to profile the corresponding epitope-tagged proteins. Anti-H3K4me3 and IgG1 antibodies were included as positive and negative controls, respectively.

Following antibody incubation, samples were washed twice with cell permeabilization buffer and incubated with 2.5 µL of protein A/G–micrococcal nuclease (pAG-MNase) for 10 min at room temperature. After two additional washes, targeted chromatin cleavage was initiated by adding 1 µL of 100 mM CaCl_2_ per reaction, followed by incubation for 2 h at 4°C. The digestion reaction was terminated by adding stop buffer, and samples were incubated at 37°C for 10 min to facilitate the release of cleaved chromatin fragments.

Following magnetic separation, supernatants containing released CUT&RUN DNA fragments were collected. DNA was purified using solid-phase reversible immobilization (SPRI) magnetic beads according to the manufacturer’s protocol.

Sequencing libraries were prepared from purified CUT&RUN DNA using the CUTANA™ CUT&RUN Library Prep Kit (EpiCypher), following the manufacturer’s instructions. Library preparation included end repair, adapter ligation, dual indexing, PCR amplification, and bead-based purification. Libraries were sequenced on an Illumina NextSeq 2000 platform using a P1 100-cycle reagent kit.

#### Plaque assays

Confluent human foreskin fibroblast (HFF) monolayers grown in 6-well plates were inoculated with 50 *T. gondii* tachyzoites per well and allowed to form plaques for 7 days without disturbance. Following the 7-day incubation, HFF monolayers were fixed with 100% ethanol for 10 min and stained with crystal violet for 5 min to visualize plaques.

#### Western blot analysis

Antibodies used are listed in **Table S13**. For *T. gondii*, intracellular parasites were harvested by mechanical disruption of infected host cells through a 26-gauge needle, followed by filtration through a 3 µm pore-size filter to remove host-cell debris. Purified parasites were washed three times with phosphate-buffered saline (PBS) and lysed in a buffer containing 50 mM Tris-HCl (pH 7.3), 150 mM NaCl, protease inhibitor cocktail (1:50 dilution; Sigma-Aldrich, P8849), and 1% SDS. Whole-cell lysates corresponding to 2 × 10⁷ parasites were resolved on NuPAGE 4–12% Bis-Tris gels (Invitrogen, NP0321BOX) and transferred onto polyvinylidene difluoride (PVDF) membranes (Thermo Scientific, 88520). Membranes were blocked overnight at 4°C in PBS containing 5% nonfat dry milk, 1% bovine serum albumin (BSA), and 0.1% Tween 20. Membranes were incubated with primary and secondary antibodies for 1 h each at room temperature, with three 10-min washes in PBS containing 0.1% Tween 20 (PBST) after each incubation. Chemiluminescent signals were developed using SuperSignal West Femto Maximum Sensitivity Substrate (Thermo Scientific, 34095) and visualized using an Odyssey M imaging system (LI-COR).

For *P. falciparum*, tightly synchronized schizont-stage parasites were collected at 45 and 99 h following anhydrotetracycline (ATc) removal. Infected erythrocytes were treated with ice-cold 0.03% (w/v) saponin in PBS for 15 min on ice, followed by centrifugation at 3,300 × g for 10 min. Parasite pellets were washed three times with ice-cold PBS containing EDTA-free protease inhibitors (cOmplete, Roche) and resuspended in 20 volumes of the same buffer. Samples were mixed with 4× LDS sample buffer and 10× reducing agent (Invitrogen) and denatured at 95°C for 10 min. Proteins were separated on 4–20% Mini-PROTEAN TGX precast gels (Bio-Rad) using Tris/glycine/SDS running buffer and transferred onto nitrocellulose membranes using a Trans-Blot Turbo system (Bio-Rad). Membranes were blocked for 1 h at room temperature in Intercept (PBS) blocking buffer (LI-COR) and incubated with mouse anti-HA and rabbit anti-Pfaldolase antibodies for 1 h, followed by goat anti-mouse IRDye 800CW and donkey anti-rabbit IRDye 680CW secondary antibodies for 1 h. Membranes were washed three times with PBS containing 0.05% Tween 20 after each antibody incubation. All antibody incubations and washes were performed at room temperature, and fluorescence signals were detected using an Odyssey CLx imaging system (LI-COR).

#### Co-immunoprecipitation

Intracellularly replicating *T. gondii* tachyzoites were harvested at 30 h post-inoculation by passage through a 26-gauge needle and pelleted by centrifugation at 1,000 × g for 15 min at 4°C. Pellets were washed under the same centrifugation conditions and resuspended in 200 µL of ice-cold lysis buffer containing 10 mM Tris-HCl, 150 mM NaCl, 0.5 mM EDTA, 0.5% IGEPAL CA-630, and protease inhibitors. Samples were briefly vortexed and bath-sonicated for 15 min at 4°C. Lysates were clarified by centrifugation at 1,800 × g for 5 min at 4°C, and the resulting supernatants were collected for downstream analysis.

Reciprocal Co-IP experiments were performed using Cdc5-Myc/SNIP1-Ty and SNIP1-Myc/Cdc5-Ty parasite strains. For immunoprecipitation, 25 µL of pre-equilibrated Myc-Trap® Magnetic Agarose beads (ChromoTek) were added to the cleared lysates and incubated for 1 h at 4°C with gentle end-over-end rotation. Beads were subsequently washed three times with 500 µL of wash buffer containing 50 mM Tris-HCl, 300 mM NaCl, 1 mM EDTA, and 0.1% Triton X-100. Bound proteins were eluted by heating the beads in 2× SDS sample buffer at 95 °C for 10 min.

Total lysate (TL), flow-through (FT), wash fractions (W1–W3), and eluates (E) were analyzed by SDS–PAGE and western blotting using anti-Myc, anti-Ty, and anti-GRA1 antibodies. Co-IP experiments were performed in three independent biological replicates. Western blotting was used for qualitative assessment of protein co-immunoprecipitation, without quantitative normalization of band intensities. Co-immunoprecipitation was considered supported when the Ty-tagged partner was detected in the Myc-Trap eluate in both reciprocal experiments, together with recovery of the corresponding Myc-tagged bait. GRA1 immunoblotting was used to assess non-specific protein carryover into the eluate.

#### Pull-down and mass spectrometry

Intracellularly replicating *T. gondii* parasites were grown in a T175 flask and harvested when host cells were heavily infected (approximately 30 h post-infection). Parasites were released by passage through a 26-gauge needle, filtered through a 3 µm membrane to remove host-cell debris, and counted prior to collection. Parasite pellets were washed three times with ice-cold PBS by centrifugation at 1,000×*g* for 10 min at 4°C.

Parasites were lysed in an ice-cold lysis buffer supplemented with a protease inhibitor cocktail. Lysates were briefly vortexed, subjected to bath sonication at 4°C, and clarified by centrifugation at 1,800×*g* for 5 min at 4°C. The clarified lysates were transferred to low protein-binding micro-centrifuge tubes and diluted with dilution buffer supplemented with protease inhibitors.

For affinity purification, Myc-Trap® Magnetic Agarose beads (ChromoTek) were equilibrated according to the manufacturer’s instructions and incubated with the clarified lysates for 1 h at 4°C with gentle end-over-end rotation. Following incubation, the beads were washed three times with a wash buffer to remove nonspecifically bound proteins. During the final wash, the beads were transferred to a fresh low protein-binding tube to minimize sample contamination.

Bound proteins were eluted with 8 M urea in PBS by heating at 95°C for 10 min. The beads were subsequently washed twice with ammonium bicarbonate buffer, and the wash fractions were combined with the urea eluate. The pooled samples were reduced with dithiothreitol (DTT), alkylated with iodoacetamide (IAA), diluted with PBS, and digested overnight with sequencing-grade trypsin at 37°C. Digestion was terminated by the addition of formic acid, and insoluble material was removed by high-speed centrifugation. The resulting peptide-containing supernatants were collected for LC–MS/MS analysis.

Prior to mass spectrometry, peptide samples were desalted using C18 solid-phase extraction cartridges. Columns were conditioned with acetonitrile, equilibrated with low-pH buffer A, loaded with peptide samples, washed extensively with buffer A, and eluted with low-pH buffer B. Eluted peptides were dried by vacuum centrifugation before LC–MS/MS analysis.

**LC–MS/MS and protein identification.** LC-MS/MS was performed on an Orbitrap Exploris 240 mass spectrometer with Xcalibur version 4.4 (Thermo Fisher Scientific) coupled to a Dionex UltiMate 3000 RSLCnano system. Desalted peptides were resuspended in 100% water buffer A (100% H_2_O and 0.1% formic acid) and loaded onto an Acclaim PepMap 100C18 loading column (Thermo Scientific) for further elution onto an Acclaim PepMap RSLC column (Thermo Scientific), where the peptides were separated with a 2-h gradient ranging of 5–25% buffer B (20% H2O, 80% acetonitrile, and 0.1% formic acid) in buffer A at a flow rate of 0.3 µL/min. The spray voltage was set to 2.1 kV. One full MS1 scan (120,000 resolution, 350-1,800 m/z, RF lens 65%, automatic gain control (AGC) target 300%, automatic maximum injection time, profile mode) was obtained every 2 s with dynamic exclusion (repeat count 2, duration 10 s), isotope exclusion (assigned), and apex detection (30% desired apex window) enabled. A variable number of MS2 scans (15,000 resolution, AGC 75%, maximum injection time 100 ms, centroid mode) were obtained between each MS1 scan based on the highest precursor masses, filtered for monoisotopic peak determination, theoretical precursor isotopic envelope fit, intensity (5E4), and charge state (2–6). MS2 analysis included the isolation of precursor ions (isolation window 2 m/z) followed by higher-energy collision dissociation (HCD) (collision energy 30%).

LC-MS/MS data were analyzed on Thermo Proteome Discoverer version 2.4 and searched using SequestHT and Percolator algorithms against a *Toxoplasma gondii* protein database. The protease for data analysis was specified as trypsin with a maximum of 2 missed cleavages. Peptide precursor tolerance was set to 10 ppm with a fragment mass tolerance of 0.02 Da. Cysteine alkylation (+57.021) was set as a static modification, methionine oxidation (+15.995), acetylation (+42.011), and methionine loss (+131.040) of the protein N-terminus were set as dynamic modifications. The false discovery rate (FDR) was set to 1% for peptide identification.

#### Recombinant protein expression and binding

**Constructs.** TgAAF1 and TgAP2XI-5 gene sequences were codon-optimized the for expression in Expi293F cells and had them synthesized by Twist Bioscience. Using a ligation-independent cloning approach, TgAAF1 and TgAP2XI-5 DNA sequences were fused 3’ to a FLAG_3_-His_6_- MBP-3C solubility tag (MBP: maltose-binding protein, 3C: human rhinovirus cysteine protease cleavage site) within the backbone of a pCAGGS derivative vector [70] (Creative Biogene). Each plasmid generated (pCAGGS-FLAG_3_-His_6_-MBP-3C-TgAAF1/AP2XI-5) was sequenced to confirm its accuracy.

**Protein expression and complex purification by SEC.** Recombinant FLAG_3_-His_6_-MBP-3C-TgAAF1/TgAP2XI-5 proteins were co-expressed using the mammalian suspension Expi293F system (Gibco^TM^). Briefly, we cultured 30-60 mL Expi293F cells seeded at 1.5 × 10^6^ cells/ml in Expi293TM Expression Medium (Gibco). Expi293F cells were transfectd with 30-60 µg of the appropriate plasmid using ExpiFectamine (80-160 µg, Thermo Fisher). Cultures were maintained in a shaker incubator (37°C, 8 % CO_2_) for 48 hours. Enhancer 1 (150-300 µl) and Enhancer 2 (1.5-3 mL) were added to the medium 16 h post-transfection. Cells were harvested and lysed 48 h post-transfection with lysis buffer (LB) at a 1:1 (v/v) ratio with the culture volume (LB: 50 mM HEPES (pH 7.5), 300 mM NaCl, 0.5 mM EDTA, 0.1% Triton X-100, 1 mM DTT, eComplete protease inhibitor tablets (Roche) for 1 hr at 4°C. Cell debris was removed by centrifugation (10,000 x g at 4°C, 30 min) and the clarified lysate incubated with a pre-equilibrated amylose resin (NEB) for one h at 4°C on a nutator. Recombinant proteins were purified by gravity flow, washing with 15 column volumes (CV) of LB lacking Triton X-100. We eluted tagged recombinant proteins using 12-15 mL of elution buffer (EB) (EB: 50 mM HEPES (pH 7.5), 300 mM NaCl, 0.5 mM EDTA, 1 mM DTT, 30 mM Maltose). When required, the N-terminal FLAG_3_-His_6_-MBP-3C tag was removed using an on-bead cleavage method by adding glutathione S-transferase-3C (GST-3C, Acro Biosystems) protease for 2.5 h at room temperature, followed by overnight incubation at 4°C (one unit of GST-3C for 100 µg of protein after the wash step). After cleavage, untagged proteins were recovered by collecting the flow-through. Samples were concentrated using a 10 kDa cutoff (Amicon Ultra filter, Millipore Sigma). Recombinant proteins were further purified by size-exclusion chromatography (SEC) using a Superdex Increase 10/300 GL column (Cytiva) equilibrated with SEC buffer (SECB: 50 mM HEPES (pH 7.5), 200 mM NaCl, 0.5 mM EDTA, 1 mM DTT). Protein purity and complex presence was assessed by SDS-PAGE analysis and identity of TgAAF1 and TgAP2XI-5 confirmed by submitting destained, excised gel bands corresponding to each recombinant protein to the Taplin Mass Spectrometry Facility (Harvard Medical School).

#### Ortholog PfCdc5 in P. falciparum

**ATc titration and phenotype assessment:** The pKD-PfCdc5-TetR-DOZI mutant was grown at 500, 100, 20, 4, 1 and 0 nM ATc for 165 h, with parasitaemia monitored at 19, 41, 69, 96, 117, 141 and 165 h post induction, to identify the lowest concentration capable of maintaining the mutant culture growing and to determine how many cycles the parasite takes to show the phenotype (**Table S16**). Control and cKD cultures were subsequently maintained at ± 20 nM ATc and assessed at 20 h for schizont segmentation.

Within the cKD sample, perturbed cells were compared to rings and schizonts. Differentially expressed genes from each comparison were expanded by the number of introns present using the *P. falciparum* genome annotation. GO enrichment was performed separately on the upregulated and downregulated genes of each comparison (**Table S5** and **Table S6**).

### Computational methods

#### Single-cell RNA-seq data processing

**Libraries, reference and quantification.** Twenty-two single-cell RNA-seq libraries were analyzed: one wild-type RH library (WT) and one library for each of 21 knockdown (KD) lines. All libraries were generated using the 10x Genomics Chromium Single Cell 3′ v3 gene-expression chemistry (**Table S2**); library preparation and sequencing are described in Section “Single-cell RNA sequencing (scRNA-seq)”. The alignment reference was built from the *T. gondii* ME49 assembly and annotation in ToxoDB release 68 [71]. The GFF annotation was converted to GTF with AGAT [72], and a reference containing 8,778 genes was generated with Cell Ranger mkref [73]. Reads were aligned and counted separately for each library using Cell Ranger count with default settings. Two Cell Ranger versions were used across the 22 libraries (**Table S15**; **Fig. S6A**).

**Cell filtering.** A common filtering procedure based on read-level alignment statistics was applied to all 22 libraries to reduce differences introduced by filtering in the KD–WT comparisons. For each barcode, a custom script (bam_qc_summary, using pysam [74]) summarized mapped reads using three features: the mean soft-clipped fraction (the proportion of read bases left unaligned), its standard deviation, and the mean base quality divided by 10. To limit the influence of noisy estimates at low coverage, scoring was restricted to barcodes with more than a library-specific minimum number of reads, *_n_*_min_ (50–3,000 reads, depending on sequencing depth; **Table S17**), and fewer than one mismatch per read on average. A three-dimensional Gaussian kernel density estimate was fitted to these gated barcodes using SciPy [75]. The quality score *_qi_* for barcode *_i_* was calculated as

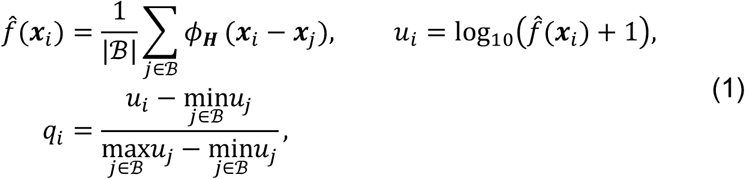

where *_f_*^’^(***x****_i_*) is the density around the feature vector ***x****_i_* of barcode *i*, *u_i_* is the log-scaled density, and *φ**_H_*** is a Gaussian kernel with a data-derived bandwidth matrix ***H***. The set ℬ contains the gated barcodes from the same library and defines the range used for scaling. Thus, *q_i_* → 1 in the dense cell core and *q_i_* → 0 in the diffuse ambient cloud. The score describes how typical a barcode is within its library and provides a common scale on which to apply the same threshold across libraries. A two-component Gaussian mixture model was fitted separately to each library using log_10_ UMI count and *q_i_* (scikit-learn [76]). Barcodes in the higher-count component with more than 100 UMIs were designated candidate cells, and candidates with *q_i_* > 0.9 were retained. Excluded candidates with 0.8 < *q_i_* ≤ 0.9 occupied the same cell-cycle manifold as retained cells but were enriched in early G1. Cell-cycle density calls remained stable across quality-score thresholds from 0.90 to 0.975 (**Fig. S6B-C**).

**Doublet removal.** Doublets were identified from the raw counts of retained cells using Scrublet [77] and scDblFinder [78], each with its automatic score threshold. Scrublet scores cells by the proportion of simulated doublets among their neighbors and was run with an expected doublet rate of 0.1; scDblFinder uses a gradient-boosted classifier. Cells were retained as singlets only if neither method classified them as doublets. Of 25,423,881 raw barcodes, 483,050 were candidate cells, 249,877 passed the quality threshold, and 211,052 were retained as singlets. Filtering selected barcodes without modifying their counts. No ambient-RNA correction, organelle-fraction filter or upper UMI cutoff was applied (**Fig. S6D**).

**Normalization.** Counts were normalized to 10,000 per cell, log1p-transformed and scaled without centering using the per-gene standard deviation calculated across cells with *q_i_* > 0.8. Processing was performed in Python and R [79] using sclab [80] and scanpy [81]. Software versions, including supporting R and Python packages [82–106], are listed in **Table S15**, and per-library values are reported in **Table S2**, **Table S17** and **Table S18**.

#### Cell-cycle clock

**Reference clock.** Asynchronously dividing tachyzoites collectively span the cell cycle, forming a closed loop in gene-expression space. A single clock was constructed from the WT loop and used to assign KD cells to matched cell-cycle positions. Independently fitted clocks would not provide directly comparable coordinates across libraries and could be unreliable when a knockdown distorts the cycle. The 6,985 WT singlets were normalized, scaled and projected by principal component analysis using all genes detected in WT. Genes were not preselected for variability, allowing genes that change along the cycle to remain in the analysis. Cells were initially ordered by their polar angle in the plane of the first two principal components, and a closed periodic smoothing spline was fitted to each of the first 20 components. For each component, the signal-to-noise ratio (SNR) is the variance of the fitted curve under this initial ordering divided by the component’s total variance. These ratios were scaled to the largest SNR. Components with a scaled SNR above 0.75 (components 1–4) were retained as the cell-cycle signal, and the closed curve ***γ***(*s*) through these components defines the clock. Each cell *i* was projected onto the closest point *s_i_* on the curve, and its distance to the curve was recorded. Pseudotime *τ_i_* is the normalized arc length up to that point, so that equal pseudotime increments represent equal distances along the expression trajectory:

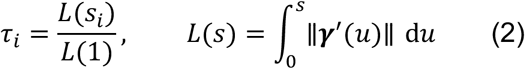

Pseudotime runs from 0 to 1 and is periodic. Because a closed curve has no natural start or orientation, the origin was placed at an inflection point of the reciprocal cell density along the curve next to the densest stretch of the cycle, between C and G1a phases and correcting the sign direction if necessary. Pseudotime was converted to hours with the cumulative density of the WT cells:

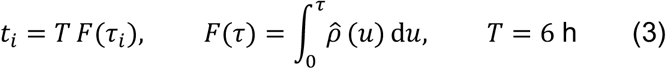

where *ρ*U is a circular kernel density estimate of WT pseudotimes that integrates to 1, and *T* is the fixed cycle length. This conversion assumes constant cell flux through the cycle and a uniform age distribution, such that the proportion of cells at a given position reflects the time spent there. Equal time intervals therefore contain equal numbers of cells. The first 30% of the arc, containing the two dense G1 regions and 64% of WT cells, spans the first 3.8 h; the remaining 70% spans 2.2 h. An exponential-growth age distribution would shift the assigned times by up to approximately 0.5 h, primarily in mid-cycle. The clock was computed using sclab (**Fig. S7A-B**).

**Phase labels.** Cell-cycle phases (G1a, G1b, S, M and C) were transferred to WT cells from a Smart-seq2 dataset of 595 tachyzoites (GSE145080), in which Xue et al. annotated phases using unsupervised clustering and DNA content [11]. The transfer process used the joint embedding and weighted vote described below, with the Smart-seq2 cells as the reference. Any WT cells that remained unlabeled were assigned labels by propagation over the WT neighbor graph. The transferred phases followed the sequence G1a, G1b, S, M and C along the assigned time axis, with median times of 0.85, 2.64, 4.29, 5.23 and 5.75 h, respectively (**Fig. S7C**).

**Transfer features.** Periodic genes were identified in WT cells by cosinor regression to select features carrying the cell-cycle signal for mapping knockdowns [107–109]. The cycle was divided into 10 equal circular pseudotime bins. Counts were summed within each bin, normalized and log-transformed, and each gene’s ten-point profile was fitted with a single sinusoid with a period of one cycle. An *F* test of zero amplitude with 2 and 7 degrees of freedom, followed by Benjamini–Hochberg correction [110], identified 5,377 of 7,536 genes as periodic at q ≤ 0.05. The transfer features are the union of these genes and 949 cell-cycle phase-marker genes (5,426 genes). A more stringent set of 3,929 periodic genes (q < 0.01) was used to construct the shared embedding described below (**Fig. S7D**).

**Transfer of time and phase to knockdowns.** Each KD library was mapped pairwise to the WT reference. The two libraries were restricted to the transfer features, normalized and scaled separately to unit variance without centering, and embedded jointly by canonical correlation analysis (CCA; 50 components [111]). The leading components were dominated by shared variation, primarily the cell cycle. In the first 10 components, a batch-balanced k-nearest-neighbor graph (BBKNN [112]) linked cells to their 20 nearest WT cells and 20 nearest cells of the same library by cosine distance, with UMAP-style connectivities *w_i_*_j_ as edge weights. Each KD cell *i* received a phase, pseudotime and time from its WT neighbors N*_i_*, weighted by *w_i_*_j_. The assigned phase was the phase with the greatest total neighbor weight. For time transfer, neighbor times were shifted relative to their median before averaging to account for the periodic boundary at 0/6 h. This avoided averaging neighboring cells on opposite sides of the boundary into mid-cycle. The time error *e_i_* is the weighted standard error of the transferred mean:

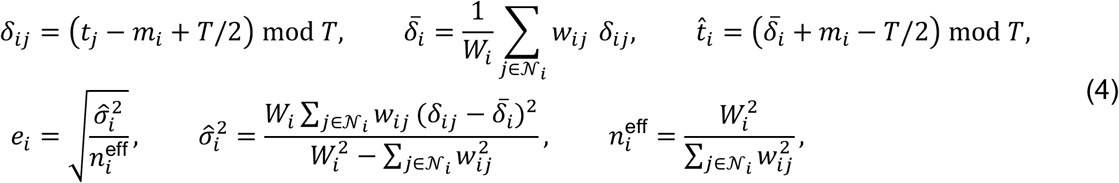

where *t*_j_ is the time of WT neighbor *j*, *m_i_* is the median neighbor time, and *δ_i_*_j_ is the shifted time, with the median positioned at mid-period. The total neighbor weight is *W_i_* = ∑_j∈N_*_i_ w_i_*_j_, ^*_i_* is the transferred time, *σi*^2^ is the unbiased weighted variance of the shifted times, and *ni*^eff^ is the effective number of neighbors. The error *e_i_* approaches 0 h when WT neighbors agree on time and increases as their times become more dispersed. Pseudotime was transferred by the same procedure with a period of 1. Cells with fewer than 15 WT neighbors were left unassigned (214 of 211,052 singlets), and cells with *e_i_* < 0.25 h were retained for downstream analysis (198,284 of 211,052 singlets; **Fig. S7E**).

**Validation of the transfer.** Transfer precision was assessed by mapping WT cells onto the reference using the same procedure. Because these cells were also present in the reference, this control represents best-case precision. The mean circular absolute difference between transferred and reference times was 0.061 h, phase labels agreed for 91.1% of cells, and 6,924 of 6,985 WT cells (99.1%) passed the time-error filter. Across the 21 KD libraries, 191,360 of 204,067 singlets passed, with per-library retention ranging from 85.4% for HDAC3 to 97.6% for SNIP1. KD cells inherit the clock positions of their nearest WT neighbors, including when they occupy states absent from WT. The time error therefore measures agreement among reference neighbors. The mean distance to those neighbors in the joint space was also recorded to flag KD cells lacking a close WT counterpart. All transferred quantities remain dependent on the WT clock assumptions, including the 6 h cycle, uniform age distribution and externally derived phase labels (**Fig. S7F**).

#### Cell-cycle density and deviation

**Density along time.** Slowing or blocking a cell-cycle stage can produce an accumulation of cells at that stage and a depletion in subsequent stages. Under the residence-time assumption used to construct the clock, these changes can be described by comparing cell densities along the time axis. Density estimates used the transferred times of the 198,284 cells with a time error below 0.25 h. For each library *l*, a von Mises kernel was used to account for the periodic boundary at 6 h:

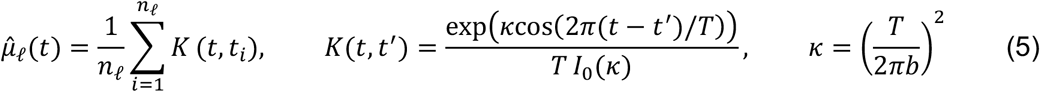

where *t_i_* denotes the times of the *n_l_* cells in the library, *T* = 6 h, *I*_0_ is the modified Bessel function of order zero, and *b* = 0.5 h is the bandwidth (*κ* = 3.65). Densities were evaluated at 513 grid points over one cycle. Each density integrates to 1, so accumulation in one region is balanced by depletion elsewhere. The WT density is approximately uniform at 1/*T* by construction of the time scale. Each density was accompanied by a pointwise band with half-width 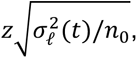, where *σ_l_*^2^(*t*) is the variance of kernel values across cells at time *t*, and *z* = 1.4395 defines an 85% two-sided band. A fixed nominal sample size of *n*_0_ = 2,000 replaced the observed cell count to make band widths comparable across libraries. An interval was classified as an *accumulation* when the KD density exceeded the WT density by more than the sum of their band half-widths, and as a *depletion* when it fell below WT by more than that sum. The time of the greatest KD excess over WT was recorded as the peak if it fell within an accumulation interval; the time of the greatest deficit was recorded as the trough if it fell within a depletion interval. These classifications are descriptive: the bands use a fixed nominal sample size and a single WT reference, are pointwise, and are not corrected for comparisons across the 513 grid times. The intervals were not subjected to formal hypothesis testing (**Fig. S18A-B**).

**Shared embedding.** To compare all knockdowns with WT in a common coordinate system, the 22 libraries were jointly embedded using scVI [37] in scvi-tools [113]. The model was trained on raw counts with library identity as the batch covariate. Input genes comprised the union of WT highly variable genes, WT periodic genes at q < 0.01, and genes differentially expressed in at least one KD, restricted to genes detected in all libraries. Including differentially expressed genes retained perturbation-related information in the input. For this selection, a gene was considered differentially expressed if, in at least one time window of the limma contrasts (Section “Differential expression within time windows”), it had an adjusted P value below 0.05, an absolute robust z score [114] of log fold change above 2.576 within the window, and detection in more than 5% of cells. The model used two hidden layers of 128 units, a dropout rate of 0.2, a 10-dimensional latent space and a zero-inflated negative binomial likelihood. For early stopping, 10% of cells were held out to monitor validation loss. Each cell was represented by the posterior mean of its latent vector, and a two-dimensional UMAP [115] was calculated for visualization. Because each KD was represented by a single library and library identity was the batch covariate, library and knockdown effects are confounded in this embedding.

**Deviation from WT.** Density captures changes in the distribution of cells along the cycle. To describe differences in transcriptional state at matched cycle positions, each cell was also compared with the expected WT state at its transferred time. The deviation of cell *i* is *δ_i_* = ‖***z****_i_* − ***f***_WT_(*t_i_*)‖_2_, where ***z****_i_* contains the cell’s latent coordinates and ***f***_WT_(*t_i_*) is the expected WT state. A cell lying on the WT trajectory therefore has *δ_i_* = 0. The reference trajectory ***f***_WT_ was constructed by smoothing WT latent coordinates along time with a centered 0.5 h rolling mean, wrapping across the 6 h boundary, and fitting a periodic cubic spline in each latent dimension. Deviations were calculated for cells included in the density analysis that also had coordinates in the shared embedding. They were displayed against the WT deviations, summarized by a rolling mean ± one standard deviation over 1 h. This baseline is likely biased downward because WT cells were evaluated against a trajectory fitted to the same cells. Deviations are distances within a single learned latent space and were therefore compared only among libraries represented by the same model. Errors in transferred time can also inflate deviation where expression changes rapidly along the trajectory. The deviation measure was consequently treated as descriptive (**Fig. S18C**).

#### Differential expression within time windows

**Time windows.** Knockdowns can alter both gene expression and the distribution of cells along the cycle (Section “Cell-cycle density and deviation”). These effects are confounded in an unstratified comparison: an accumulation of cells in one phase can appear as induction of genes expressed in that phase. KD and WT cells were therefore compared within matched time windows. The 0–6 h cycle was divided into 24 segments of 15 min, with each set of four consecutive segments defining a 1 h window that wrapped at 6 h. The resulting 24 windows were labeled by their centers, from 0 to 5.75 h (**Table S19**), and grouped into four sets offset by 15 min. Each set tiled the cycle with six nonoverlapping windows, placing each cell in one window per set and four windows overall. Because window width was fixed, cell numbers varied across windows and libraries. Retained cells had an estimated time standard error below 15 min, one-quarter of the window width (**Fig. S19A**).

**Pseudo-replicates.** Each of the 21 KD factors was compared with the single WT library using raw counts from cells with a time error below 0.25 h (Section “Cell-cycle clock”). Comparisons comprised 24 window contrasts and one bulk contrast that included all cells (Section “Differentially expressed genes”). Each condition was represented by one library without biological replicates. To provide the multiple observations required by the linear model, cell barcodes within each contrast and condition were shuffled and divided into five sets of approximately equal size. Counts were summed within each set to form a pseudo-replicate (sclab). These pseudo-replicates capture only cell-to-cell sampling variation within a culture. The partitioning was repeated with seeds 0 to 19 to assess sensitivity to the arbitrary assignment of cells to pseudo-replicates. The seeds changed only this assignment; repeated partitioning did not provide biological replication. Pooled result files, seeds and cell numbers are listed in **Table S20** (**Fig. S19B**).

**Model and pooling.** Lowly expressed genes were removed with the edgeR expression filter [116], applied to the pseudobulk matrix and design of each fit. Library sizes were normalized by the trimmed mean of M values (TMM) [117]. Counts were then transformed to log counts per million and assigned precision weights with voom [118]. Models were fitted with limma [119] in R, using the design ∼ 0 + group + replicate (group = condition × window) and empirical-Bayes moderation of residual variances. The six windows in each set were fitted jointly, sharing one residual variance per gene (60 samples, 44 residual degrees of freedom). The bulk contrast was fitted separately (10 samples, 4 residual degrees of freedom), and each window yielded one KD – WT contrast. The fits from the *m* = 20 partitions were treated as 20 imputations of the cell assignment and pooled for each gene and contrast using Rubin’s rules [120]:

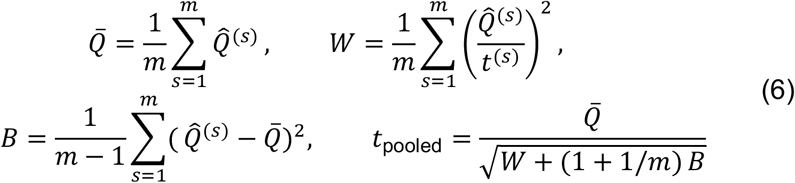

Here, *Q*^|(^*^s^*) and *t*^(*s*)^ are the log2 fold change and moderated t statistic for partition *s*, *Q̄* is the pooled log2 fold change, *W* is the mean squared moderated standard error, representing sampling variation within a partition, and *B* is the variance of the log2 fold change across partitions. If all partitions yield the same estimate, *B* is zero and the pooled variance reduces to *W*. Gene-level P values were calculated from *t*_pooled_ using the small-sample Barnard–Rubin degrees of freedom [121], which decrease as the contribution of between-partition variance increases. Because expression filtering was performed separately for each fit, only genes tested in all 20 partitions were retained. P values were adjusted for the false-discovery rate (FDR) across genes within each contrast using the Benjamini–Hochberg procedure [110]. With one library per condition, the pseudo-replicates capture only cell-to-cell sampling variance; consequently, the P values and FDR estimates overstate significance. All tested genes were retained in the result tables. Criteria for differentially expressed genes (DEGs) are given in Section “Differentially expressed genes” (**Fig. S19B**).

#### Differentially expressed genes

**Time-course DEGs.** A gene tested in a window is classified as a time-course DEG for a factor when FDR < 0.05, |*Q̄* | > 1 (a twofold change) and *π* > 0.05, where *π* = max(*π*_KD_, *π*_WT_) and *π_c_* is the fraction of cells in condition *c* within that window with a nonzero count. Direction was determined by the sign of *Q̄* : increased expression in the knockdown (*Q̄* > 0) was interpreted as repression by the factor, and decreased expression (*Q̄* < 0) was interpreted as activation. For each factor, time-course DEGs were summarized by window and direction, as the union across all 24 windows, and by cell-cycle phase. In the union, a gene that increased in some windows and decreased in others contributed to both directions. For the phase summary, windows were assigned to the phase containing their center, with M and C pooled. Each phase included genes called in any of its windows; when a gene had multiple calls, its direction was assigned from the call with the largest |*Q̄* |. These summaries are descriptive because windows overlap and FDR was controlled only within individual contrasts (**Fig. S19C**).

**Bulk contrast.** The bulk contrast included all time-filtered cells from the two libraries and measured the overall response to each knockdown, including changes in cell-cycle composition. Bulk DEGs were identified using the time-course criteria, with *π* calculated across all time-filtered cells in each condition. These calls were used to summarize per-factor DEG counts, distinguish direct from indirect targets and calculate direction balance (Section “Enrichment analysis”, **Table S21**).

**Use in the networks.** CUT&RUN binding was measured once per factor, so the window contrasts provide all temporal resolution in the networks. An edge from factor *f* to gene *g* in window *w* requires that *g* was tested in *w* and passed a differential-expression gate in the knockdown of *f* (Section “Functional-network construction”). This gate fixes FDR ≤ 0.05 and |*t*_pooled_| ≥ 2 while varying the thresholds on |*Q̄* | and *π* (Section “Parameter sweep”). The resulting networks therefore span a family of DEG definitions around the criteria above. A separate selection, based on the robust z score of the log2 fold change within each window, supplied genes for the shared embedding (Section “Cell-cycle density and deviation”; **Fig. S19D**).

#### Velocity along pseudotime (PfCdc5)

**Velocity along pseudotime (*Pf*Cdc5).** A single-cell RNA-seq library from the knockdown of *Pf*Cdc5, the *P. falciparum* ortholog of Cdc5 (CEF1), was quantified with Cell Ranger count [73] against the *P. falciparum* 3D7 genome and annotation from PlasmoDB release 68 (5,720 genes). Cells were selected from the raw feature-barcode matrix using the per-barcode alignment statistics described in Section “Single-cell RNA-seq data processing”. Here, thresholds were applied to robust z-scores [114], defined as the distance from the median in units of the scaled median absolute deviation. Barcodes with low read counts, low base or mapping quality, high mismatch counts or an atypical soft-clipped fraction were excluded. Requiring at least 50 UMIs and 5 genes per cell retained 5,995 cells. Counts were normalized, log1p-transformed and scaled as for *T. gondii*. Cells were analyzed using PCA, a cosine nearest-neighbor graph, UMAP [115] and Leiden clustering [122]. Parasite stages were assigned with sc-type [123] using marker genes identified by differential expression between annotated stages in the public Malaria Cell Atlas [124] (**Table S22**). Removal of two clusters of gametocytes and outlying ring-stage cells left 5,339 cells. Periodic pseudotime along the intraerythrocytic cycle was then fitted using the closed-curve procedure described for the WT clock (Section “Cell-cycle clock”). The initial order was given by each cell’s angle on an ellipse fitted to the first four principal components, and components 1–3 captured the cycle signal. Cells more than three to five robust standard deviations from the median trajectory in any component were excluded, leaving 4,416 cells. A UMAP of these cells was used for display.

**RNA velocity and display.** Spliced and unspliced counts were obtained from the Cell Ranger alignments with velocyto [125]. Two closed guide paths, representing an outer and an inner loop through the cycle, were drawn on the UMAP using the guided-pseudotime tool in sclab. Each cell was assigned a position by projection onto each path. RNA velocity was estimated with scVelo [126] separately for two subsets: an outer subset containing all cells except gametocytes and one Leiden cluster labeled perturbed (3,805 cells), and an inner subset containing non-gametocyte cells within 4 UMAP units of the inner path, including the perturbed cluster (2,980 cells). Within each subset, genes with fewer than 20 counts shared between the spliced and unspliced layers were removed, and both layers were normalized per cell. First-order moments were computed over 30 nearest neighbors in 30 principal components, and the stochastic model was fitted, yielding 171 and 135 velocity genes. Velocities were projected onto the UMAP through the velocity graph. For display, projected vectors from each subset were averaged on a regular grid using a Gaussian kernel with a bandwidth of 0.8 UMAP units, truncated at 2.4 units. The averaged field was integrated into streamlines within 1.05 units of a smoothed guide curve, and grid points distant from all cells were masked. Fixed-length arrows placed every 1.9 units along each guide indicated the direction of the kernel-averaged velocity. The display included 4,076 cells, of which 2,709 belonged to both subsets.

#### Splicing efficiency

Spliced and unspliced transcript counts were generated per cell with velocyto and imported from the resulting loom files (hdf5r [127]). For each sample the spliced matrix was used to build a Seurat [128,129] object and the unspliced matrix added as a second assay. Cells were filtered on detected genes (*T. gondii*: 200–3000 for RH, 100–3000 for Cdc5-cKD and SNIP1-cKD; *P. falciparum*: 100–3000 for WT and control, 100–2700 for Cdc5-cKD), and analyses restricted to genes detected in all samples of each species. Counts were log-normalised, the 2000 most variable genes selected by variance-stabilising transformation, scaled, and principal components computed (30 PCs); UMAP embeddings were generated with uwot on the first 10 PCs (n_neighbors = 30, cosine metric, min_dist = 0.3). Cell-cycle position was assigned differently in the two species: *T. gondii* cells were labelled (G1.a, G1.b, S, M, C) by anchor-based label transfer from a published ME49 reference dataset, whereas *P. falciparum* cells carried intraerythrocytic developmental cycle stage labels (ring, trophozoite, schizont) assigned independently and matched to cells by barcode.

Splicing efficiency was quantified in two ways. The global per-cell spliced ratio was defined as total spliced counts divided by the sum of spliced and unspliced counts across all genes in that cell. For the intron-resolved analysis, genes were binned by exon number taken from the genome annotation (PlasmoDB-68 *P. falciparum 3D7*) into exclusive bins of exactly 1, 2, 3, 4, 5 and more than 5 exons, corresponding to 0, 1, 2, 3, 4 and more than 4 introns; for each cell and bin the spliced ratio was computed per gene and averaged across genes in that bin, with genes lacking counts in a cell excluded. Intron-deprived (single-exon) genes have a spliced ratio of 1 by definition and were excluded from the intron-number comparisons where indicated. Splicing efficiency was compared between knockdown and control cells globally, within each cell-cycle stage and within each intron bin. Comparisons of per-cell values used a linear model with estimated marginal means (emmeans), which accommodates unequal cell numbers; binned comparisons, in which each cell contributes one value per bin, used a linear mixed-effects model with a random intercept per cell (lmerTest). Ratios were scaled ×100 before fitting for numerical stability and effect sizes back-transformed. Equivalent Wilcoxon rank-sum tests gave the same conclusions. Significance is shown as ***P < 0.001, **P < 0.01, *P < 0.05. Boxplots show median, interquartile range and 1.5 × IQR whiskers; cell and gene numbers per group are given on each panel.

#### Single-cell ATAC-seq processing

**scATAC-seq processing.** One 10x Genomics single-cell ATAC-seq library was processed with Cell Ranger ATAC [130] against the ToxoDB release 68 ME49 genome and annotation. From the unfiltered fragment file, 8,925 cell barcodes were retained, with 4,640 to 56,377 fragments per barcode (median 9,973).

**Promoter accessibility.** Fragments were counted per cell with Signac [131] within a strand-aware 2 kb promoter window centered on the annotated 5′ gene boundary (±1 kb) of each of the 8,178 genes on the 14 nuclear chromosomes. These windows were also used for CUT&RUN peak-gene assignment. Phase and time were transferred from an earlier version of the WT scRNA-seq clock (16,810 cells) to the scATAC cells through a WT single-cell multiome dataset (10x Genomics), in which RNA expression and chromatin accessibility were measured in the same cells. First, phase and time were transferred from the WT scRNA-seq cells to the multiome cells using the RNA measurements, integrated by CCA (30 components) and a BBKNN graph (20 components, 25 neighbors per batch). They were then transferred from the multiome cells to the scATAC cells using promoter gene-activity matrices integrated by CCA (30 components) and a BBKNN graph (20 components, 25 neighbors per dataset). The time error is defined as for the knockdowns (Section “Cell-cycle clock”). Cells with a time error < 0.25 h were retained (3,526 of 8,925; **Fig. S20A**).

#### Expression and accessibility profiles

**Normalization and imputation.** Gene clustering, community profiles and motif timing analyses require smooth gene-level functions of time. Each cell is measured once, and most counts are zero. Continuous WT mRNA expression and promoter accessibility profiles were therefore constructed over the 0–6 h cycle using a common procedure. Expression profiles were derived from counts in the WT reference cells, whose assigned times are uniformly distributed over the cycle by construction (Section “Cell-cycle clock”). Accessibility profiles were derived from promoter-window counts in the 3,526 scATAC cells with a time error below 0.25 h (Section “Single-cell ATAC-seq processing”), whose assigned times are concentrated in S, M and C. Genes with fewer than 30 counts across these cells were excluded. Counts were normalized to 10,000 per cell, log1p-transformed and scaled to unit variance without centering, preserving zero values for cells with no count. The same treatment was applied to the scATAC counts, without term-frequency or latent-semantic transformation. Each matrix was then imputed with ALRA [132] to fill technical zeros, because window means of unimputed values reflect both expression level and the fraction of cells in which a gene is detected (**Fig. S20B**).

**Spline fit and scaling.** For each gene, imputed values were averaged across cells within one grid step (0.047 h) of each of 128 equally spaced points on the periodic time grid. Thus, each cell contributed to approximately two windows, and windows near 0 h included cells near 6 h. A periodic cubic smoothing spline was fitted to these averages after rescaling time to a unit period (sclab NDBSpline, penalty *λ* = 7.8 × 10^−5^). Three copies of the cycle were fitted, and the profile was extracted from the central copy to ensure smooth closure at 0 h = 6 h. The fitted function *F*_g_(*t*) was evaluated at 257 equally spaced times from 0 to 6 h; because the final point repeats the first, one cycle was represented by 256 distinct points. Each profile was divided by max(2*σ*_g_, *m*_g_), where *σ*_g_ is the standard deviation of the fit residuals across cells and *m*_g_ is the maximum fitted value at the cells’ times. The scaled profile therefore peaks at 1 when 2*σ*_g_ ≤ *m*_g_ and at *m*_g_/(2*σ*_g_) otherwise, reducing the amplitude of noisy profiles. Genes excluded by the count filter and genes with constant fitted profiles were treated as missing. Genes on unplaced scaffolds have no promoter window and therefore no accessibility profile (**Fig. S20B**).

**Use.** The scaled profiles define the expression (E) and accessibility (A) layers used for gene clustering (Section “Ensemble clustering of genes”). These layers center each profile and scale it to unit length, retaining only its shape. Scaling by max(2*σ*_g_, *m*_g_) affects analyses that average profiles, such as community summaries (Section “Gene communities”), by reducing the contribution of genes with noisy fits. The profiles also provide the timing measures used in the motif analyses (Section “Palindromic motifs and lytic-cycle timing”). Genes lacking a profile retained their network edges but were excluded from clustering (**Fig. S20C**).

#### CUT&RUN data processing

**Processing.** Library preparation and sequencing are described in Section “CUT&RUN”. Pairedend reads were checked with FastQC [133] and trimmed with cutadapt [134] to remove adapters and low-quality 3′ ends (Phred < 25). Pairs were discarded if either read was shorter than 30 bp. Reads were aligned to the ToxoDB release 68 ME49 genome with BWA-MEM [135] at default settings and sorted with SAMtools [136,137]. Properly paired primary alignments on the 14 nuclear chromosomes were retained without a mapping-quality filter. Duplicates were marked with Picard [138] and removed before peak calling, and fragments were extracted with bedtools [139]. Software versions, including supporting tools [140–144], are listed in **Table S15**.

#### CUT&RUN peak analysis

**Peak calling and library selection.** Peaks were called for each library with MACS2 [145] in paired-end mode, using duplicate-removed alignments and a shared IgG CUT&RUN control. Calls used --keep-dup all and an effective genome size of 62,483,827 bp, corresponding to the 14 *T. gondii* chromosomes. MACS2 compares fragment pileup with the local background and reports the fold enrichment at each peak summit and a Benjamini–Hochberg q value. Peak sets at five q-value thresholds (0.025, 0.05, 0.075, 0.10 and 0.125) were used to construct the networks; figure panels use the q = 0.075 set (q < 0.075). One library per factor was selected by examining unique fragments, signal-to-noise, fraction of reads in peaks (FRiP) and peak yield relative to the IgG control. This yielded one library for each of the 21 factors represented in the knockdown dataset (**Table S23**, **Table S24**, **Table S25**; **Fig. S21A**).

#### Peak-gene assignment

Peaks were assigned to genes using 2 kb promoter windows centered on the annotated 5′ gene boundary (±1 kb). A peak was assigned to every gene whose window it overlapped by at least 1 bp; broad peaks could therefore be assigned to multiple promoters. Peaks outside these windows and genes on unplaced scaffolds received no assignment. For each gene, the overlapping peak with the highest MACS2 fold enrichment, *x*, was retained (48,615 factor–gene pairs across 6,888 genes at *q* = 0.075). Within each peak file, *x*à and MAD denote the median and median absolute deviation of *x* across assigned genes. Each value of *x* was converted to a robust Z score *z* and a normalized signal *s*:

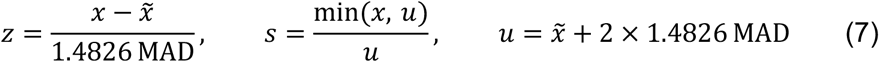

Here, 1.4826 MAD estimates the standard deviation while limiting the influence of the strongest peaks, and the cap *u* is two such units above the median. The normalized signal increases linearly with *x* up to this cap and equals 1 above it, preventing a few strong peaks from dominating a factor’s weights. The Z score was used only to trim low-signal peaks in the edge rule, and the normalized signal contributed to the edge weight. Both quantities were recalculated for each peak file, so edge weights rank targets within a factor and remain on factor-specific scales (**Fig. S21B-C**).

#### Functional-network construction

**Edge rule.** Knockdown responses can reflect direct or indirect regulation, and binding peaks can occur with or without a regulatory effect. The networks combined these two sources of evidence using an explicit rule, without fitting a model or assigning P values to edges. An edge from factor *f* to gene *g* in window *w* was included if and only if *g* passed a differential-expression gate in the knockdown of *f* during *w* and had a promoter peak for *f* that passed a CUT&RUN gate. The differential-expression gate required that *g* was tested in *w*, FDR ≤ 0.05, |*t*_pooled_| ≥ 2, |*Q̄* | ≥ *L* and *π* ≥ *P*, where *π* is the larger fraction of cells with a nonzero count across the two conditions. Setting *L* = 0 imposed no effect-size floor. The |*t*_pooled_| criterion was never limiting because all genes with FDR ≤ 0.05 had |*t*_pooled_| ≥ 2.09. Each gene was evaluated only against the knockdown of *f*. The CUT&RUN gate required *z* > Φ⁻¹(1 − *c*/100): *c* = 95 excluded peaks with *z* ≤ −1.645, and *c* = 100 retained all peaks. The four thresholds controlled peak-calling stringency (*q*), trimming of weak peaks (*c*), the effect-size floor (*L*) and the detection floor (*P*); their grid is defined in Section “Parameter sweep”. At each setting (*q*, *L*, *P*, *c*), the network comprises all (window, factor, gene) triples passing both gates, weighted by the signed expression response scaled by the binding signal:

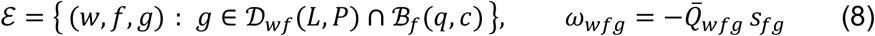

Here, D_w*f*_(*L*, *P*) and ℬ*_f_*(*q*, *c*) are the genes passing the differential-expression and CUT&RUN gates for factor *f* in window *w*. Genes whose expression decreased in the knockdown (*Q̄*_w*f*g_ < 0) received positive weights, indicating activation; genes whose expression increased received negative weights, indicating repression. Weight magnitude increased with the knockdown effect and binding signal, up to the cap on the latter. Each network resolved regulation across the 24 windows described in Section “Differential expression within time windows”. All peaks within a promoter window were treated equally regardless of their distance from the annotated 5′ gene boundary. An edge indicates that the factor binds a gene’s promoter and that the gene responds to its knockdown within a window. This evidence does not exclude an indirect response mediated by another bound target. Similarly, an absent edge does not exclude regulation: a gene may be untested in a window, bound outside the defined promoter window or just below a threshold. Promoter accessibility did not enter the edge rule; it contributed to gene clustering (Section “Gene communities”; **Fig. S21D-E**).

#### Ensemble clustering of genes

**Gene layers.** Target genes were clustered using three layers. The regulatory layer (R) represents each gene’s regulatory fingerprint: its edge weights across all (window, factor) rows of a network, with 0 for absent edges. Activation and repression were placed in separate halves to prevent opposite regulatory effects from canceling; the resulting vector was square-root transformed and scaled to unit length. Genes with no shared nonzero coordinate, including genes regulated in opposite directions by the same factors, have a regulatory similarity of 0. A zero coordinate indicates only that no edge was called and does not exclude regulation. The expression (E) and accessibility (A) layers were derived from the scaled WT profiles (Section “Expression and accessibility profiles”). Each profile and its periodic derivative were centered, scaled to unit length and combined with weights of Ä1/2. These layers capture temporal shape, with the inner product of two gene vectors equal to the average correlation of their profiles and their derivatives. E and A are identical across networks; only R varies. Clustering included only genes with all three layers, excluding genes with flat profiles that lacked an E or A layer (**Fig. S22A**).

**Fused distance.** For layer *l*, the distance between genes *g* and ℎ is the angle between their unit vectors divided by the maximum angle permitted by that layer:

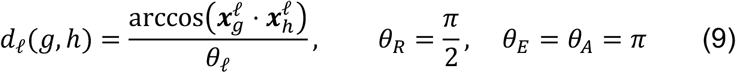

This maximum is *π*/2 for the nonnegative R vectors and *π* for the centered E and A vectors, placing all three distances on a 0–1 scale. The normalized angular distance satisfies the triangle inequality, which one minus the cosine does not. The fused distance is the unweighted mean, *d* = (*d_R_* + *d_E_* + *d_A_*)/3. Gene pairs without a shared regulatory coordinate have *d_R_* = 1, so their fused distances differ only through E and A (**Fig. S22B-C**).

**Ensemble and consensus cut.** To assess sensitivity to gene inclusion and uncertainty near cluster boundaries, clustering was repeated on subsamples. In each of 120 replicates, a random 80% of genes were sampled without replacement and clustered by average linkage on *d*. The resulting tree was partitioned using the hybrid dynamic tree cut [50] (deep split 1, PAM stage on, minimum module size 10), which uses branch shape to determine splits. Genes left unassigned by this cut were treated as singletons for that replicate.

The co-association *a*(*g*, ℎ) is the fraction of replicates containing both genes in which they belonged to the same cluster. A consensus tree was constructed by average linkage on 1 − *a* and cut at height 0.7, allowing groups to merge only when their mean co-association was at least 0.3. A gene was committed to a community if its mean co-association with the other group members, termed its support, was at least 0.3 and the group retained at least 10 genes. Smaller groups were dissolved, and committed genes retained their groups without subsequent merging. Remaining genes were left unassigned, a clustering outcome with no inherent biological interpretation (**Fig. S22D-E**).

All settings were fixed in advance and were independent of network-selection objectives and enrichment results. Subsamples for all replicates were drawn using one random-number generator initialized once with seed 0.

**Use.** Clustering was applied to all 630 networks in the parameter sweep and contributed to two scoring objectives, reach and stability (Section “Parameter sweep”). It was then applied to the consensus network to define gene communities (Section “Gene communities”). Variants using R alone or R and E together were used only to assess the sensitivity of the Pareto front.

#### Parameter sweep

**Parameter sweep.** The edge rule requires four thresholds (Section “Functional-network construction”): the peak-call *q*, which selects the MACS2 peak set; the effect-size floor *L*, which sets the minimum knockdown response |*Q̄* | on the log_2_ scale; the detection floor *P*, which sets the minimum detection fraction *π*; and the trim *c*, which determines whether the weakest peaks for each factor are excluded. These choices affect both the genes entering a network and the communities identified downstream.

Networks were constructed for all combinations of *q* (0.025, 0.05, 0.075, 0.10, 0.125), *L* (0 to 1.5 in steps of 0.25), *P* (0.01 to 0.09 in steps of 0.01) and *c* (95 or 100), yielding 5 × 7 × 9 × 2 = 630 networks with all other edge-rule settings fixed. The levels of *L*, *P* and *c* are nested, so a stricter level keeps a subset of the edges of a looser one. The levels of *q* are nested at *c* = 100 and to within 0.02% of edges at *c* = 95, because the trim score *z* is recomputed for each peak file. The equally spaced grid defines the one-step neighbors used to assess stability below. Networks contained 40,724 to 442,326 edges and 2,683 to 5,373 target genes (**Fig. S23A-B**).

An automated Nextflow pipeline [146] constructed each network, clustered its genes and calculated the scoring objectives. Steps were implemented in Python and run as independent jobs on a high-performance computing cluster. Software versions, including supporting packages [147], are listed in **Table S15**.

**Scoring and Pareto front.** Networks were evaluated on three objectives: reach, stability and GO recall. *Reach* is the fraction of the 5,373 target genes observed across all 630 networks that were committed to clusters by a network’s ensemble clustering (Section “Ensemble clustering of genes”). This fixed denominator prevented a network from increasing reach simply by reducing its gene set. *Stability* is the minimum adjusted mutual information [148] between a network’s clusters and those of each network one grid step away in *q*, *L* or *P*, calculated over genes committed in both networks. High stability indicates that small changes in thresholds did not substantially reorganize the clusters (**Fig. S23C**).

*GO recall* measures whether genes with similar regulatory fingerprints share functions. It uses the R layer alone, without clustering, because E and A are identical across networks. GO classes from the ToxoDB release 68 annotation were propagated to ancestor terms and retained if they contained 5 to 200 network genes, with all three namespaces pooled. For each query gene and class, the Mann–Whitney area under the curve [149] measures the probability that a class member is closer to the query in fingerprint distance than a nonmember; the query was excluded from the class. The area was compared with a coverage-matched null to account for the tendency of genes with many edges to lie close to other genes [150]. Null classes contained the same number of members as the observed class within each of five strata of fingerprint coverage. One-sided P values were adjusted for each query gene by the Benjamini–Hochberg procedure [110]. GO recall is the fraction of known gene–class pairs with FDR ≤ 0.05. Because this objective contributed to network selection, subsequent GO enrichment of communities was interpreted as a consistency check (**Fig. S23D**).

The three objectives were maximized jointly. The Pareto front comprises networks for which no other network performed at least as well on every objective and better on at least one. Exact comparisons, without a tolerance, identified 61 front members (**Table S26**). These included all five peak-call *q* values, all nine *P* values and both values of *c* (34 at *c* = 95 and 27 at *c* = 100). They included only *L* ≤ 0.5: 25 at *L* = 0, 35 at *L* = 0.25 and one at *L* = 0.5. Single-network examples used the *representative setting q* = 0.075, *L* = 0.25, *P* = 0.05 and *c* = 95 . This front member contained 311,905 edges from 20 factors to 4,750 genes; HDAC5 had no edges at this setting (**Fig. S23E-F**).

#### Consensus networks

**Consensus network.** The 61 Pareto-front members were combined by voting. An edge *e* = (*w*, *f*, *g*) was accepted if it appeared in at least half of the members, with its weight calculated as the mean across members containing that edge:

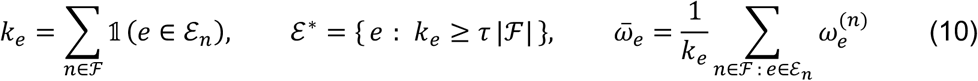

Here, ℱ is the Pareto front, ℰ*_n_* is the edge set of member *n*, *k_e_* counts edge presence regardless of sign and *τ* = 0.5 is the consensus threshold, requiring *k_e_* ≥ 31 of 61. Candidate edges were the union of all member edge sets: 442,326 edges across 5,373 genes, matching the most permissive member. Support was calculated over the entire front, so a member unable to contain an edge still counted against its inclusion. Weights were averaged only over members containing the edge to avoid dilution by absent edges. No quantities were reestimated, and no edge changed sign across members because all networks used the same differential-expression tables. The 660 of 5,373 genes without an accepted edge were removed, and all 21 factors retained edges. The resulting consensus network contained 308,098 time-resolved edges from 21 factors to 4,713 genes (**Table S27**). Votes were not independent because thresholds were nested and the trim levels produced nearly identical networks. Support therefore measures how broadly an edge was retained across the parameter space represented by the front. Sensitivity to the consensus threshold and to the clustering used to define the front is reported in **Table S28**. Consensus calculations were performed in Python (**Fig. S23G**).

#### Gene communities

**Communities of the consensus network.** Ensemble clustering (Section “Ensemble clustering of genes”) was applied to the consensus network. Of its 4,713 target genes, 4,619 had all three layers and were eligible for clustering. The analysis identified 108 communities containing 10 to 181 genes (median 19), numbered in decreasing order of size. Together, these communities contained 3,838 genes (83.1% of those clustered), with member support ranging from 0.30 to 0.86 (median 0.56). Of the remaining 781 genes, 157 were unassigned because of the support floor and 624 because of the size floor. Each community was summarized by the mean scaled expression and accessibility profiles of its members. On each circular mean profile, peak and trough times are the times of the maximum and minimum, amplitude is their difference, and the half-amplitude level is their midpoint. Rise time is the last upward crossing of this level before the peak, and fall time is the first downward crossing after the peak. The full width at half maximum, calculated as fall time minus rise time modulo 6 h, represents the duration of the community’s active state. Coherence is the mean pairwise Pearson correlation of member profiles across time, approaching 1 for coincident profiles and 0 for unrelated shapes (**Table S29**; **Fig. S22F**).

#### Enrichment analysis

**GO annotation.** GO annotations were obtained from the ToxoDB release 68 *T. gondii* ME49 “Curated” association file [71], with the full file used for a sensitivity analysis. Annotations were mapped to the Gene Ontology release 2026-05-19 [151,152] and propagated through is_a and part_of relations with GOATOOLS [153]. A manually curated set of 45 daughter-budding genes, including 42 of the 4,619 clustered genes (**Table S7**), was added as a custom term.

**Differentially expressed genes.** Per-factor DEGs and their directions are defined in Section “Differentially expressed genes”.

**Community enrichment.** Each of the 108 communities was tested for GO-term enrichment using Fisher’s exact test, with the 1,906 annotated committed genes as the background. Ontology roots and terms represented by fewer than three background genes were excluded. P values were adjusted by the Benjamini–Hochberg procedure [110] across all over-represented community–term pairs. Enriched terms (FDR < 0.05) were reported only when the same community had no enriched descendant term. Enrichment for hyperLOPIT compartments [32], using assignments with TAGM-MCMC [154] joint probability ≥ 0.5, was evaluated by a one-sided hyper-geometric test against the 4,713 network genes.

**Directional tests of regulator activity.** All directional tests used a one-sided hypergeometric test for each direction. For each factor and cell-cycle phase (G1a, G1b, S, M/C), bound genes, defined by a promoter peak at q < 0.075, were tested for over-representation among genes that decreased or increased in that factor’s knockdown (per-phase time-course DEGs; Section “Differentially expressed genes”). The background comprised the 8,178 genes with a promoter window. Bonferroni correction was applied across the 168 tests shown in **Fig. 3C** (21 factors × 4 phases × 2 directions). Each community was also tested within each window for overrepresentation of a factor’s activated targets (positive consensus weights) or repressed targets (negative weights), using the 4,713 network genes as the background. Benjamini–Hochberg adjustment was applied across 108,864 tests. A factor was classified as activating or repressing a community if the corresponding test had FDR < 0.05 in any window; a factor could meet both criteria. In **Fig. 5B**, a community received an activation or repression marker in a window only when at least 70% of significant (factor, mode) tests at the stricter threshold FDR < 10⁻¹⁰ supported that mode. A factor’s direction balance is the percentage of its bound bulk DEGs whose expression decreased in the knockdown. Factors were classified as activators, repressors or dual regulators (**Table S30**, **Table S31**, **Table S32**, **Table S33**).

#### Community identity: Functional Annotator

**Functional Annotator.** Approximately half of the clustered genes lack a GO annotation in the ToxoDB “Curated” file, limiting the interpretation of communities from existing annotations alone. The Functional Annotator Tool assigned candidate functions by guilt by association, based on the tendency of genes with shared regulators and similar expression and accessibility timing to share functions [150]. For each query gene and annotated gene set, the analysis asked whether other members of the set were closer to the query than nonmembers.

Distances were calculated over the 4,619 clustered genes using three combinations of the ensemble-clustering layers (Section “Ensemble clustering of genes”): R alone (*d_R_*), RE ((*d_R_* + *d_E_*)/2) and REA (the fused distance *d*). Functional calls and community compositions used REA, and R and RE provided sensitivity views. The effect size is the tie-aware Mann–Whitney AUC for set members relative to other genes, excluding the query from both groups [149]. This is the same statistic used for GO recall; here, its P value was obtained from a one-sided normal approximation with tie correction. An AUC of 1 indicates that all set members are closer to the query than all nonmembers, and 0.5 indicates no difference (**Fig. S24A**).

For each query, P values were Benjamini–Hochberg-adjusted across a family of 696 GO terms containing 5 to 500 genes (306 biological process, 280 molecular function and 110 cellular component terms). This family was defined before inspecting results, using the Curated annotations propagated to ancestor terms (Section “Enrichment analysis”). A term was called when AUC > 0.5 and FDR < 0.05. Calls were removed if a descendant in the same namespace was also called, and at most three calls were retained per gene, ranked by AUC.

To account for the tendency of genes with many edges to lie close to other genes [150], a predeclared sensitivity analysis computed the AUC within ten strata of fingerprint support, defined by the number of positive R coordinates. Strata were fixed before any gene set was examined, and the stratified AUC was reported alongside the global AUC. A separate specificity test held the gene set fixed and ranked the query against the *B* genes outside the set in the same support stratum, each evaluated as a query against that set. The one-sided empirical P value is *p_s_* = (1 + *k*)/(1 + *B*), where *k* is the number of alternative queries at least as close to the set as the original query; ties therefore count against the query. Specificity P values were reported alongside calls but did not affect the calls or community compositions. hyperLOPIT marker sets [32] (26 classes) and three manually curated gene sets (daughter budding, inner membrane complex and rhoptry) were evaluated as separate families (**Fig. S24B-C**).

**Community identity.** Each community member contributed one gene-equivalent, divided equally among its calls in the REA analysis. Community composition was then expressed as the fraction of members assigned to each term, summing masses across the three GO namespaces. Members with no call (unresolved) or no score (unavailable) remained in the denominator, preserving the contribution of missing annotation (**Fig. S24D**).

Compositional tightness is defined as the sum of squared term fractions, interpreted as the probability that two randomly drawn members share a term. It equals 1 when all members share one term and approaches 0 as mass is distributed across many terms. Because genes with high support or existing annotations are easier to annotate, observed tightness was compared with 1,000 random gene sets of the same size, matched exactly on scorability, support stratum and the presence of any recorded GO annotation. The one-sided empirical P value is (1 + *b*)/(1 + 1,000), where *b* is the number of random sets at least as tight as the observed community (**Fig. S24E**).

Functional calls propose candidate functions supported by the network; genes without calls remain unresolved. GO-based calls and community identities were treated as consistency checks because GO recall contributed to network selection. hyperLOPIT localization and immunofluorescence are independent sources of evidence because neither informed network selection. Community compositions were interpreted alongside temporal profiles (Section “Gene communities”) and enrichments (Section “Enrichment analysis”; **Table S34**, **Table S35**, **Table S36**).

#### Regulator similarity

**Factor fingerprints and closeness.** Regulator partnerships were examined by comparing factors in the consensus network. Within each of the 24 windows, a factor’s fingerprint comprised its consensus edge weights across the 4,619 target genes, with activation and repression held in separate non-negative channels, so that a gene regulated in opposite directions by two factors adds nothing to their similarity rather than canceling other shared targets. As for the gene R layer, edge weights entered as signed square roots. Because both channels are non-negative, the cosine similarity of two fingerprints lies between 0 and 1, and closeness was defined as 1 − 2⁄*π* arccos(cos). Closeness ranges from 0 for factors with no shared targets regulated in the same direction to 1 for identical fingerprints. An integrated closeness pooled all windows in which both factors were present (24 for most pairs, 18 for pairs involving HDAC5) by summing the per-window products before taking the cosine. A pair was scoreable in a window if both factors had accepted edges in it and shared at least 20 accepted targets, and scoreable overall if it shared at least 100 accepted target × window cells. Of the 210 factor pairs, 146 were scoreable overall, and 3,263 of the 5,040 pair × window frames were scoreable (**Fig. S25A**). For example, AAF1 and AP2XI-5 shared 5,092 accepted cells, 90% with the same sign, giving a cosine of 0.547 and a closeness of 0.369.

**Null model and per-window calls.** In each window, observed closeness was compared with 1,000 Curveball randomizations [155] of the factor × gene matrix of accepted edges. Curveball preserves exactly the number of accepted targets of each factor and the number of accepted regulators of each gene. After each randomization, a factor’s observed signed weights were reassigned in random order to its new targets, so each factor kept its weight distribution and its balance of activation and repression. Integrated null values were pooled across windows in the same way as the observed ones. Excess closeness is observed closeness minus the null mean. Because pairs with more shared targets tend to show larger excess closeness, a support residual was also computed: within each window, and separately for the integrated comparison, excess closeness was fitted across scoreable pairs against log(1 + number of shared accepted targets) by local linear regression (LOESS, span 0.75) [156], and the residual is a pair’s excess closeness above this trend. A factor pair was called similar in a window if its closeness was at or above the 99th percentile of its null distribution, at least 0.10 above the null mean, at least 0.30, and had a support residual of at least 0.02 (**Fig. S25B**). Of the 3,263 scoreable frames, 377 met all four criteria. No multiple-testing correction was applied across frames or pairs: the null percentile is one of four fixed thresholds, and the calls are intended as a screen for shared regulatory programs, not as tests with a controlled error rate.

**Active windows and display.** Active windows are those in which a pair was called similar, summarized as a count out of 24 and as the longest run of consecutive active windows around the circular cycle. A pair was persistent if active in at least 16 windows; otherwise it was an episode if its longest run spanned at least two windows and a pulse if it spanned one. Pairs with no active window were inactive, and pairs with no scoreable window were unscoreable. Examples included AAF1–AP2XI-5, persistent in 18 windows, and Cdc5–SNIP1, an episode of three windows at the end and start of the cycle. Separately, a pair belonged to the integrated core if its integrated closeness exceeded all 1,000 null values, was at least 0.20, was at least 0.15 above the null mean, and had a support residual of at least 0.02. Twenty-seven pairs, including AAF1– AP2XI-5, met these criteria (**Fig. S25A,C**). For display, windows were grouped into six consecutive 1-h blocks. A pair was shown in a block if it was called in at least half of the four overlapping 1-h windows whose centers fall in that block, with opacity indicating the fraction of windows called (**Fig. 3E**). These calls describe similarity between regulatory programs and do not by themselves establish physical interaction.

#### Motif discovery, shared motif vocabulary, and promoter scoring

**Discovery sequences and promoter blocks.** The shared motif vocabulary shown in **Fig. 6A** and **Fig. S11A** was built from the 42,387 peaks in the 21 selected CUT&RUN libraries, called against IgG with MACS2 2.2.7.1 at *q* < 0.075 (Section “CUT&RUN peak analysis”). For motif discovery, the 8,178 ToxoDB release-68 strand-oriented promoter windows (annotated 5′ gene boundary ±1 kb) were merged wherever they overlapped, yielding 6,134 non-overlapping blocks: 4,288 blocks containing one gene start and 1,846 containing two to five. This representation prevented the same sequence or peak from being counted repeatedly at overlapping gene-centered promoters. The predictive model described below instead used one annotated 5′ boundary ±1-kb promoter per release-68 gene.

**Dinucleotide-shuffled discovery.** Positive sequences were full *q* < 0.075 peaks whose summits fell in single-gene promoter blocks of functional consensus-network targets, defined as an edge at any of 24 time points in at least 50% of the 61 Pareto-front networks. Discovery in this arm used the seven prespecified chromosomes Ib, III, V, VIIa, VIII, IX, and X; sequences came from the ToxoDB release-68 ME49 genome. Seventeen factors had at least 50 positive peaks on these discovery chromosomes and at least 30 on the remaining seven chromosomes. Each peak sequence was paired with one dinucleotide-preserving shuffle generated with fastashuffle-letters -kmer 2 -copies 1, using seed 20260928 plus the factor’s zero-based position in the alphabetical list of the 17 eligible factors. MEME 5.5.9 was run in discriminative ZOOPS mode on both strands (-objfun de -mod zoops -minw 6 -maxw 20 -evt 0.05 -searchsize 0 - maxsize 20000000); it stopped at the first motif with E > 0.05 and returned 1–7 retained motifs per factor [157]. STREME 5.5.9 returned five motifs at widths 6–15 with a 10% holdout and an order-2 background estimated from the shuffled negatives; all five were retained without an E-value filter. HOMER 5.1 (findMotifs.pl, -len 6,8,10,12,15 -S 5 -noredun) requested five raw motifs at each width, or 25 per factor, and its nonRedundant.motifs result retained 7–12 per factor without a *P*-value filter [158]. AP2XII-9 and Myb3 had no MEME motif at E ≤ 0.05 and were excluded together with their other-tool motifs, leaving 265 PWMs from 15 factors (55 MEME, 75 STREME, and 135 HOMER). The remaining seven chromosomes were used only for the factor-inclusion count and did not filter these PWMs.

**Matched-background discovery.** This arm used all 14 chromosomes and the ToxoDB release-52 ME49 genome. Positives were 37,460 summit-centered 101-bp windows. Each promoter block of width *W* contributed floor(*W*/101) centered, adjacent 101-bp grid windows, with the unused remainder divided between its ends; this yielded 130,021 windows. One control pool comprises grid windows that overlap no peak of the tested factor and whose centers are at least 250 bp from every summit of that factor; a second comprises windows centered on summits of the other 20 profiled factors and more than 100 bp from every summit of the tested factor. Controls were sampled without replacement within accessibility decile, window-GC quintile, number of other bound factors capped at 16, and block-position quartile, using seed 20261005 plus factor index. Position is floor(4*x*), where *x* is fractional block position oriented 5′→3′ for single-gene blocks and from the genomic start for multi-gene blocks. Two controls per positive were used when at least 95% of the attempted two-control draws matched the complete stratum, and one otherwise; this decision was made separately by factor and control pool. When an exact match was unavailable, the fixed relaxation order dropped position, then GC content, then coarsened accessibility to quintiles, then dropped accessibility; 94.5% of 92,498 draws were exact, and the other-factor count was never relaxed. STREME 5.5.9 was run separately against each control pool with eight motifs, widths 6–15, seed 0, a 10% holdout, and an order-2 background estimated from the negatives. Seventeen of 240 reported motifs, 12 from factor-free windows and five from other-factor summits, had holdout E ≤ 0.05 and were retained.

**Reference PWMs and candidate set.** Four prespecified reference sequences were added before similarity analysis: GCTAGC (AP2XI-5) [17], TGCATGCA (the *T. gondii* ribosomal-promoter element subsequently associated with AP2XII-8) [6,57], GTGTGTGT (from the preliminary AP2XI-4 analysis), and TATAWAWR (a core-promoter comparator). Each sequence was converted to a PWM by distributing total probability 0.94 equally among the IUPAC-allowed bases at each position and 0.06 equally among the remaining bases. These reference PWMs entered the same Tomtom comparison and representative-selection procedure as the discovered PWMs; they were not used as negative controls or validation data. Together with the 265 shuffled-background and 17 matched-background PWMs, they produced 286 candidates. Only the final cluster representatives entered the promoter-wide predictive model.

Tomtom 5.5.9 compared all 286 PWMs with themselves on both strands using Pearson distance, at least five overlapping positions, no small-sample correction, and a reporting threshold of *q* ≤ 0.05. Self-matches were discarded, the smaller *P* and *q* from the two query–target directions was used, and unreported pairs were assigned *P* = *q* = 1. With *L* = −log₁₀(*P*), similarity is *s* = *L*/(*L* + 4) and distance is *d* = 1 − *s*. Average-linkage clustering cut at *d* = 0.5 produced 99 initial clusters.

**Representative selection and non-redundancy.** Every candidate PWM was scanned against the summit, factor-free, and other-factor windows of all 15 factors on all 14 chromosomes. Scoring used base-2 log odds against the ToxoDB release-52 genomic background after adding 0.01 to each PWM cell and renormalizing each column. A window was counted as a hit when its score reached the PWM-specific 99th percentile over 69,244 peak-free promoter-grid windows, with ties included. For each PWM, the maximum Haldane-corrected Woolf *z* across factors was calculated first for summit versus other-factor windows (specificity) and then for summit versus factor-free windows (usage). A cluster representative is the member with the largest specificity *z* when any member reaches 3.29, otherwise the member with the largest usage *z* when any member reaches 3.29, and otherwise the member with the greatest total information content; ties follow cluster order. Clusters whose current representatives matched at Tomtom *q* < 0.01 were merged iteratively from the smallest *q*, with the representative reselected after each merge. Twenty-seven merges produced 72 non-redundant motifs (**Fig. S11B**): 18 representatives selected by specificity, nine by usage, and 45 by information content; 65 came from the shuffled arm, six from the matched-background arm, and one, TATAWAWR, from the reference set. The representatives for the GCTAGC, TGCATGCA, and GTGTGTGT families were de novo PWMs rather than the reference sequences themselves. We selected the 72-representative set after a post hoc comparison with a smaller representative-set alternative; summit association calls agreed apart from motif labels.

**Promoter-wide motif scoring.** Each of the 72 PWMs was scanned on both strands of the complete strand-oriented annotated 5′ boundary ±1-kb promoter of all 8,178 genes. A pseudo-count of 0.01 was added to each base probability and each PWM position was renormalized. With background frequencies *q* = (0.2379, 0.2620, 0.2620, 0.2381) for A, C, G, and T, respectively, a full-width site beginning at position *s* received the score

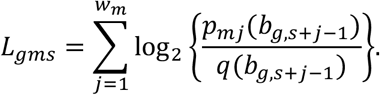

Sites containing N were skipped, and the continuous motif feature is *M*(*g,m*) = maxₛ *L*(*g,m,s*). No thresholded motif-presence call or target-specific motif subset entered the occupancy model. Scores were calculated once for all 8,178 genes; the primary model subsequently excluded 62 genes as described below, leaving 8,116. Within every model fit, each motif column was centered and scaled using only the fitting rows, and that transformation was applied unchanged to validation or test rows. The scanner agreed with an explicit brute-force implementation within 10⁻⁹ and with planted full-width optimal sites on both strands within 10⁻¹². The shared 72-motif vocabulary was defined before predictive modeling using information from all chromosomes. Leave-one-chromosome-out evaluation therefore tests whether these prespecified scores improve prediction on an unseen chromosome; it does not independently validate motif discovery.

**Discovery-stage factor–motif associations.** For each of 1,080 factor–motif pairs (15 factors × 72 motifs), hit rates in summit windows were compared with factor-free windows (usage) and other-factor summit windows (specificity) using Haldane-corrected ORs, Woolf 95% confidence intervals, two-sided Fisher exact tests, and BH correction over 1,080 tests per comparison [110]. A summit-specific and centered association requires a specificity OR ≥1.5, BH *q* ≤0.05, a 95% interval from 500 chromosome-bootstrap draws above one, a usage OR >1 with BH *q* ≤0.05, and a summit-to-flank centrality ratio ≥1.3. Sixteen associations met these criteria. As a replication check for these discovery-stage associations, STREME motifs were learned on one chromosome half, assigned to their best matching cluster at Tomtom *q* ≤0.05, and evaluated on the opposite half; 12 of the 16 summit-specific and centered associations reproduced. This replication check was separate from the primary leave-one-chromosome-out occupancy model.

#### Chromosome-held-out prediction of factor occupancy

**Analysis population and occupancy response.** The chromosome-held-out model shown in **Fig. 6B-D** and **Fig. S12** used one release-68 gene promoter as the unit of analysis, defined as the strand-oriented annotated 5′ boundary ±1 kb. The production motif scanner scored all 8,178 promoters. A promoter-similarity audit excluded 62 genes from the primary model so that sequence from a held-out chromosome could not be evaluated against a near-duplicate promoter already present in the fitting chromosomes. The frozen exclusion is the union of a 40-gene cross-half homology flag and 34 genes with same-half, cross-chromosome BLAST matches at ≥90% nucleotide identity across ≥100 bp; 12 genes belong to both sets. This conservative union contained all 55 genes with a direct cross-chromosome BLAST match at those thresholds. The two chromosome groups were used only to construct the exclusion and were not the validation folds of the final model. The resulting primary population contained 8,116 promoters; all 8,178 were retained in a robustness analysis.

The response *Y*(*f,g*) is one when factor *f* consensus-binds gene *g*. Binding was reconstructed from the same 61 Pareto-front network members by retaining each member’s CUT&RUN threshold, promoter assignment, strongest-peak-per-gene rule, and robust fold-enrichment gate while removing the knockdown differential-expression and tested-gene requirements. No occupancy-only Pareto front was reoptimized. A factor–gene pair is consensus-bound when it passes in at least 31 of the 61 members. Before the homology exclusion, this produced 48,603 binding pairs across all 21 profiled factors and 6,888 genes; the 15 modeled targets accounted for 47,845. The primary 8,116-promoter population retained 48,316 pairs across all 21 factors and 47,566 for the 15 modeled targets. Of the 48,603 reconstructed assignments, 12,207 lacked a knockdown row and were intentionally retained because occupancy, not functional response, was modeled.

**Modeled targets and partner pool.** The 15 targets were HDAC3, MORC, Cdc5, AP2X-7, AP2III-2, AP2VIII-7, AP2XI-5, AP2XII-8, DMAP1, EPL1, SNIP1, AP2IX-7, AP2XI-4, AAF1, and AP2IX-5. This named cohort was inherited from earlier **Fig. 6** development and fixed before the final leave-one-chromosome-out run; it does not estimate the prevalence of sequence dependence across all 21 profiled factors. For target *f*, *X*(*f*) contains the binary consensus-bound indicators of the other 20 profiled factors. All 20 indicators entered the complete model. The screenable partner pool comprises 18 factors that meet the inherited eligibility criterion of at least 30 consensus-bound promoters in each of two prespecified chromosome groups: the 15-target cohort plus Myb3, AP2XII-9, and ASF1. These groups were used only for eligibility, not as validation folds. Removing the modeled target left 17 individual partner tests per target. AP2XII-2, HDAC5, and AP2VIIa-6 remained nuisance covariates but were not screened individually.

**Promoter opportunity, factor environment, and sequence features.** Promoter opportunity comprises accessibility *A*(*g*) and five architecture variables *R*(*g*). Accessibility is the RH wild-type scATAC pseudobulk signal in the promoter, calculated from 15,091 Cell Ranger-called barcodes and 170,274,105 nuclear fragment rows. Each fragment row was counted once when it overlapped the 2-kb promoter by at least 1 bp, normalized to counts per million nuclear fragments and per kilobase, and transformed as ln(1 + CPM/kb). Architecture comprises GC fraction over non-N bases of the 2-kb window; an indicator that the window overlaps another gene’s annotated 5′ boundary ±1-kb window by at least 1 bp; indicators that the nearest upstream annotated 5′ boundary in the gene’s direction of transcription lies within 10 kb on the opposite strand (divergent) or the same strand (tandem); and log₁₀(distance in bp to the nearest other annotated 5′ boundary + 1). Accessibility, GC fraction, and nearest-boundary distance were centered and scaled on fitting rows only. Binary architecture variables remained 0/1. Zero-variance columns in a fitting set were omitted and recorded.

The factor environment comprises the number and identities of the other 20 bound factors. The raw partner count is *C*(*f,g*) = Σₕ *X*(*f,g,h*) and was standardized using the fitting rows. To represent which factors were present beyond how many, the 20-column indicator matrix was residualized separately in every fitting set. With *S* = [1,*C*],

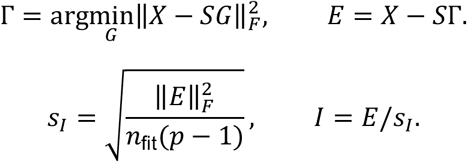

The fitting-set Γ and *s*_I_ were applied unchanged to validation and test rows. All *p* = 20 residual columns were retained and were not standardized individually. Each fit verified that row sums of *E* were below 10⁻⁹, ǁ*S*ᵀ*E*ǁ∞ <10⁻⁸, and rank[1,*C,E*] equaled rank[1,*X*]. Sequence is the 72 continuous promoter motif scores described above.

**Complete occupancy model and comparison models.** For each target factor, the complete occupancy model contains all five fitted components: accessibility (*A*), architecture (*R*), motif scores (*M*), partner count (*C*), and residual partner identities (*I*):

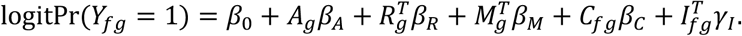

The training-prevalence reference contains only an intercept and assigns the same probability to every promoter on the omitted chromosome. Whole-layer contributions were evaluated by comparing the complete model with a reduced model that omitted the stated layer. The sequence-omission model removes all motif scores; the factor-environment-omission model removes both partner count and partner identities; and the identity-omission model removes residual partner identities while retaining count. Individual screens removed one motif score or one sufficiently represented partner from the complete model. The intercept, accessibility, architecture, and partner count were unpenalized. The sequence and residual-identity blocks had separate ridge penalties. With η(*g*) denoting the linear predictor, the minimized data-fit term is

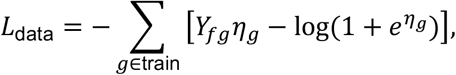

and the complete objective is

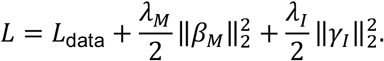

The custom Newton/iteratively reweighted least-squares solver used step halving, a convergence tolerance of 10⁻⁸, and at most 100 iterations. Non-convergence, extreme unpenalized coefficients, and fitted probabilities near zero or one were recorded for every fit.

**Nested leave-one-chromosome-out validation.** The outer loop left out one complete nuclear chromosome, fit on the other 13, and repeated this for all 14 chromosomes. Within each outer training set, all 13 remaining chromosomes served once as an inner validation fold, with the other 12 used for fitting. Every center, scale, zero-variance decision, and identity transformation was fitted only on the corresponding fitting rows. The motif and identity penalties were selected independently from 10⁻², 10⁻¹·⁵, …, 10⁵, plus literal omission of the entire block. Selection maximized pooled inner held-out natural-log likelihood across the 13 validation chromosomes. Exact ties favored greater shrinkage or omission, then the larger motif penalty. All 32 coalitions of *A*, *R*, *M*, *C*, and *I* were fitted, and each included penalized block was tuned independently; the complete criterion surface was retained.

**Held-out model value and feature-group Shapley decomposition.** For target *f* and feature coalition *S*, model value was pooled across the 14 outer folds before calculation:

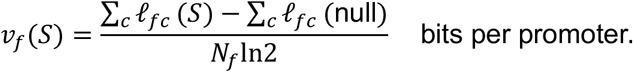

Here, *N*_f_ is the number of promoters pooled across the 14 held-out folds for factor *f* (8,116 in the primary analysis), held-out log likelihood ℓ(*f,c*) is the Bernoulli log likelihood summed over genes on test chromosome *c*, and the training-prevalence reference predicts the target’s prevalence in the corresponding training set. Held-out McFadden R² is 1 − Σ_c_ ℓ(*f,c;S*)/Σ_c_ ℓ(*f,c;null*).

#### Predictive performance on omitted chromosomes

Predictive performance was summarized from the saved per-promoter held-out linear predictors of the existing 14-fold leave-one-chromosome-out analysis; no model was refitted (**Fig. S13**). Predictions were pooled across the 14 omitted chromosomes so that each of the 8,116 promoters contributed once. We compared the complete **A** + **R** + **M** + **C** + **I** model; the raw partner count **C** and raw RH scATAC accessibility **A**, each used directly as a ranking without model fitting; and the 72 motif scores **M** fitted alone with the same fold-specific ridge-logistic procedure. Because **C** and **I** are CUT&RUN occupancy calls for the other profiled factors from the same series, these are conditional occupancy predictions and cannot be applied to a new factor or to a promoter lacking the corresponding occupancy measurements.

AUROC is the probability that a bound promoter ranks above an unbound promoter, with tied bound–unbound pairs contributing one half. Average precision, reported as AUPRC, used the exact expectation over random orderings within each tied-score group [159]; the random-ranking reference is the fraction of promoters bound. Precision among the top *n* uses *n* equal to the observed number of bound promoters and counts a tied group crossing the cutoff in proportion to the fraction selected; *n* was recomputed within each chromosome-bootstrap draw. This prevalence-matched cutoff is an evaluation summary rather than a fixed probability threshold. Calibration curves grouped the pooled held-out probabilities into deciles. Recalibration slope *b* and intercept *a* were estimated as descriptive diagnostics from logit Pr(*Y* = 1) = *a* + *bη*, where *η* is the pooled held-out log odds; *a* = 0 and *b* = 1 indicate perfect calibration, and no probabilities were recalibrated.

AUROC differences were calculated pairwise on the same promoters. Ninety-five percent intervals used the same 2,000 chromosome-bootstrap draws as **Fig. 6C-D** (seed 20261022): each draw sampled 14 chromosomes with replacement, both scores were evaluated with the same chromosome multiplicities, and the 2.5th and 97.5th percentiles were reported. These intervals are descriptive and conditional on the fitted models because the models were not refitted. The shared 72-motif set was selected using all-chromosome CUT&RUN data; leave-one-chromosome-out evaluation therefore holds out performance, not motif selection, and estimates for the complete model and **M** alone may be optimistic.

The production run fitted all 32 coalitions of the five model components. For the biological summary shown in **Fig. 6C**, accessibility and architecture were grouped as promoter opportunity *O* = (*A,R*), partner count and residual partner identity as factor environment *E* = (*C,I*), and motif scores as sequence *Q* = (*M*). Exact three-player Shapley values were calculated from the eight corresponding fitted coalitions:

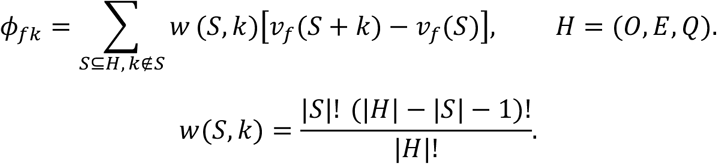

Here, *S* + *k* denotes coalition *S* after adding layer *k*. Shapley values allocate shared held-out predictive information and do not imply causal effects. Descriptive 95% intervals used 2,000 chromosome bootstraps with seed 20261022. Across-target percentages were calculated as the mean contribution divided by the mean complete-model value, not as the mean of target-specific percentages. The displayed mean shares were 39.34% for promoter opportunity, 55.06% for factor environment, and 5.60% for sequence.

**Feature-group omission tests.** Writing *v*_f_(*A,R,M,C,I*) for the complete model, the primary contrasts are

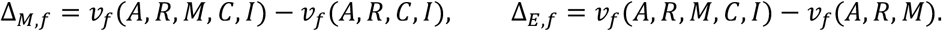

The identity-only contrast is

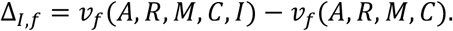

The sequence headline is the equal-weight mean Δ_M_ across the 15 targets. For any factor-level comparison of the complete model *F* with a model omitting feature or layer *j*, the contribution of omitted chromosome *c* is

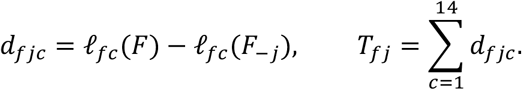

The one-sided screening value exhaustively enumerates the 2¹⁴ sign patterns:

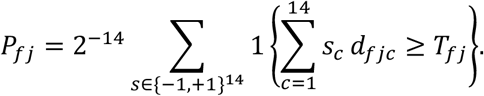

Thus, chromosomes—not genes, motifs, or factors—were the flipped units. For the aggregate sequence headline, contributions were first summed over the 15 targets within each chromosome and the same 14 chromosome signs were flipped jointly; the reported mean gain remained the equal-weight mean of the 15 target gains. These screening *P* values are approximate because leave-one-chromosome-out training sets overlap and sign symmetry is not guaranteed. Chromosome-bootstrap intervals were descriptive and never gated a call.

**Individual motif and partner omission screens.** Individual motifs were evaluated by removing one continuous PWM-score column from the complete model while locking the complete model’s fold-specific λ_M_ and λ_I_ (**Fig. 6D**, **Fig. 6I** and **Fig. S12B**). The motif model OR was calculated from the motif coefficient in the complete model:

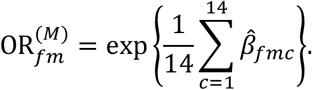

Here, motif *m* was standardized on each outer fold’s fitting promoters, and an omitted or zero-variance fold contributed β^ = 0. This OR is the conditional odds multiplier per one fitting-set SD increase in motif score; it is not the comparison between the complete and motif-omission models. For partner *h*, *h* was removed from both *X* and *C*, the residual identity block was rebuilt from the remaining 19 indicators on fitting rows, the complete model’s common identity scale was retained, and its penalties were locked. This is a removal and refit under the same modeling rule, not a literally nested penalized fit. The partner model OR instead toggles *h* from unbound to bound in the complete model while holding all other raw partner indicators and non-partner features fixed. Because *C* = Σ_h_ *X*_h_, this simultaneously increases partner count by one and updates the residual identity contrast:

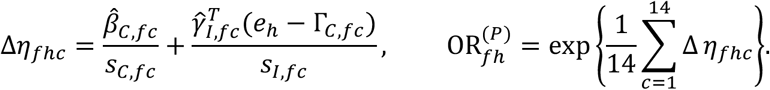

It is a conditional co-occupancy association, not the drop-one gain, an observed contingency-table OR, recruitment, or causality. Individual omission gains are not additive.

**Exploratory multiple-testing procedure and robustness checks.** BH adjustment was first applied separately across the 15 Δ_M_ tests and the 15 Δ_E_ tests. For the final exploratory figure and manuscript report, a layer passes when its gain is positive and BH *q* ≤0.10. This 0.10 reporting threshold was adopted after the production fit. The frozen production motif table records the prespecified *q* ≤0.05 procedure, under which nine detected sequence layers gave a within-target motif threshold of 0.05 × 9/15 = 0.03. Under the final post-production *q* ≤0.10 reporting rule, 10 sequence layers and all 15 factor-environment layers passed, giving within-target thresholds of 0.10 × 10/15 = 0.0667 for the 72 motif tests and 0.10 × 15/15 = 0.10 for the 17 partner tests. Only reporting labels were recalculated from the frozen screening *P* values; effect estimates and model fits were unchanged.

Within each layer that passed this first step, BH adjustment was applied separately within each target over its 72 motif tests or 17 partner tests. A motif or partner improves held-out prediction after multiple-testing correction when its gain is positive and its within-target *q* meets the corresponding threshold above. Predictive motifs with motif model OR ≥1.25 or ≤0.80 were marked separately; this model-OR criterion did not change the multiple-testing call. This two-stage procedure does not control one global false-discovery rate across all targets, motifs, and partners. For every element, the output includes held-out gain in bits per promoter, change in McFadden R², 14 chromosome contributions, coefficient direction, model OR, screening *P*, within-target *q*, and the model-OR decision. Partner findings do not establish simultaneous occupancy, physical interaction, recruitment, or direction of regulation.

The prespecified AP2XI-5 recovery check removed C01/GCTAGC with locked penalties and applied the same approximate sign-flip screen at *P* <0.05 outside the BH families. C01 is a learned PWM containing the published core rather than the reference sequence itself. For AP2XI-4, the prespecified related-feature block comprises C04, C21, C23, and C30, each matched to the GTGTGTGT reference PWM by Tomtom *q* <0.01. Two additional robustness checks were retained. First, 100 motif panels were formed by independently permuting the positions of every PWM, preserving width, columns, and information content; each was scanned identically and evaluated with the complete model’s fold-specific penalties locked. Percentiles were descriptive, not *P* values. Second, the four tuned model coalitions needed to calculate the three primary omission contrasts were repeated without excluding the 62 genes in the conservative promoter-homology set, using all 8,178 promoters. The production analysis was run with CPython 3.12.9, NumPy [160] 2.5.4, pandas [161] 3.0.6, and SciPy 1.18.1. Input, code, environment, and output SHA-256 values are recorded in production run fig5final_20261022_a99a853c, manifest digest 8501462bffee38312d1cd9e52a438f0a265c03a73eab0cd7c22a792c6d178836.

#### Palindromic motifs and lytic-cycle timing

**Analysis universe and palindromic 6-mer enrichment. Fig. 6E-H** used a pre-existing motif-timing analysis that was independent of the 8,116-promoter occupancy model. Promoter coordinates followed the release-68 annotated 5′ boundaries, while sequence extraction retained the ToxoDB release-52 ME49 genome used by the original analysis. The 64 DNA 6-mers that equal their reverse complement were scored as exact matches in the strand-oriented core window from −385 to +228 bp of the annotated 5′ gene boundary in each of 4,288 single-gene promoter blocks. One-strand scanning therefore captured both orientations; overlapping occurrences were counted. Equal-length gene-body controls from +1,000 to +1,613 bp were available for 3,892 genes with release-68 annotated spans of at least 1,700 bp. The 4,288 promoter and 3,892 control windows were compared as unpaired sets. For each 6-mer, the Mantel–Haenszel (MH) OR compared motif presence between the two sets within GC quintiles computed over the pooled windows. Robins–Breslow–Greenland standard errors gave 95% confidence intervals, Cochran–Mantel–Haenszel *P* values used the uncorrected one-degree-of-freedom statistic, and BH correction covered the 64 motifs. These analyses were exploratory.

**Wild-type timing and early-versus-late comparisons.** Scaled wild-type expression and promoter-accessibility profiles were used as supplied at 257 positions from 0 to 6 h; the repeated 6-h endpoint was removed, leaving 256 time points, and no additional scaling was applied. Peak is the earliest maximum. Rise is the time of maximum positive circular central difference after wrapped Gaussian smoothing with SD 0.25 h; profiles with range <10⁻⁹ have neither a peak nor a rise. Early is [0,3) h and late [4,6) h; [3,4) h was excluded. For each of 64 motifs and four measures—mRNA peak, mRNA rise, accessibility peak, and accessibility rise—motif occurrence was compared between early and late promoters by an MH OR within 50 accessibility-decile × core-window-GC-quintile strata. BH adjustment covered all 256 tests. **Fig. 6E** shows accessibility peak (1,353 early, 1,723 late, and 1,197 excluded promoters). These timing values are trajectories inferred from asynchronous cells rather than synchronized time courses.

**Promoter position and composition-preserving null.** For **Fig. 6F**, every offset in the strand-oriented −1,000 to +1,000-bp sequence of each of the 4,288 promoters was scanned. Each overlapping occurrence was placed at its first base, from −1,000 to +995 bp; 6-mers containing an unresolved base were excluded. Counts were summed over promoters, Gaussian-smoothed with SD 20 bp using nearest-end extension, and expressed as sites per 10 bp per 1,000 promoters. Positional share is the number of sites beginning from −150 through 0 bp divided by the number from −1,000 through +995 bp. For the null, each 2,001-bp promoter was divided into nineteen 100-bp bins and one 101-bp bin. Each bin was independently shuffled by a uniform Eulerian-path dinucleotide shuffle that preserved its first and last bases and all 16 dinucleotide counts, and the bins were rejoined before scanning so junction-spanning sites were retained. In each of 1,000 draws (seeds 30261008–30262007), every promoter was shuffled once. The one-sided empirical *P* value is {1 + number of null shares at least as large as observed}/1,001 and BH correction covered 64 motifs.

**Motif frequency across the cycle and binding status.** For **Fig. 6G**, the 4,273 promoters with an accessibility peak and the 3,515 with an mRNA peak were placed in 12 half-hour bins. Relative frequency in bin *j* is *r*_j_ = *q*_j_/*q*, where *q*_j_ is motif prevalence in that bin and *q* is prevalence over all promoters with a peak for that measure. Intervals were the 2.5th–97.5th percentiles from 1,000 chromosome-bootstrap samples (seed 20261019), recomputing *q*_j_ and *q* in each draw. Display lines were circular Gaussian weighted averages with SD 0.4 h and bin-count weights, evaluated from 0.015 to 5.985 h; this smoothing entered no test. Promoters with an early or late accessibility peak were divided into those with at least one *q* < 0.075 summit from any of the 21 profiled factors in the block (2,793 promoters; 1,248 early and 1,545 late) and those with none (283; 105 early and 178 late). Within each group, the early-versus-late MH OR used the same 50 strata. Heterogeneity was tested by comparing the two log ORs with their Robins–Breslow– Greenland standard errors; the two heterogeneity *P* values were not multiplicity-corrected.

**Timing of genes at factor-occupied promoters.** To describe where each factor’s occupied promoters fell in the lytic-cycle expression program, genes with a wild-type mRNA profile were assigned to the 1-h interval in which their mRNA peaked. A gene was counted as occupied when its single-gene promoter block contained a *q* < 0.075 CUT&RUN peak for that factor. For each factor and interval, we estimated the odds of occupancy among genes peaking in that interval relative to genes peaking in the other five intervals, within accessibility-decile × other-factor-count strata; BH correction covered the 90 factor-by-interval tests. A factor was classified as late-associated when at least one 4–6 h interval was enriched (OR ≥1.5; BH *q* <0.05) and no 0–3 h interval was enriched. Early-associated is defined conversely. No factor met both rules; the remaining factors met neither rule. The figures use this single primary classification. The labels describe the wild-type expression timing of genes at factor-occupied promoters, not when the factor binds or acts.

**Factor–motif associations.** For **Fig. 6H**, binding retained the original single-gene-block *q* < 0.075 peak calls used by the independent motif-timing analysis rather than the consensus-bound response of the leave-one-chromosome-out model. For each of the 15 targets and each of GCATGC and GCTAGC, binding was compared between promoters with and without the motif by an MH OR within accessibility-decile × other-factor-count strata, with the count capped at 16. BH correction covered 30 tests; enrichment requires OR ≥1.5 and depletion requires OR ≤1/1.5, with *q* <0.05.

**Predictive-partner network display.** Across the 17 eligible partner tests per target, 78 directional partner features passed the post-production *q* ≤0.10 reporting rule, had positive held-out gain, and had partner model OR ≥1.25 or ≤0.80. **Fig. 6I** is restricted to links among the 15 modeled factors: 71 directional calls collapsed to 39 unordered pairs, of which 32 passed in both modeled directions and seven in one direction; two pairs included a direction with partner model OR <1. A bidirectional call is drawn as an undirected line, whereas an arrow points from the informative partner toward the factor whose occupancy was predicted. Line width is the mean held-out ΔMcFadden R² ×100 across the represented directions, and a line is dashed when at least one represented direction has partner model OR <1. Node color shows the expression timing of genes at occupied promoters: orange, late-associated; blue, early-associated; gray, neither. Line color shows the class shared by both factors: orange for late–late pairs, blue for early–early pairs, and light green for all other pairs. The seven late–late pairs accounted for 44.9% of the summed displayed ΔMcFadden R², the seven early–early pairs for 13.6%, and the remaining 25 pairs for 41.5%. The neutral shaded backgrounds are descriptive partner groups, not tested modules. They were derived among the 15 modeled factors by summing directional held-out gains into a 15 ×15 matrix, symmetrizing it as *S* = *G* + *G*ᵀ, and applying average-linkage clustering. Node placement was optimized solely for legibility and has no quantitative meaning.

#### Structure prediction of complexes

Pairwise complexes between AAF1 (TGME49_260810; 277 residues) and individual *T. gondii* ApiAP2 proteins were predicted with AlphaFold 3 v3.0.1. Sequences were taken from the ToxoDB release 68 ME49 annotated proteome, and each input specified AAF1 as chain A and one candidate partner as chain B. Unpaired MMseqs2 multiple-sequence alignments were precomputed and supplied for both chains; neither paired alignments nor structural templates were used. Because alignments were embedded in the input files, the AlphaFold data pipeline was switched off at inference. Each prediction used model seed 1, five diffusion samples and ten recycles. Pairs with a combined length of 2,500 residues or fewer were run on NVIDIA A100 GPUs and longer pairs on an NVIDIA H200; this cut-off governed hardware allocation only and played no part in filtering proteins or classifying interactions. The model with the highest AlphaFold ranking score was kept for each pair, yielding a single observation per pair.

Interface confidence was described by three measures: the interface predicted template modelling (ipTM) score, the AlphaFold ranking score, and the mean minimum inter-chain predicted aligned error (PAE), the last computed as the average of the two off-diagonal entries of the chain_pair_pae_min matrix. Higher ipTM and ranking scores, and lower inter-chain PAE, correspond to greater confidence in the predicted interface. Fifty-six ApiAP2 proteins were analyzed. AAF1–AP2XI-5 and AAF1–AP2X-5 are shown individually, with the remaining 54 pairs combined into the AAF1–AP2 group. For comparison, 100 AAF1–non-AP2 pairs were generated by sampling the ToxoDB-68 ME49 proteome without replacement under a fixed random seed of 42, having first removed AAF1 itself, the ApiAP2 proteins, any protein annotated as carrying an AP2 domain, and any protein lacking a precomputed alignment. Pair-level values are plotted as individual points over box plots showing the median and interquartile range, with whiskers extending to 1.5 × IQR.

#### Interactive application

https://umbibio.math.umb.edu/toxohub/v4/

## Acknowledgements

Sharing reagents: antisera kindly shared by Drs. Peter J. Bradley (UCLA) and Iain M. Cheeseman (Whitehead Institute).

Funding through National Institutes of Health awards: AI152387 to MJG, AI174592 to KZ and MJG, AI167570 to MTD, KZ and MJG.

## Author contributions

Computational analyses: AA, NS, SA, PrPr, SD, OG, KZ

*P. falciparum* experiments: JiLi, GMD

*T. gondii* experiments: YW, JiLo, PO Proteomics experiments: PaPa, EW

Size exclusion chromatography: LSIT, MLA Wrote the draft: AA, SA, YW, MTD, KZ, MJG Funding acquisition: MTD, KZ, MJG

