## Supplementary Material for "A MultiOmics time-resolved systems map of gene regulation in *Toxoplasma gondii*"

#### Supplementary figures

Figure S1. Feature overview of all 21 selected factors in this study.
Figure S2. Genotype validation of all *T. gondii* transgenic parasite lines.
Figure S3. Plaque assays of all factor tagged and wild type parasite lines.
Figure S4. Expression pattern of all tagged proteins across the cell division cycle.
Figure S5. KD phenotypes for each mutant.
Figure S6. Single-cell RNA-seq processing.
Figure S7. Cell-cycle clock.
Figure S8. Cdc5 and SNIP1 splicing profiles.
Figure S9. Donut plot of spliceosome including all names.
Figure S10. Conditional PfCdc5 depletion phenotype analysis.
Figure S11. Construction and provenance of the shared motif set, related to Fig. 6
Figure S12. Held-out tests resolve factor-environment and motif contributions to occupancy prediction, related to Fig. 6
Figure S13. Conditional predictive performance of the occupancy model on omitted chromosomes, related to Fig. 6
Figure S14. AAF1 GRN GO-terms and AlphaFold-3 of AAF1 paired with all AP2 factors in *T. gondii* and *P. falciparum*
Figure S15. AAF1 pull down
Figure S16. Alphafold-3 modeled interaction domains in the AAF1-AP2 complexes.
Figure S17. All IFA panels corresponding with Fig. 7H-I.
Figure S18. Cell-cycle density and deviation from WT.
Figure S19. Differential expression within time windows.
Figure S20. scATAC processing; expression and accessibility profiles.
Figure S21. CUT&RUN processing, peak calling, peak–gene assignment and the edge rule.
Figure S22. Ensemble clustering and gene communities.
Figure S23. Parameter sweep and consensus networks.
Figure S24. Community identity (Functional Annotator).
Figure S25. Regulator similarity (method for Fig. 3E).

#### Legends to Supplementary Tables

#### Legends to Supplementary Movies

| Factor type | GeneID: TGME49_ | Name | Description | Fitness |
|---|---|---|---|---|
| | 253380 | AP2III-2 | AP2 Domain Transcription Factor AP2III-2 | -3.22 |
| | 203050 | AP2VIIa-6 | AP2 Domain Transcription Factor AP2VIIa-6 | +1.27 |
| | 269010 | AP2VIII-7 | AP2 Domain Transcription Factor AP2VIII-7 | -1.01 |
| | 290630 | AP2IX-7 | AP2 Domain Transcription Factor AP2IX-7 | -4.22 |
| | 289710 | AP2IX-5 | AP2 Domain Transcription Factor AP2IX-5 | -5.29 |
| AP2 | 214840 | AP2X-7 | AP2 Domain Transcription Factor AP2X-7 | -3.39 |
| | 315760 | AP2XI-4 | AP2 Domain Transcription Factor AP2XI-4 | +0.29 |
| | 216220 | AP2XI-5 | AP2 Domain Transcription Factor AP2XI-5 | -4.57 |
| | 217700 | AP2XII-2 | AP2 Domain Transcription Factor AP2XII-2 | -1.16 |
| | 250800 | AP2XII-8 | AP2 Domain Transcription Factor AP2XII-8 | -4.32 |
| | 251740 | AP2XII-9 | AP2 Domain Transcription Factor AP2XII-9 | -4.32 |
| | 321450 | Myb3 | Myb family DNA-binding domain protein 3 | -4.21 |
| Myb | 275480 | Cdc5 | Cell Division Cycle 5 like protein | -4.24 |
| | 213890 | DMAP1 | DNA Methyltransferase 1 Associated Protein 1 | -2.72 |
| | 227290 | HDAC3 | Histone Deacetylase 3 | -5.36 |
| EpiF | 202230 | HDAC5 | Histone Deacetylase 5 | -3.66 |
| | 305340 | MORC | Microrchidia Protein | -4.69 |
| | 209140 | ASF1 | Anti-Silencing Protein 1 | -4.03 |
| | 225000 | EPL1 | Enhancer of Polycomb-like Protein 1 | -5.36 |
| other | 201790 | SNIP1 | Smad Nuclear interacting Protein 1 | -3.70 |
| | 260810 | AAF1 | AP2 associated Protein 1 | -3.97 |

**Figure S1. Feature overview of all 21 selected factors in this study.** DMAP1 was originally selected as a Myb factor, but is an epigenetic factor; EPL1 was originally selected for its poly-comb-like DNA binding domain, but is an epigenetic factor.

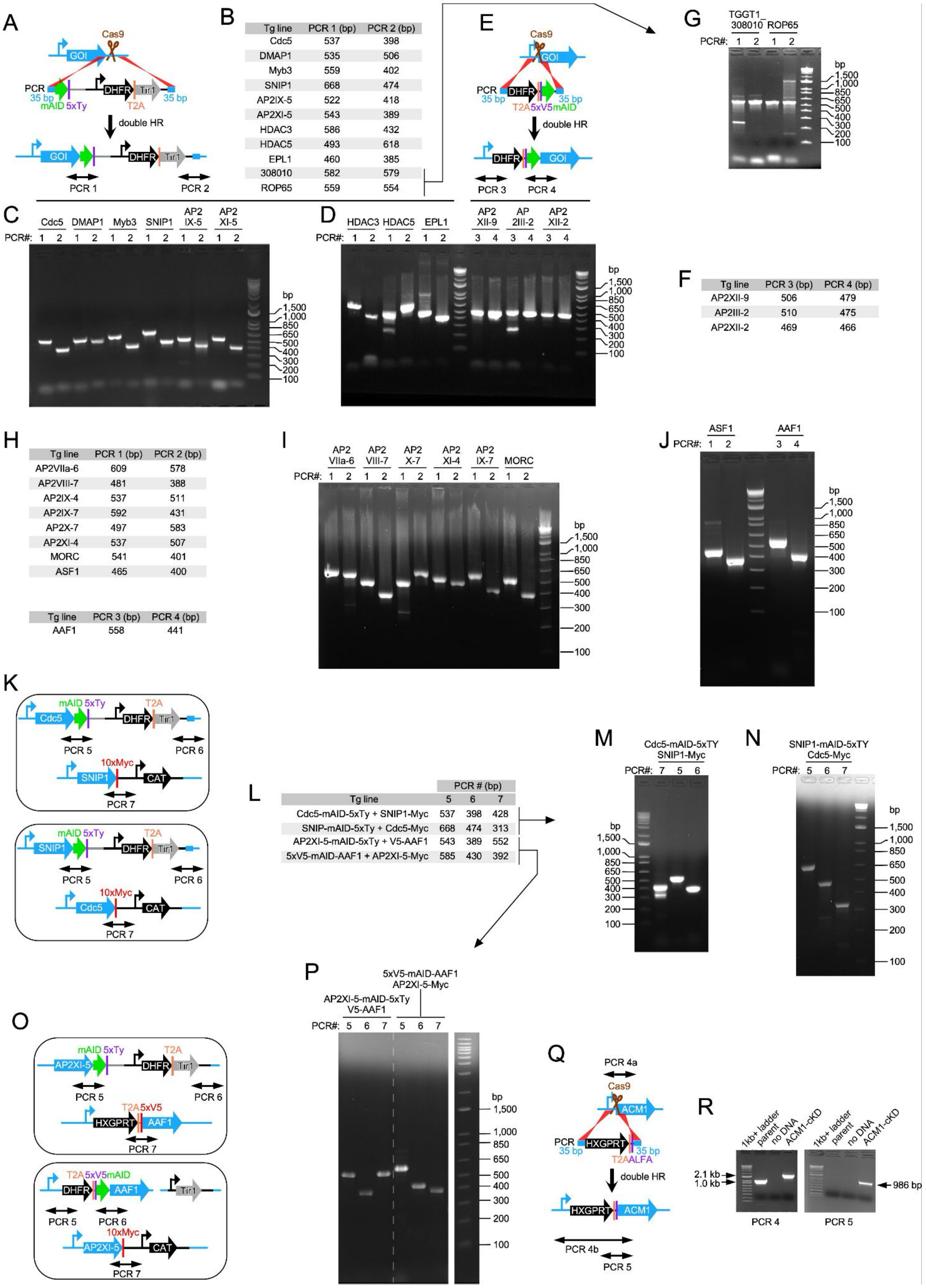


**Figure S2. Genotype validation of all *T. gondii* transgenic parasite lines.** (A, E, K, O, and Q) Schematics of transgenic *T. gondii* strains generated.
(B, F, H, and L) Predicted PCR band sizes; PCR numbers correspond with schematics.

(C, D, G, I, J, M, N, P, and R) Agarose gels of the Genotype validation by diagnostic PCRs. PCR numbers correspond with schematics.

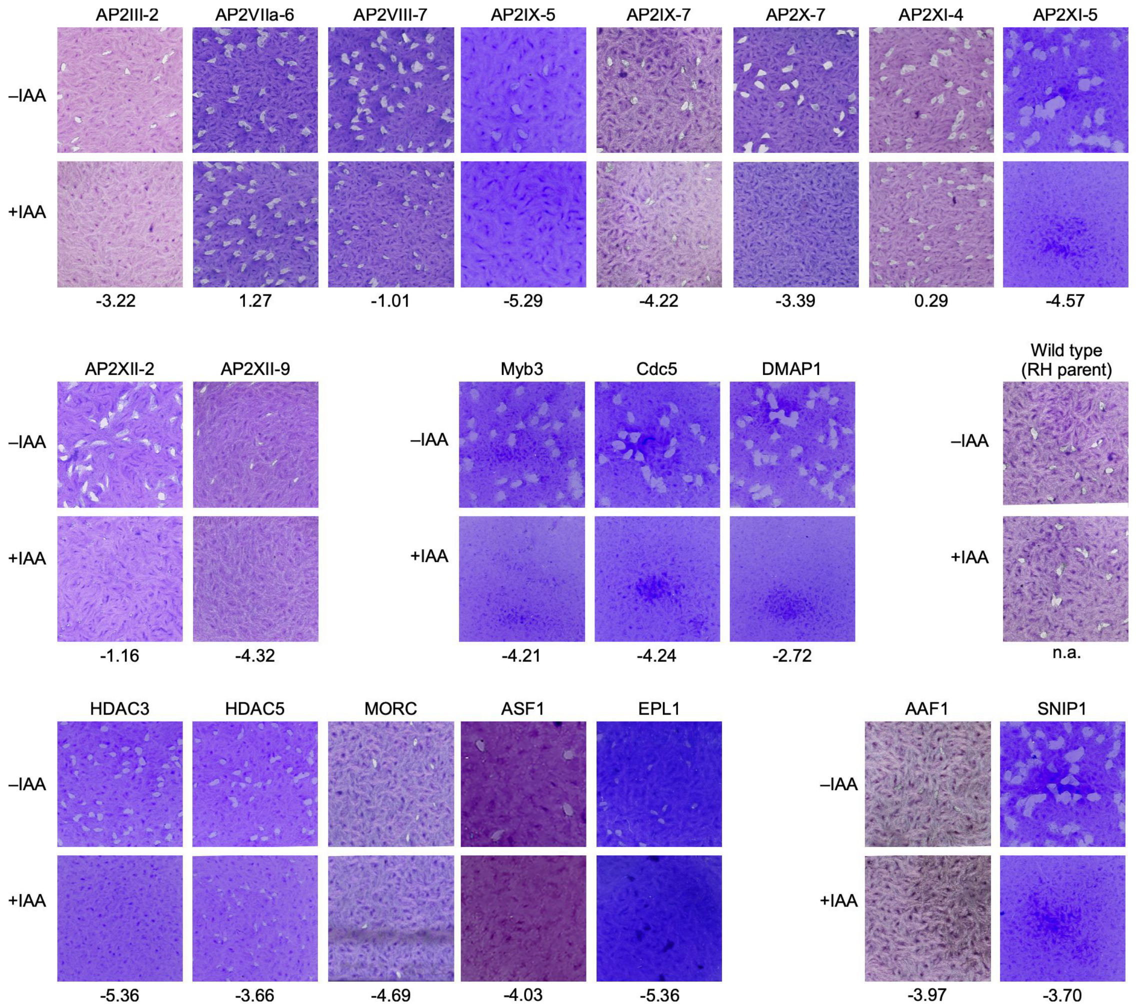


**Figure S3. Plaque assays of all factor tagged and wild type parasite lines.** Plaque assays were performed for 8 days ± IAA as indicated. The number below the panel refers to the genome wide tachyzoite fitness score [31].

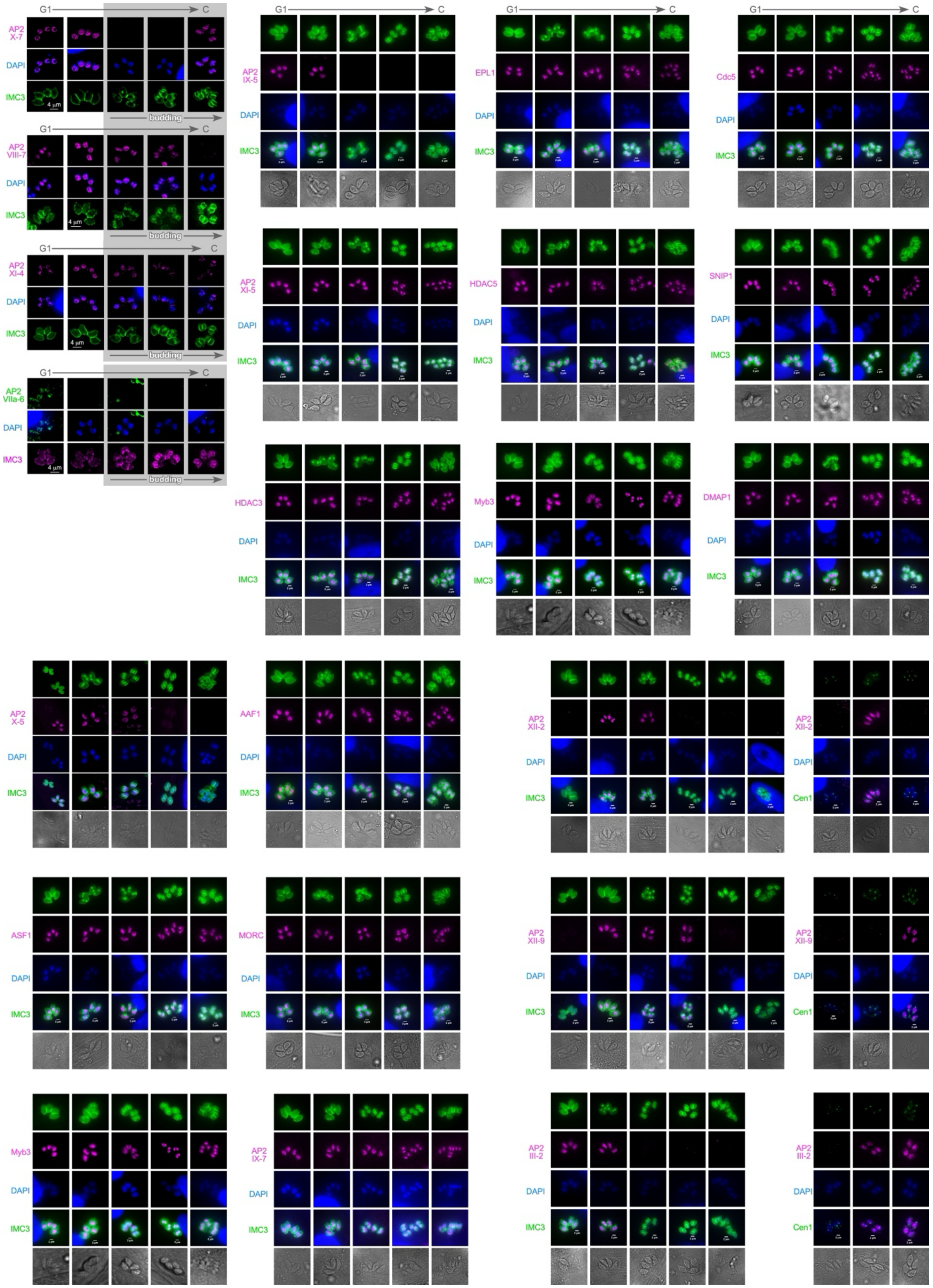


**Figure S4. Expression pattern of all tagged proteins across the cell division cycle.** Timing of factor presence by Ty epitope tag measured against daughter scaffold (IMC3) and DAPI cycle. Time progresses from left to right, starting in G1 and ending with completion of cell division. For factors peaking around S- and M-phase, more precise timing was assessed by co-staining the Ty-tagged protein with α-Centrin1 for the centrosome cycle.

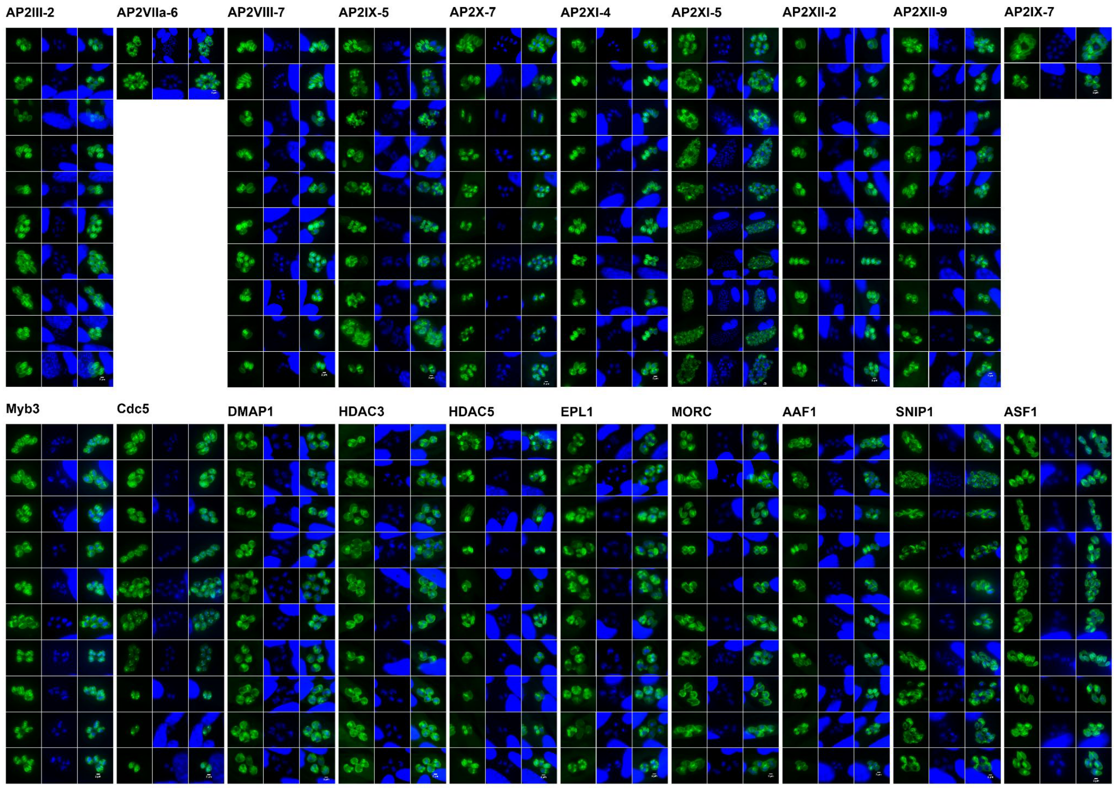


**Figure S5. KD phenotypes for each mutant.** Ten panels are shown, unless less than 10 out of 100 vacuoles observed showed a deviating phenotype.

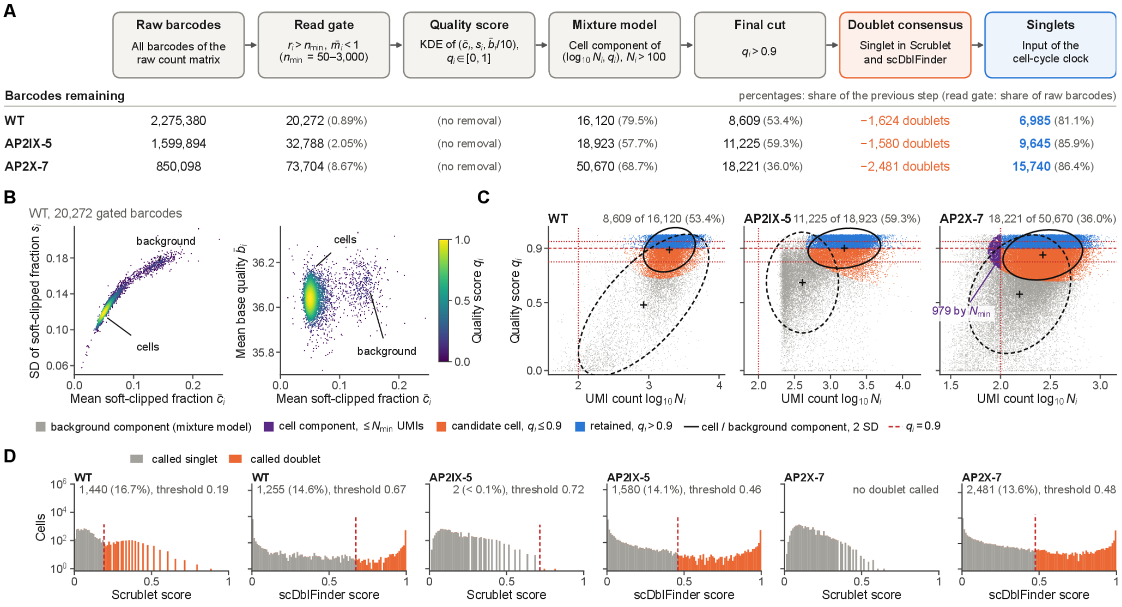


**Figure S6. Single-cell RNA-seq processing.**
**(A)** Barcodes remaining after each processing step, shown for WT, AP2IX-5 and AP2X-7. Steps, left to right: raw barcodes; the read gate; the quality score q (no barcodes removed); the mixture model; the final cut q > 0.9; doublet consensus; singlets. Percentages give the share of the previous step; for the read gate, the share of raw barcodes. Example: WT goes from 2,275,380 raw barcodes to 6,985 singlets.
**(B)** What the quality score q measures (WT, 20,272 gated barcodes). Left: SD against mean of the soft-clipped fraction. Right: mean base quality against mean soft-clipped fraction. Points are colored by q; cells and background separate.
**(C)** WT candidate cells by score band in the mixture model (WT, AP2IX-5, AP2X-7). Axes are UMI count (log10 N) and q; ellipses mark the cell and background components at 2 SD. Blue barcodes, q > 0.9, are retained; orange barcodes, 0.8 < q ≤ 0.9, are excluded.
**(D)** Doublet removal: Scrublet and scDblFinder score distributions with their thresholds, for the same three libraries. A cell is removed when either tool calls it a doublet.

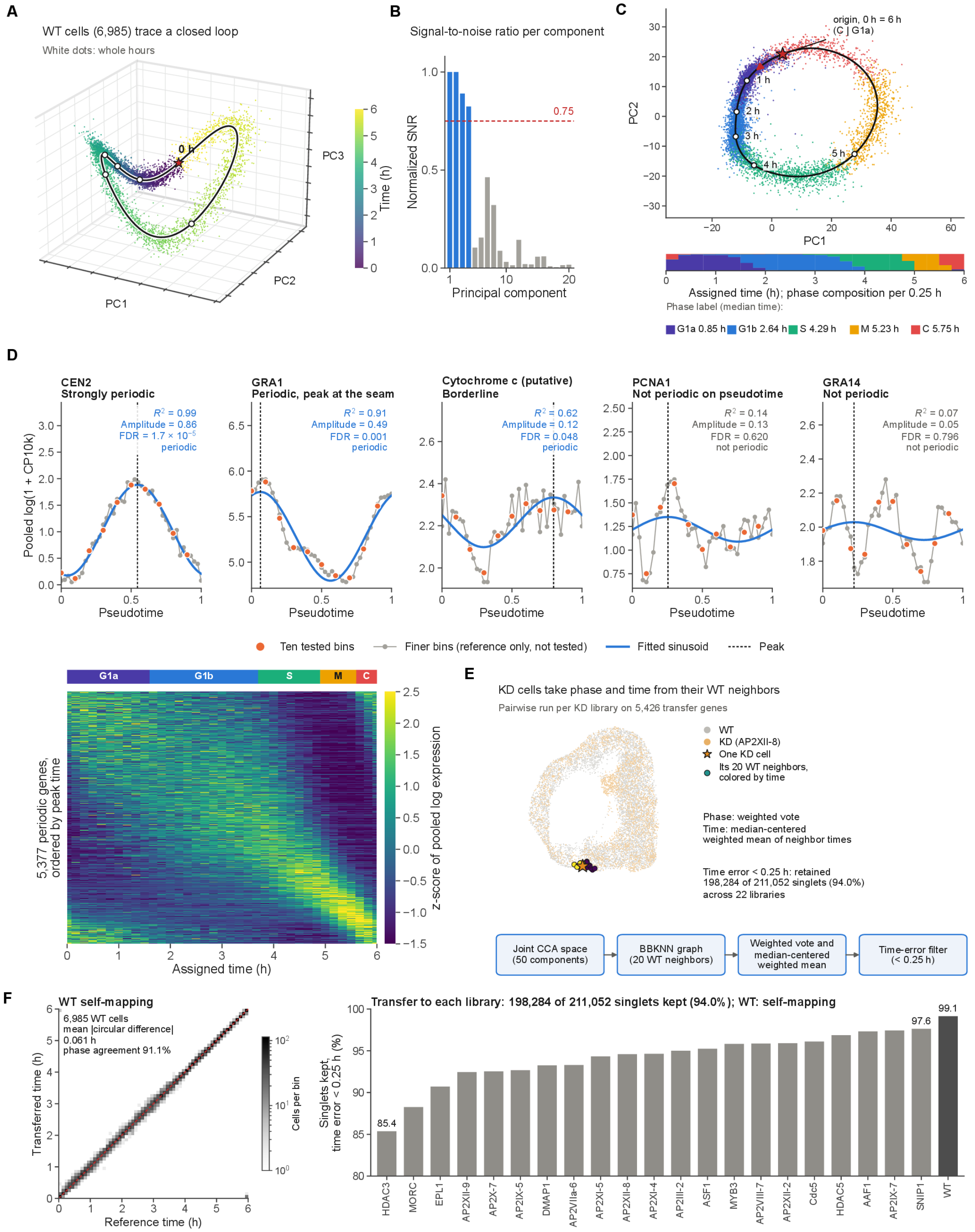


**Figure S7. Cell-cycle clock.**

**(A)** Reference clock. 6,985 WT cells trace a closed loop in the space of the leading principal components (PC1–PC3 shown). The black curve is the clock, white dots mark whole hours, and the star marks 0 h. Color shows assigned time.

**(B)** Normalized signal-to-noise ratio of each principal component along the initial order. PC1–PC4 pass the 0.75 line and are kept.

**(C)** Phase labels and the chosen origin. WT cells on PC1–PC2 are colored by phase label (G1a,

G1b, S, M, C), with the clock curve and hour marks. The origin, 0 h = 6 h, lies between C and G1a (red star); the clock's direction was checked by hand. The strip below gives the phase composition per 0.25 h of assigned time; the key gives each phase's median time. Phase labels come from the Smart-seq2 reference (GSE145080).

**(D)** Transfer features. Top: cosinor fits for five example genes, from strongly periodic (CEN2) to not periodic (GRA14). Ten tested bins, finer bins for reference only, the fitted sinusoid and its peak are shown. Bottom: z-scored pooled expression of the 5,377 periodic genes (FDR $< 0.05$), ordered by peak time. The phase strip runs above the heatmap.

**(E)** Transfer of time and phase to the knockdowns. Each KD library is run pairwise against WT on 5,426 transfer genes. The steps are a joint CCA space (50 components), a BBKNN graph (20 WT neighbors), and then a weighted vote for phase and a median-centered weighted mean of neighbor times. Cells with time error $\geq 0.25$ h are removed. AP2XII-8 is shown.

**(F)** Validation of the transfer. Left: WT self-mapping. Reference time against transferred time for the 6,985 WT cells; mean circular difference 0.061 h, phase agreement 91.1%. Right: singlets kept per library (time error $< 0.25$ h), from 85.4% for HDAC3 to 97.6% for SNIP1. Overall, 198,284 of 211,052 singlets (94.0%) are kept.

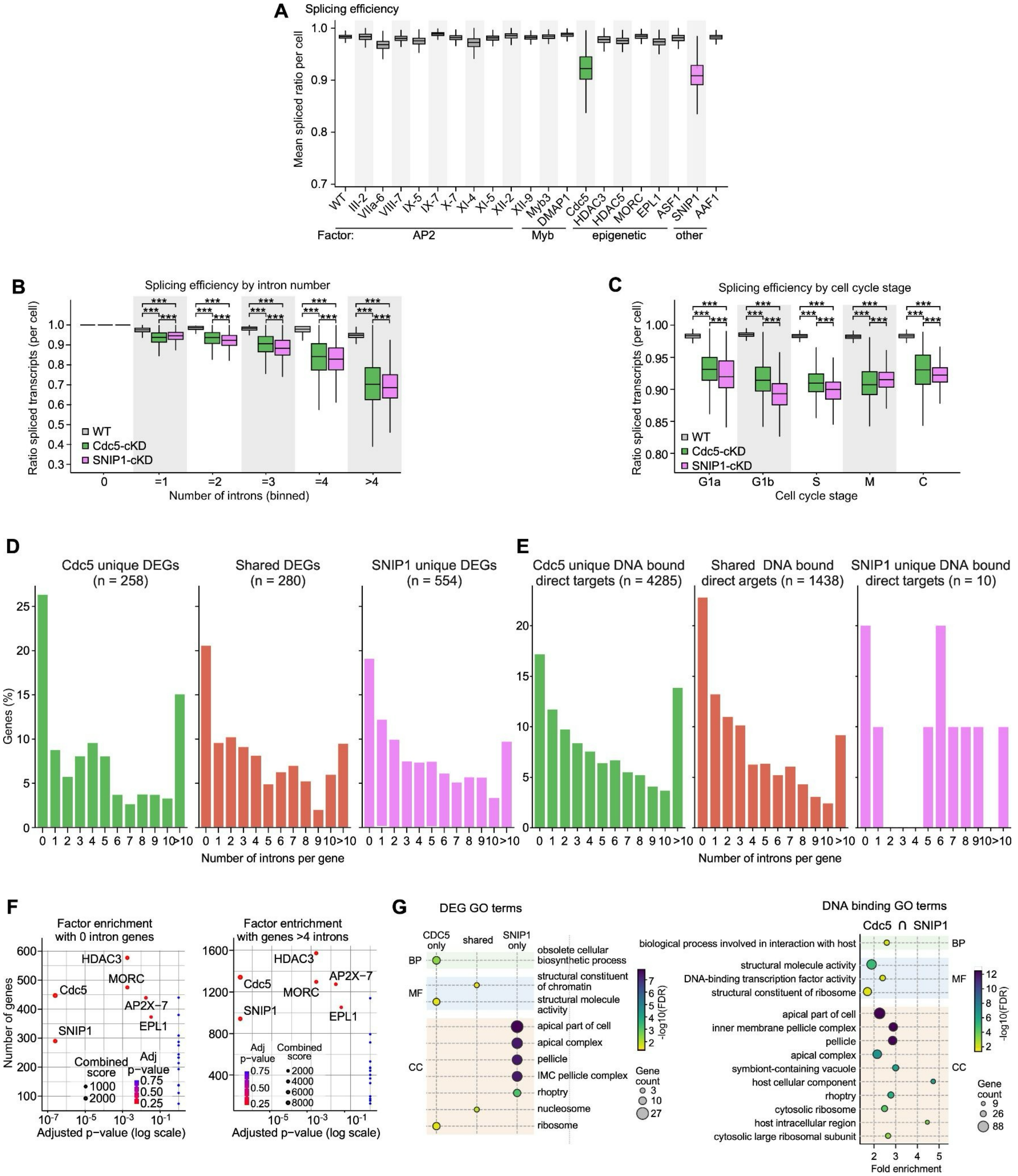

**Figure S8. Cdc5 and SNIP1 splicing profiles.**

**(A)** *T. gondii* splicing efficiency for all factors in this study shows that only depletion of Cdc5 and SNIP1 results in significantly reduced splicing.

**(B)** Splicing efficiency of wild type (WT) compared to Cdc5 and SNIP1 depleted cells resolved by number of introns reveals a trend of less complete splicing in genes with increasing intron number.

**(C)** Splicing efficiency of wild type (WT) compared to Cdc5 and SNIP1 depleted cells resolved by cell cycle stage does not reveal any differences in efficiency throughout the cell division cycle.

**(D)** Intron distribution among Cdc5- and SNIP1-depletion-responsive genes suggests roles beyond splicing, potentially involving transcriptional regulation, including intron-deprived genes.

**(E)** Intron distribution among Cdc5- and SNIP1 DNA engaged genes (CUT&RUN data) suggests roles beyond splicing, potentially involving transcriptional regulation, including intron-deprived

genes.
**(F)** Correlation of intron number with factor binding in the promoter (CUT&RUN data). Red marks: significant factors (labeled); blue marks: non-significant factors. Cdc5 and SNIP1 DNA binding are the strongest factors where binding correlates with increasing intron number.
**(G)** GO terms of genes directly engaged by Cdc5 and SNIP1.

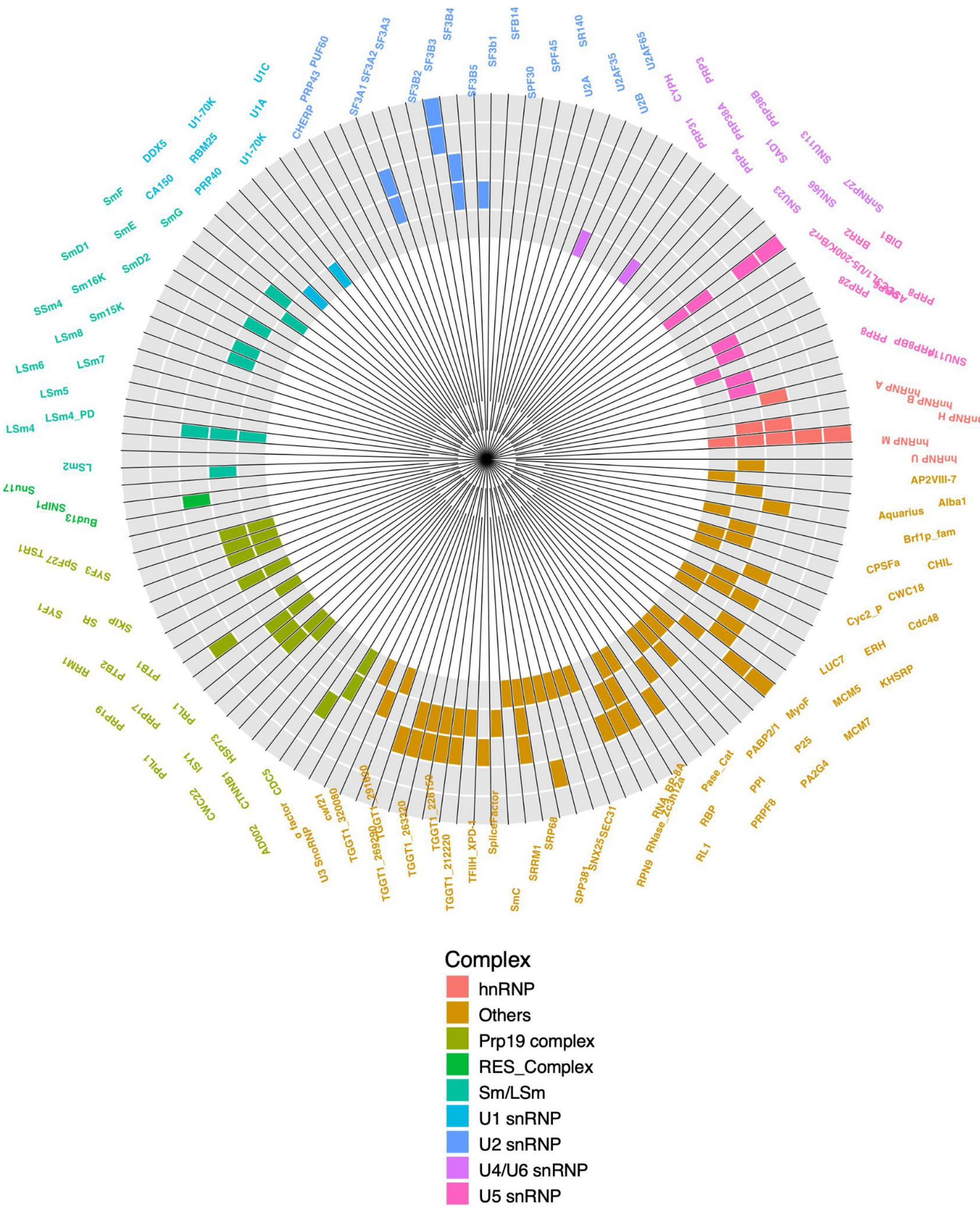


**Figure S9. Donut plot of spliceosome including all names.**

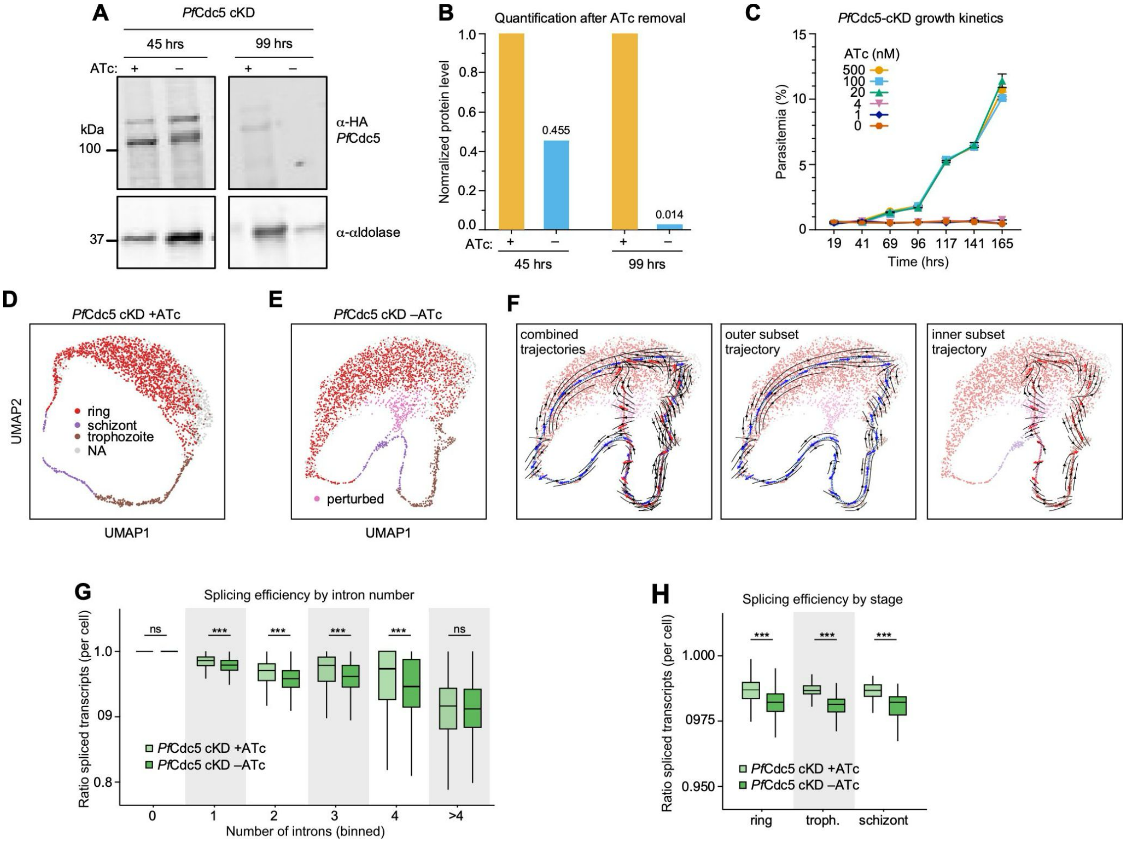


**Figure S10. Conditional PfCdc5 depletion phenotype analysis.**

**(A)** Western blot confirming depletion of HA-tagged PfCdc5 in NF54 PfCdc5-TetR-DOZI parasites at 45 h and 99 h after ATc (500 nM) removal, compared with +ATc controls. Aldolase is the loading control; molecular weight markers are in kDa.

**(B)** Densitometric quantification of the PfCdc5 western blot in panel A. PfCdc5 band intensities were normalized to the aldolase loading control and are expressed relative to the +ATc condition at each time point (45 h and 99 h).

**(C)** Titration of *Pf*Cdc5-TetR-DOZI parasites at different ATc concentrations (500, 100, 20, 4, 1 and 0 nM). Parasitemia (%) was monitored from 19 to 165 h. Parasites grown in ≥20 nM ATc expanded exponentially, reaching about 10-11% parasitemia by 165 h, with indistinguishable growth at 20, 100 and 500 nM. Growth was completely blocked at ≤4 nM ATc, with parasitemia staying at baseline, as in the no-ATc (0 nM) condition. These data identify 20 nM as the minimal ATc concentration that fully sustains growth. Concentrations at or below 4 nM give a fully penetrant growth defect.

**(D and E)** scRNA-seq UMAP projections of PfCdc5-TetR-DOZI parasites under permissive (+ATc, D) and restrictive (−ATc, E) conditions. For the restrictive condition, ATc (20 nM) was removed 20 h before harvest. Cells are colored by IDC stage (ring, trophozoite, schizont; NA, unassigned). Cells in (E) that do not map onto a normal stage are labeled “perturbed”.

**(F)** RNA-velocity analysis of PfCdc5-depleted cells (−ATc), shown for the combined, outer-subset and inner-subset trajectories. Arrow directions come from guided, pseudotime-based scVelo velocity vectors, and black streamlines are interpolated from these vectors. Colored arrows show locally averaged velocity vectors. Their placement along the guide paths and their colors are visualization choices, but their directions are velocity-derived.

**(G and H)** Splicing efficiency in PfCdc5-cKD parasites with (+ATc, light green) and without ATc (−ATc, dark green), measured as the ratio of spliced transcripts per cell. Transcripts are grouped by intron number (G; 0, 1, 2, 3, 4 and >4 introns) and by IDC stage (H; ring, trophozoite and schizont). Depletion of PfCdc5 significantly reduced splicing efficiency for transcripts with 1-4 introns but no significant difference was seen for intron-deprived transcripts(G). Depletion of PfCdc5 significantly reduced splicing efficiency for transcripts in all three stages (H) but no significant difference in gene with >4 introns (G). Box plots show the median (center line), interquartile range (IQR; 25th-75th percentiles; box) and whiskers extending to the most extreme data points within 1.5 × IQR of the lower and upper quartiles. ***$p < 0.001$; ns, not significant.

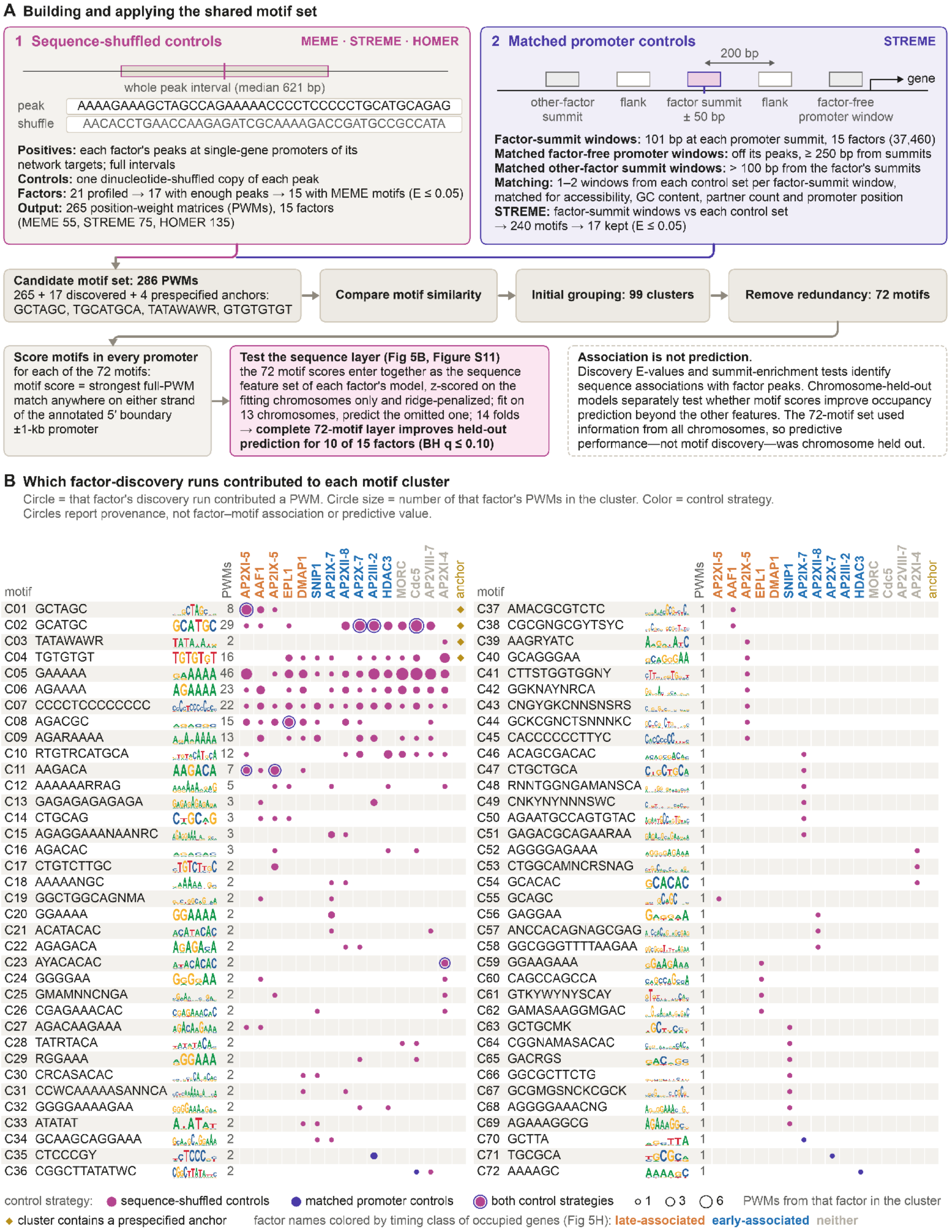


**Figure S11. Construction and provenance of the shared motif set, related to Fig. 6.**
**(A)** The shared set was built through two complementary discovery arms. In the sequence-shuffled arm, full CUT&RUN peak intervals at single-gene promoters of functional network targets were compared with one dinucleotide-preserving shuffle of each peak. The positives comprised 8,694 peaks from seven prespecified discovery chromosomes (median interval, 621 bp). Of 21 profiled factors, 17 met the peak-count requirement and 15 had at least one MEME motif at *E*

≤0.05; the retained 265 candidate PWMs comprised 55 MEME, 75 STREME, and 135 HOMER motifs from those 15 factors. In the matched-promoter arm, 37,460 factor-summit windows of 101 bp from the same 15 factors were compared separately with factor-free promoter windows located off that factor's peaks and at least 250 bp from its summits, and with other-factor summit windows more than 100 bp from its summits. One or two windows from each control pool were matched to each factor-summit window for accessibility, GC content, partner count, and promoter position. STREME reported 240 motifs, of which 17 met $E \leq 0.05$. The flanking windows shown 200 bp from the factor summit supported the separate summit-centering assessment and were not matched discovery controls. The 265 sequence-shuffled candidates, 17 matched-promoter candidates, and four prespecified anchors—GCTAGC, TGCATGCA, TATAWAWR, and GTGTGTGT—formed a 286-PWM candidate set. Similarity comparison produced 99 initial clusters; 27 subsequent representative merges yielded 72 non-redundant motifs. Each of the 72 PWMs was scanned on both strands of all 8,178 annotated 5′ boundary ±1-kb promoters, retaining the strongest full-PWM score per promoter. The primary occupancy analysis used 8,116 promoters after its promoter-similarity exclusion. Within every model fit, motif scores were standardized using only the fitting chromosomes. Models fitted on 13 chromosomes and predicted the omitted chromosome over 14 folds; the complete 72-motif layer improved held-out prediction for 10 of 15 factors under the exploratory BH $q \leq 0.10$ layer test. Discovery $E$ values and summit-enrichment tests identify motif associations with factor peaks, whereas chromosome-held-out models test predictive contribution beyond the other promoter features. Because the shared motif set used information from all chromosomes overall, predictive performance—not motif discovery—was chromosome held out.

**(B)** Provenance of the 72 non-redundant motif clusters. Each row gives the cluster identifier, representative consensus and logo, and total number of candidate PWMs assigned to that cluster. A circle indicates that the named factor's discovery run contributed at least one candidate PWM; circle size represents the number of that factor's candidate PWMs assigned to the cluster. Magenta and violet circles denote the sequence-shuffled and matched-promoter arms, respectively, and outlined circles indicate contributions from both control strategies. Gold diamonds mark clusters containing a prespecified anchor. Factor-name colors give the timing class of occupied genes, using the **Fig. 6H** definitions: orange, late-associated; blue, early-associated; gray, neither. Circles report discovery provenance only; they do not denote factor–motif association, predictive contribution, or binding of that factor to the displayed motif.

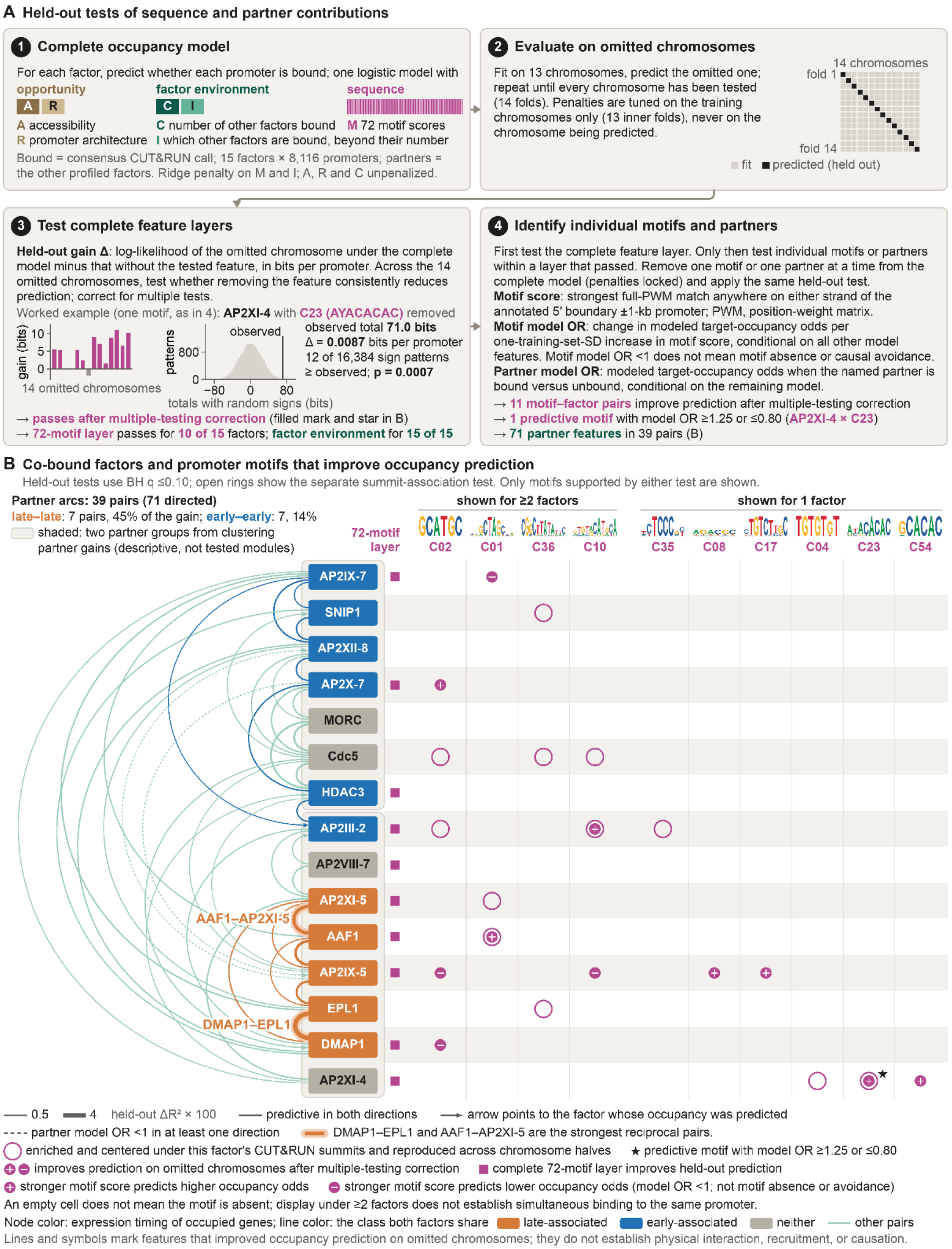


**Figure S12. Held-out tests resolve factor-environment and motif contributions to occupancy prediction, related to Fig. 6.**

**(A)** For each of 15 target factors, a nested leave-one-chromosome-out logistic model predicted consensus CUT&RUN occupancy across 8,116 gene promoters. Each outer fold fitted 13 chromosomes and predicted the omitted chromosome; penalties were tuned only within the 13 training chromosomes. The complete model contained accessibility (**A**), promoter architecture (**R**), the number of other bound factors (**C**), their identities beyond that number (**I**), and 72 promoter-wide motif scores (**M**). **M** and **I** were ridge-penalized; **A**, **R**, and **C** were unpenalized. A motif score was the strongest full position-weight-matrix match on either strand of the annotated 5′ boundary ±1-kb promoter. Held-out gain Δ was the complete-model minus reduced-model log-likelihood on omitted promoters, pooled across chromosomes and expressed in bits per promoter; Δ >0 indicates improved prediction. Approximate one-sided sign-flip tests enumerated all $2^{14}$ sign patterns of the 14 chromosome-level gains.

Testing proceeded in two stages. First, Benjamini–Hochberg (BH) correction was applied separately across the 15 target-factor tests for the complete 72-motif layer and the complete factor-environment layer; a layer passed with positive gain and BH $q$ ≤0.10. Individual features were then tested only within a layer that passed, by removing one motif or one eligible partner at a time while locking the complete model's penalties. Within each eligible target, BH correction covered 72 motifs or 17 partners, and the second-stage threshold was 0.10 × $k$/15, where $k$ was the number of targets whose corresponding layer passed. Thus, the threshold was $q$ ≤0.0667 for motifs ($k$ = 10) and $q$ ≤0.10 for partners ($k$ = 15). These exploratory $q$ ≤0.10 reporting rules were chosen after the production fit. The frozen execution plan and run metadata retain the prespecified $q$ ≤0.05 rule; the displayed $q$ ≤0.10 calls were recalculated from the saved screening $P$ values without refitting any model or recomputing any effect estimate.

The motif model odds ratio (OR) is the change in modeled target-occupancy odds per one-training-set-standard-deviation increase in motif score, conditional on all other model features. The partner model OR is the modeled target-occupancy odds when the named partner is bound versus unbound, conditional on the remaining model. Model OR ≥1.25 or ≤0.80 was reported separately as a larger-effect marker and did not alter the multiple-testing call. In the worked AP2XI-4–C23/AYACACAC example, 11 of 14 chromosome gains were positive, the observed total gain was 71.0 bits, Δ was 0.0087 bits per promoter, and 12 of 16,384 sign patterns were at least as large ($P$ = 0.0007; within-factor $q$ = 0.026; motif model OR = 1.365). Overall, the 72-motif layer passed for 10 of 15 factors and the factor-environment layer for all 15. Eleven factor–motif pairs improved held-out prediction after multiple-testing correction, and AP2XI-4–C23 was the only one that also met the model-OR criterion. The partner analysis returned 78 directed calls; the 71 connecting the 15 modeled factors are shown in (B).

**(B)** Unified partner and motif map. Within each target factor, the partner-testing family comprised the other 14 modeled factors plus MYB3, AP2XII-9, and ASF1. Seven passing calls involving MYB3, AP2XII-9, or ASF1 remain in the multiple-testing families but are not drawn; the 71 displayed directional calls connect the 15 modeled factors. They form 39 pairs: 32 reciprocal pairs are drawn without arrowheads, and seven one-way calls point toward the factor whose occupancy was predicted. All displayed partner calls had positive held-out gain, passed the second-stage BH rule, and had a partner model OR ≥1.25 or ≤0.80. Arc width is the mean held-out ΔMcFadden $R^2$ ×100 across the displayed direction or directions; dashed arcs have a partner model OR <1 in at least one direction. DMAP1–EPL1 and AAF1–AP2XI-5 are the two strongest reciprocal pairs.

Node color denotes the expression timing of occupied genes: orange, late-associated; blue, early-associated; and gray, neither. Arc color denotes the timing class shared by both factors: orange for the seven late–late pairs, which account for 44.9% of the summed held-out gain; blue for the seven early–early pairs, which account for 13.6%; and light green for the other 25 pairs, which account for 41.5%. The two neutrally shaded groups were obtained by average-linkage clustering of symmetric partner gains. They are descriptive groupings, not tested modules, and the shading has no timing meaning.

In the motif matrix, an open magenta ring marks a summit association: the motif was enriched and centered beneath that factor's CUT&RUN summits and reproduced across chromosome halves. A filled magenta disc marks a motif that improved held-out prediction after multiple-testing correction; the white plus or minus reports whether a stronger motif score predicted higher or lower modeled occupancy odds, respectively. A minus does not mean motif absence or causal avoidance. A star marks a predictive motif with model OR ≥1.25 or ≤0.80, and a magenta square marks a factor for which the complete 72-motif layer improved held-out prediction. Motif columns are grouped as shown for at least two factors or shown for one factor, based on either summit-association or held-out-prediction evidence; therefore, an open-ring-only cell need not improve prediction. An empty cell does not mean that the motif is absent, and display under at least two factors does not establish simultaneous binding to the same promoter. Summit association and conditional held-out prediction answer different questions. Neither the partner map nor the motif evidence establishes physical interaction, recruitment direction, or causation.

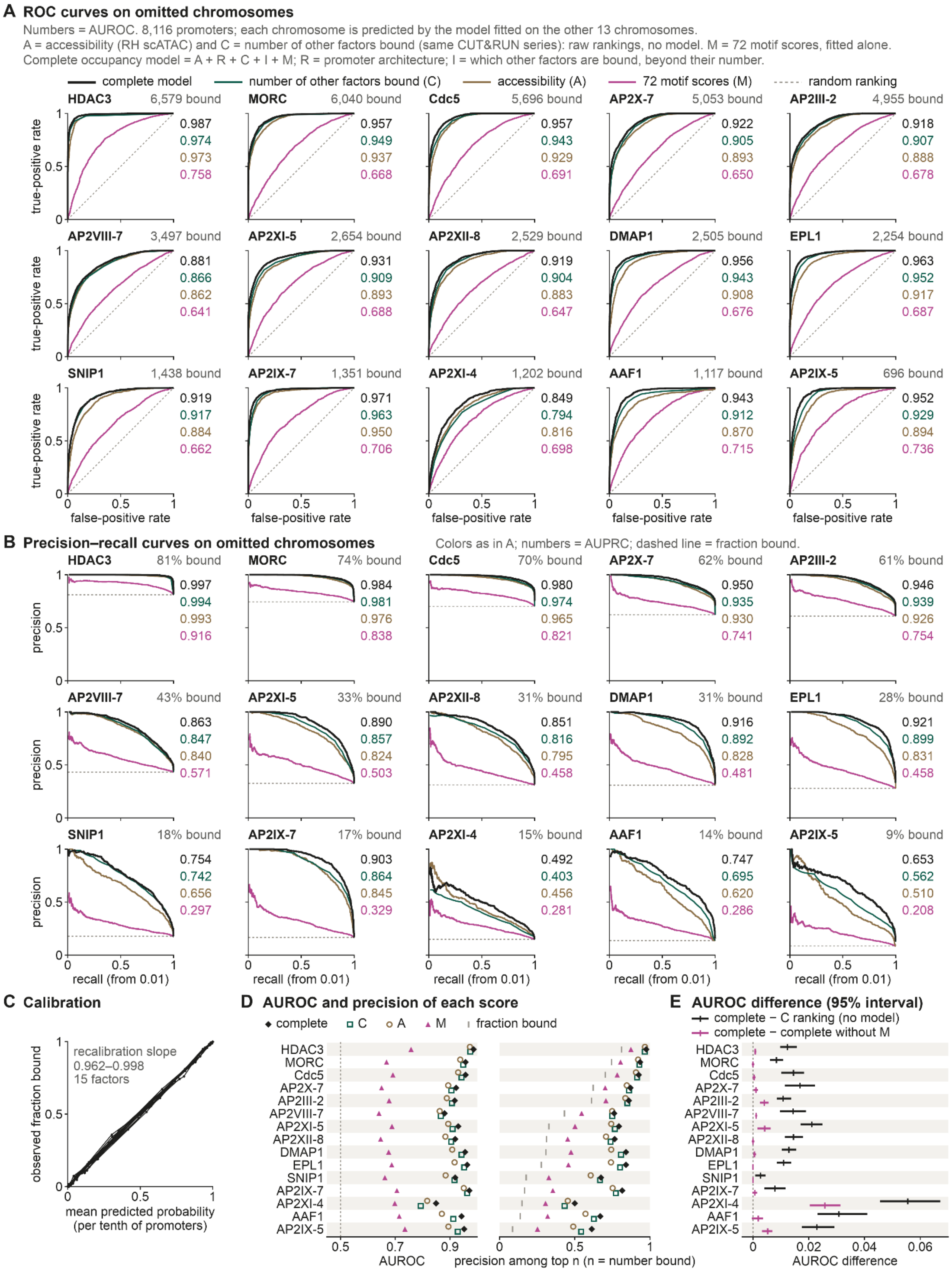


**Figure S13. Conditional predictive performance of the occupancy model on omitted chromosomes, related to Fig. 6.** AUROC and precision among the top *n* promoters (D) are the primary summary measures; every complete-model AUROC and precision-at-*n* value is shown beside ranking promoters by C alone.

**(A)** Receiver operating characteristic curves for the 15 target factors, ordered by the number of bound promoters. Each of the 14 chromosomes was predicted by the complete occupancy

model—accessibility (A), promoter architecture (R), number of other factors bound (C), identities of the other bound factors beyond their number (I), and 72 motif scores (M; **Fig. S12A**)—fitted and tuned on the other 13 chromosomes. Predictions were pooled across all 8,116 promoters; no model was refitted for this figure. Four scores are compared: the complete model; C and RH scATAC accessibility (A), each used directly as a ranking without fitting a model; and the 72 motif scores alone (M), fitted using the same leave-one-chromosome-out framework. AUROC is the probability that a bound promoter scores higher than an unbound promoter, with ties contributing one half (0.5 denotes random ranking). The complete model reached AUROC 0.849–0.987 (median 0.943), compared with 0.794–0.974 (median 0.917) for C, 0.816–0.973 (median 0.893) for A, and 0.641–0.758 (median 0.687) for M. Promoter architecture and motif scores together (R + M, not drawn) reached 0.680–0.782 (median 0.706), whereas promoter architecture alone (not drawn) reached 0.580–0.700 (median 0.618). Because C and I are derived from the other factors' CUT&RUN calls in the same experimental series, this is conditional occupancy prediction: the complete model cannot score a new factor or an unprofiled promoter without those measurements.
**(B)** Precision–recall curves for the same scores, drawn from recall 0.01 to the first point at recall 1. Numbers are average precision (AUPRC); the dashed line is the fraction of promoters bound, which is the expected AUPRC of a random ranking. Promoters with tied scores are evaluated as the expected value under random ordering within the tie. Complete-model AUPRC was 0.492–0.997, compared with bound fractions of 0.086–0.811.
**(C)** Calibration of the complete model. Promoters were divided into tenths according to predicted probability for each factor; points show the mean predicted probability and observed fraction bound in each tenth. Logistic recalibration of the observed outcome against the predicted log odds gave slopes of 0.962–0.998 and intercepts of −0.025 to 0.013; perfect calibration has slope 1 and intercept 0.
**(D)** AUROC (left) and precision among the top *n* promoters, where *n* is the number of bound promoters (right), for each score. Promoters tied at the cutoff are counted proportionally, and the faint tick marks the fraction bound expected under random ranking. A is displaced slightly above and C slightly below each row solely to keep nearly equal values visible. Complete-model precision among the top *n* was 0.500–0.975 (median 0.816), compared with 0.432–0.965 (median 0.773) for C alone.
**(E)** AUROC difference between the complete model and C alone (black), and between the complete model and the model without the 72 motif scores, A + R + C + I (magenta). Bars are 95% percentile intervals from the chromosome bootstrap used in **Fig. 6C-D**: the same 2,000 paired draws of 14 chromosomes resampled with replacement, conditional on the fitted models. The complete-model difference from C was 0.003–0.056 (median 0.014); its interval was above zero for all 15 factors, and the difference was largest for AP2XI-4 (0.056). Adding M to A + R + C + I produced a median AUROC difference of 0.0008, which was below 0.001 for 9 of 15 factors. The interval was above zero for 7 factors, with the largest difference for AP2XI-4 (0.026); all 7 were among the 10 factors whose 72-motif layer improved held-out log-likelihood in **Fig. S12A**. EPL1 had an interval below zero (−0.0001), and its motif layer did not pass the **Fig. S12A** test. AUROC measures promoter ordering, whereas held-out log-likelihood also reflects the sharpness of predicted probabilities; accordingly, AP2IX-7, AP2X-7, DMAP1, and HDAC3 passed the log-likelihood test despite AUROC differences below 0.001. The 72 motifs were selected using CUT&RUN data from all chromosomes (**Fig. 6A** and **Fig. S11A**), so omitted-chromosome evaluation does not cover motif selection and every score containing M may be optimistic. Without motif scores, A + R + C + I reached AUROC 0.823–0.986 (median 0.941); its difference from C was 0.003–0.030 (median 0.013), with the interval above zero for all 15 factors.

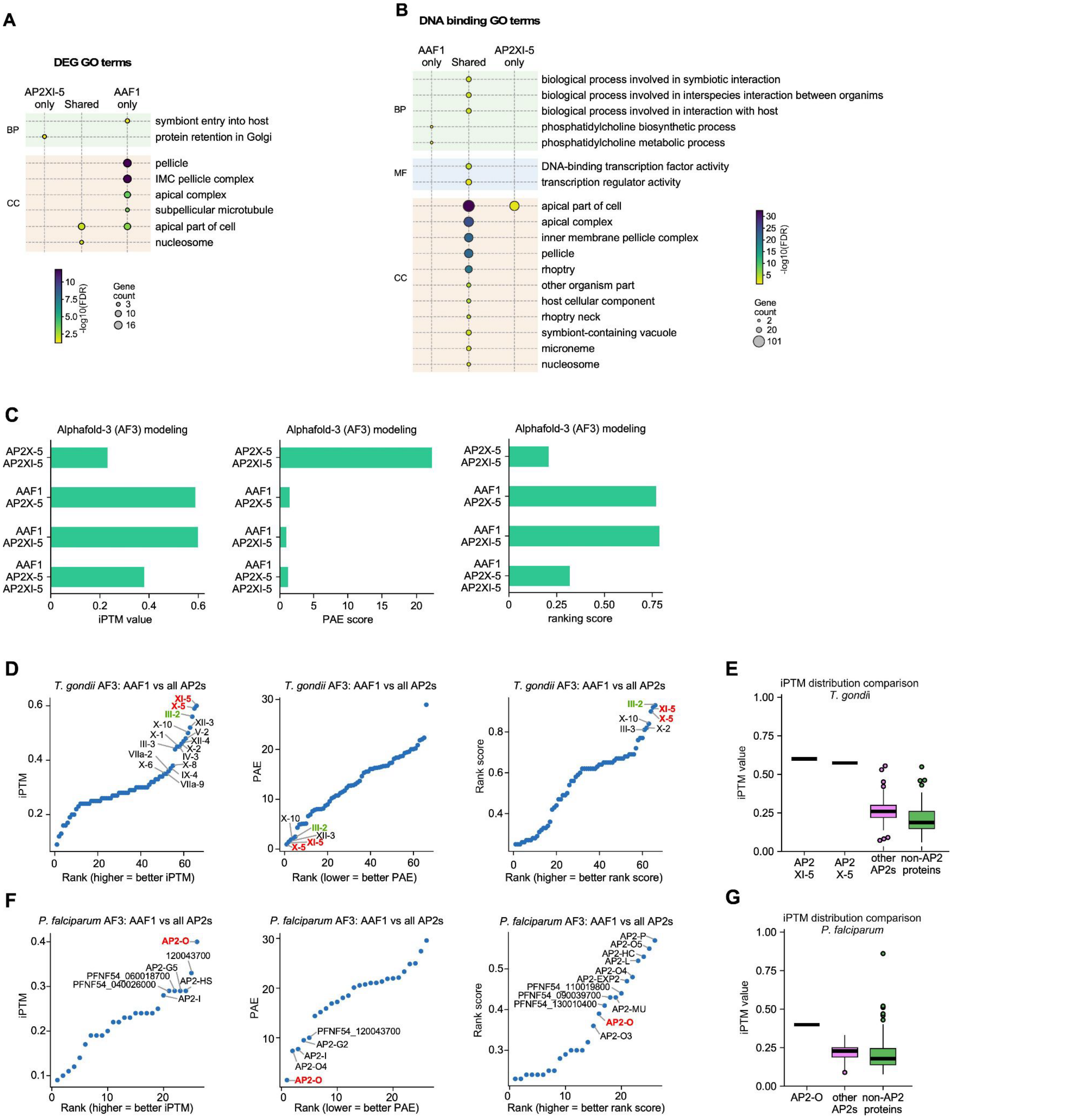


**Figure S14. AAF1 GRN GO-terms and AlphaFold-3 of AAF1 paired with all AP2 factors in *T. gondii* and *P. falciparum*.**

**(A)** GO term enrichment of differentially expressed genes (DEGs) upon AAF1 or AP2XI-5 depletion. Dot plots show enriched GO terms for DEGs shared between AAF1 and AP2XI-5 depletion and for DEGs unique to each factor, grouped by biological process (BP) and cellular component (CC). Fold enrichment is on the x-axis, dot size shows the number of genes annotated to each term, and color shows significance (-log10 FDR).

**(B)** GO term enrichment of direct target genes of AAF1 or AP2XI-5. Dot plot of GO terms for genes associated with AAF1 only, AP2XI-5 only, or both factors (shared), grouped by biological process (BP), molecular function (MF) and cellular component (CC). Dot size shows the number of genes per term, and color shows significance (-log10 FDR).

**(C)** AlphaFold 3 (AF3) modeling of TgAAF1 in complex with AP2X-5 and AP2XI-5. Interface predicted template modeling (iPTM) score, predicted aligned error (PAE) and ranking score for the indicated pairwise and three-way complexes (AP2X-5/AP2XI-5, AAF1/AP2X-5, AAF1/AP2XI-5 and AAF1/AP2X-5/AP2XI-5). Higher iPTM and ranking scores, and lower PAE indicate a more confident interface prediction.
**(D)** Ranking of *Tg*AAF1 interactions with all *T. gondii* AP2 factors. AF3 iPTM, PAE and ranking score for TgAAF1 paired with each *Tg*AP2 factor, ordered by rank (higher rank = better iPTM and ranking score; lower PAE = better). AP2X-5 and AP2XI-5 (red), and AP2III-2 (green) are highlighted, and the other top-ranking AP2 factors are labeled.
**(E)** AlphaFold3 iPTM comparison for *T. gondii* AAF1. Box plots of AF3 iPTM values for *Tg*AAF1 paired with AP2XI-5 and AP2X-5, compared with those for the 54 other *Tg*AP2 factors and 100 randomly selected non-AP2 *T. gondii* proteins. The thick line is the median, the box spans the 25th-75th percentiles, and whiskers extend to the most extreme observations within 1.5 × the interquartile range. Outliers are plotted individually. For AP2XI-5 and AP2X-5, n = 1, so the box and whiskers collapse into a single line.
**(F)** Ranking of *Pf*AAF1 interactions with all *P. falciparum* AP2 factors. AF3 iPTM, PAE and ranking score for *Pf*AAF1 paired with each *Pf*AP2 factor, ordered by rank (higher rank is better interaction for iPTM and ranking score; lower ranks is better interaction for PAE). AP2-O (red) is highlighted, and the other top-ranking AP2 factors are labeled.
**(G)** AlphaFold3 iPTM comparison for *P. falciparum* AAF1. Box plots of AF3 iPTM values for *Pf*AAF1 paired with AP2-O, compared with those for the 26 other *Pf*AP2 factors and 100 randomly selected non-AP2 *P. falciparum* proteins. Box plot elements are as in (E): median (thick line), 25th-75th percentiles (box), whiskers to the most extreme observations within 1.5 × IQR, and individually plotted outliers. For AP2-O, n = 1, so the box and whiskers collapse into a single line.

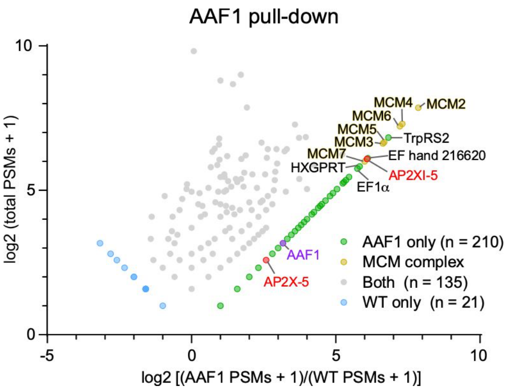


**Figure S15. AAF1 pull down.** The average of two biological replicates, with three technical replicates each, is plotted. The wild type control is the RHΔ*Ku80* parasite strain. PSM: peptide spectrum match. EF hand 216620 is a hypothetical protein where the numeral refers to the ToxoDB accession number (TGGT1_216620).

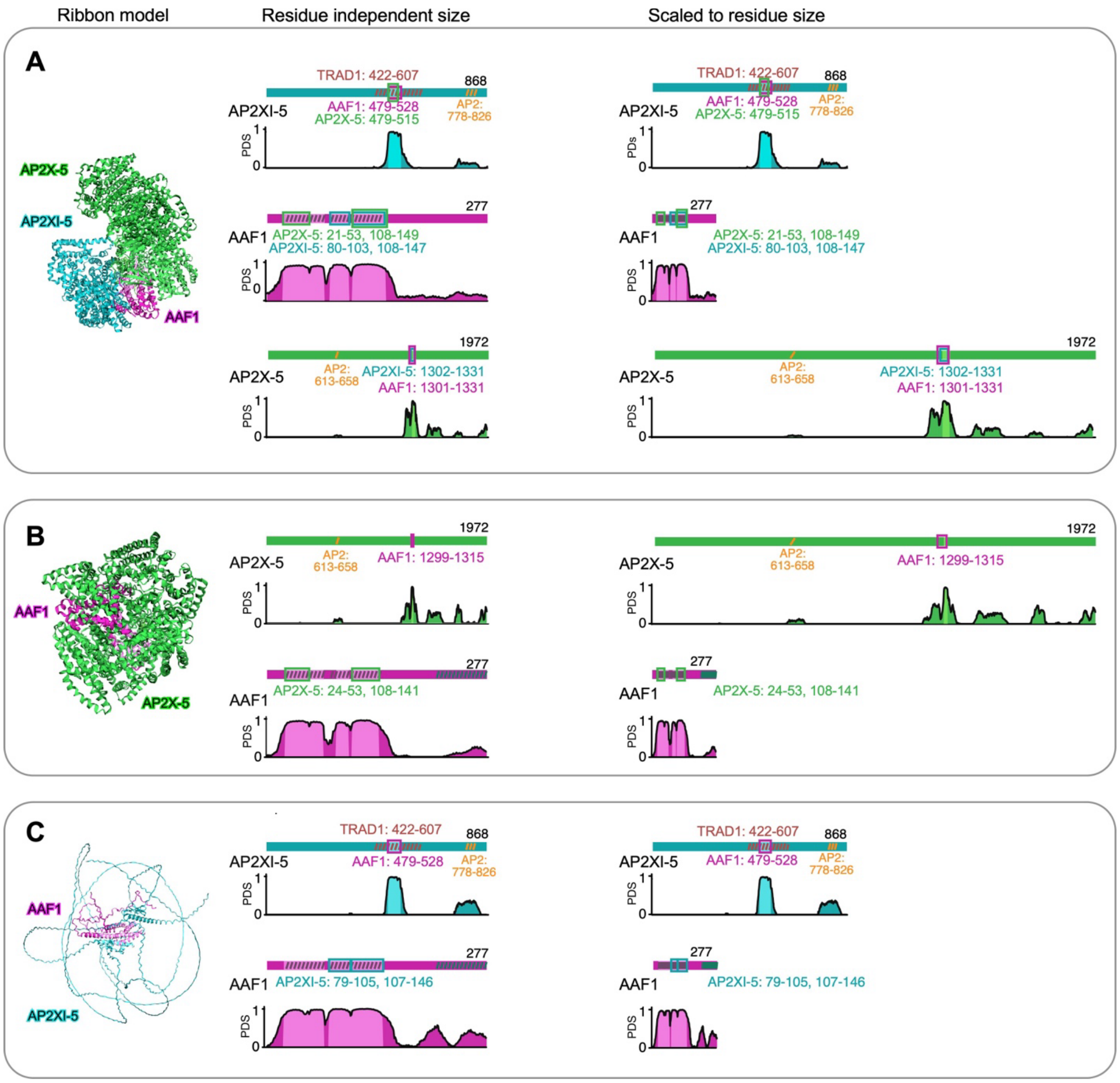


**Figure S16. Alphafold-3 modeled interaction domains in the AAF1-AP2 complexes.** Predicted structures of the AAF1-AP2X-5-AP2XI-5 tripartite complex (A) and the AAF1-AP2X-5 (B) and AAF1-AP2XI-5 (C) bipartite complexes. Left: ribbon models of each complex, with AP2X-5 in green, AP2XI-5 in cyan and AAF1 in magenta. Middle: domain maps of each protein drawn at a residue-independent size, so each protein has the same length. Right: the same maps scaled to residue number, with total lengths indicated (AP2XI-5, 868; AAF1, 277; AP2X-5, 1972 residues). Under each map, the interaction confidence profile across the protein is shown, and light shading marks the region of the protein predicted to interact with a partner (PAE-derived score, PDS; minimum inter-chain PAE ≤ 6 Å, indicating a high-confidence predicted interaction). Colored outlines mark the interacting regions, with residue ranges listed and the outline color matching the partner protein. Diagonal hatching denotes annotated or conserved domains, including the AP2 domains (orange) and the region labeled TRAD1 (brown). Gaps of ≤2 residues are filled for visualization only. Boxes and residue ranges reflect the exact analyzed regions.

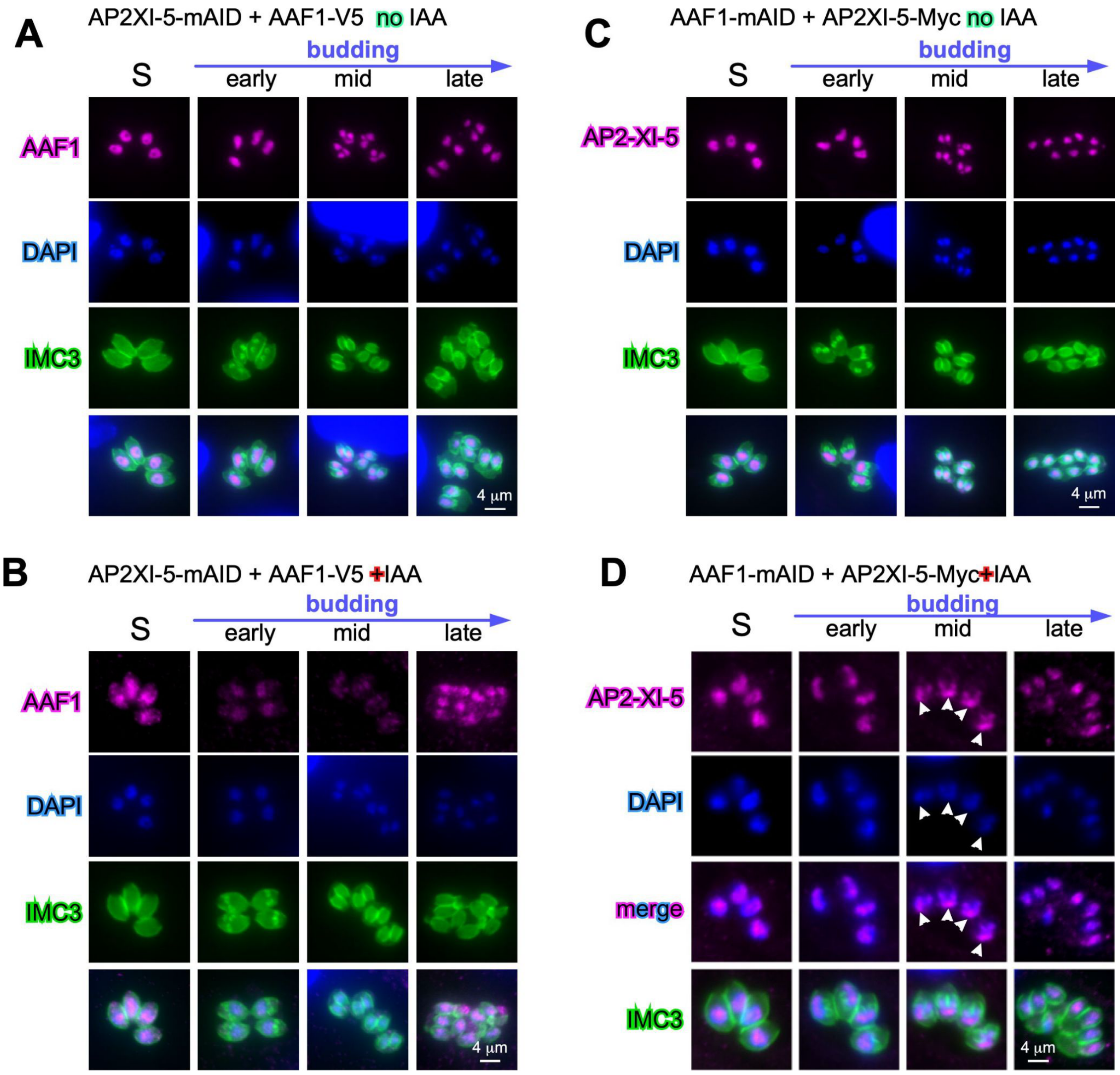


**Figure S17. All IFA panels corresponding with Fig. 7H-I.**
**(A and B)** AP2XI-5-cKD wherein AAF1 is simultaneously tagged with a Myc tag to monitor AAF1 localization upon AP2XI-5 depletion (**B**).
**(C and D)** AAF1-cKD wherein AP2XI-5 and AAF1 are simultaneously tagged with a 10xMyc or 5xV5 tag, respectively, to monitor AP2XI-5 localization upon AP2XI-5 or AAF1 depletion, respectively. Arrowheads mark AP2XI-5 accumulation at the trailing end of the splitting nucleus, where there is very little DNA as seen by DAPI.

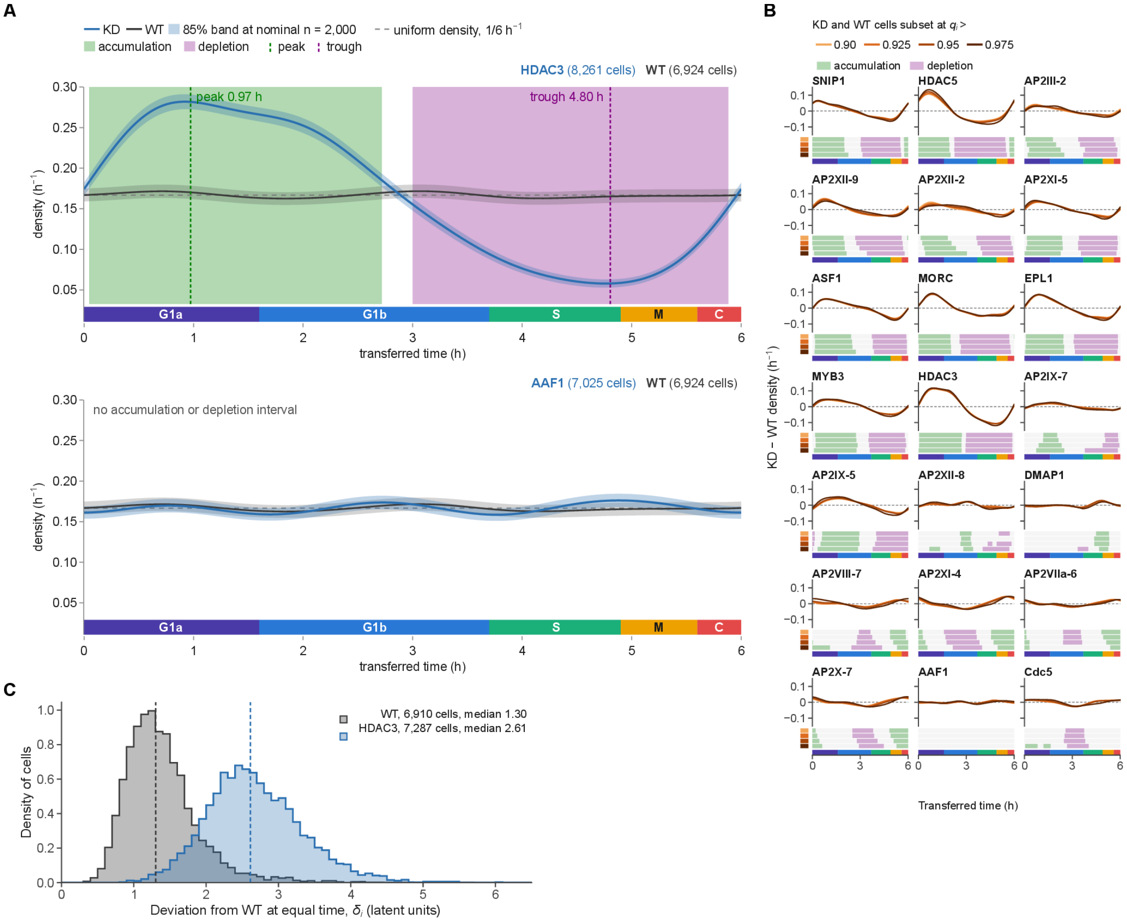


**Figure S18. Cell-cycle density and deviation from WT.**

**(A)** Cell density along transferred time for knockdown (blue) and WT (black). Top: HDAC3 (8,261 cells). Bottom: AAF1 (7,025 cells). WT has 6,924 cells. Bands show the 85% interval at a nominal n = 2,000, and the dashed line is uniform density (1/6 $h^{-1}$). Shaded intervals mark accumulation (green) and depletion (purple), with their peak and trough. AAF1 has no accumulation or depletion interval.

**(B)** Threshold sensitivity. KD − WT density for all 21 factors, with KD and WT cells subset at matched quality-score cutoffs (q > 0.90, 0.925, 0.95 and 0.975). Below each curve, one row per cutoff gives the accumulation and depletion calls; the phase strip is beneath.

**(C)** Deviation from WT at equal time. δ_i is the distance in the 10-dimensional latent space between cell i and the WT reference trajectory at its transferred time. Histograms show WT cells against the WT trajectory (6,910 cells, median 1.30) and HDAC3 cells (7,287 cells, median 2.61), with medians dashed.

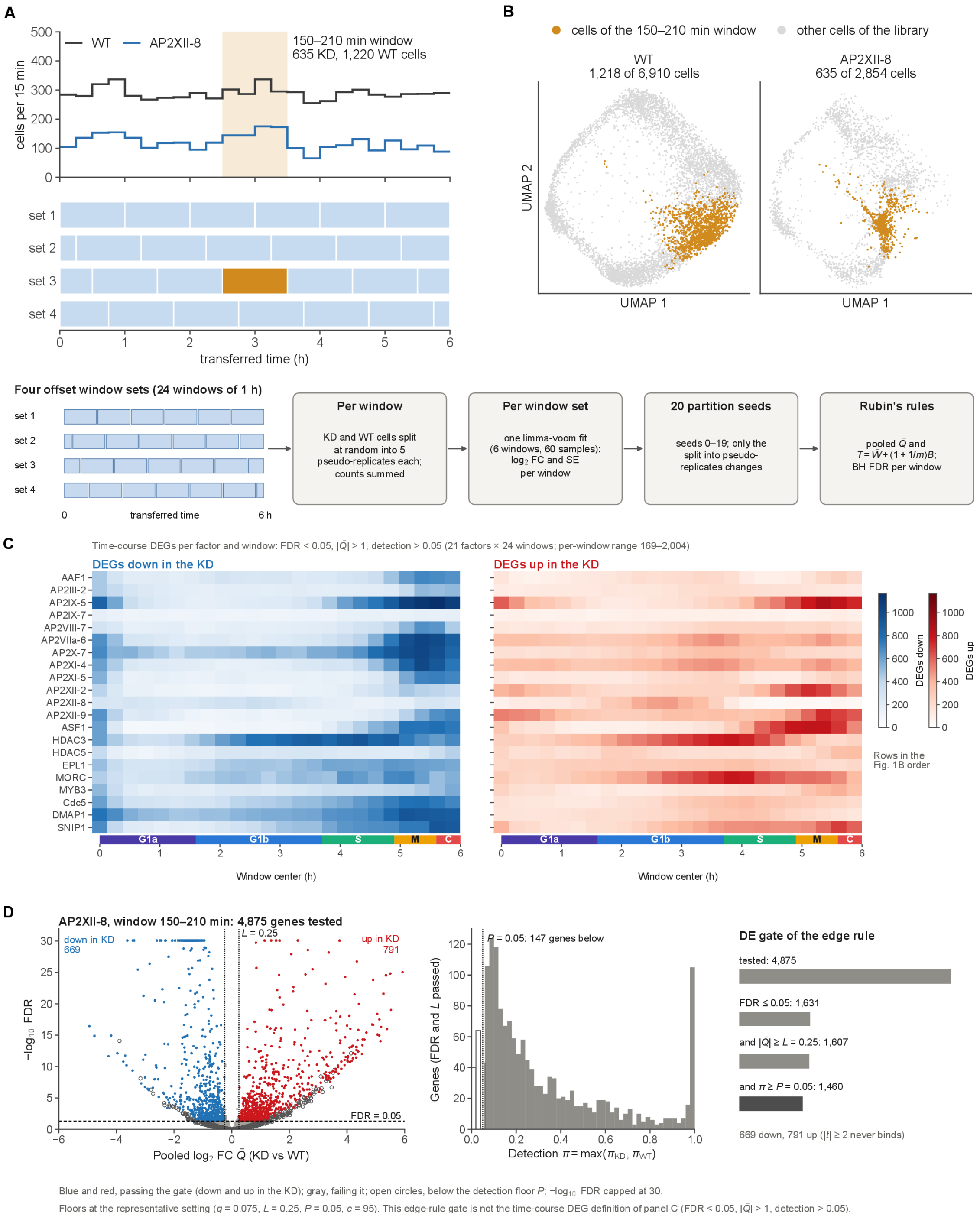

**Figure S19. Differential expression within time windows.**
**(A)** Time windows, shown for WT and AP2XII-8. Top: cells per 15 min of transferred time; the 150–210 min window (635 KD, 1,220 WT cells) is shaded. Bottom: the four window sets, each tiling the cycle with 1 h windows, offset by 15 min; together they give 24 sliding windows. The highlighted window belongs to set 3.
**(B)** Pseudo-replicates, model and pooling. Top: cells of the 150–210 min window (orange) among the other cells of each library, in WT (1,218 of 6,910 cells) and in AP2XII-8 (635 of 2,854). Bottom: **per window**, KD and WT cells are split at random into five pseudo-replicates each and their

counts summed; **per window set**, one limma-voom fit (6 windows, 60 samples) gives log2 fold changes and standard errors per window; **20 partition seeds** (0–19) change only the split into pseudo-replicates; **Rubin's rules** pool the seeds to a mean log2 fold change $\bar{Q}$ and a total variance $T = \bar{W} + (1 + 1/m)B$, with Benjamini–Hochberg FDR per window.
**(C)** Time-course DEGs: counts per factor and window, down-regulated (blue) and up-regulated (red) in the knockdown. A DEG has FDR < 0.05, |log2FC| > 1 and detection > 0.05. Rows follow the order of **Fig. 1B**. Per factor and window, 169 to 2,004 DEGs.
**(D)** Use in the networks: the DE gate of the edge rule in one window (AP2XII-8, 150–210 min, 4,875 genes tested). Left: volcano plot of −log10 FDR (capped at 30) against $\bar{Q}$. The effect-size floor $L = 0.25$ and FDR = 0.05 are marked, and open circles fall below the detection floor. Middle: detection $\pi = \max(\pi\_KD, \pi\_WT)$ of the genes that pass FDR and L, with the floor $P = 0.05$ (147 genes below). Right: genes remaining after each condition, 4,875 → 1,631 → 1,607 → 1,460 (669 down, 791 up). Floors are at the representative setting ($q = 0.075$, $L = 0.25$, $P = 0.05$, $c = 95$). This gate is not the time-course DEG definition of panel C.

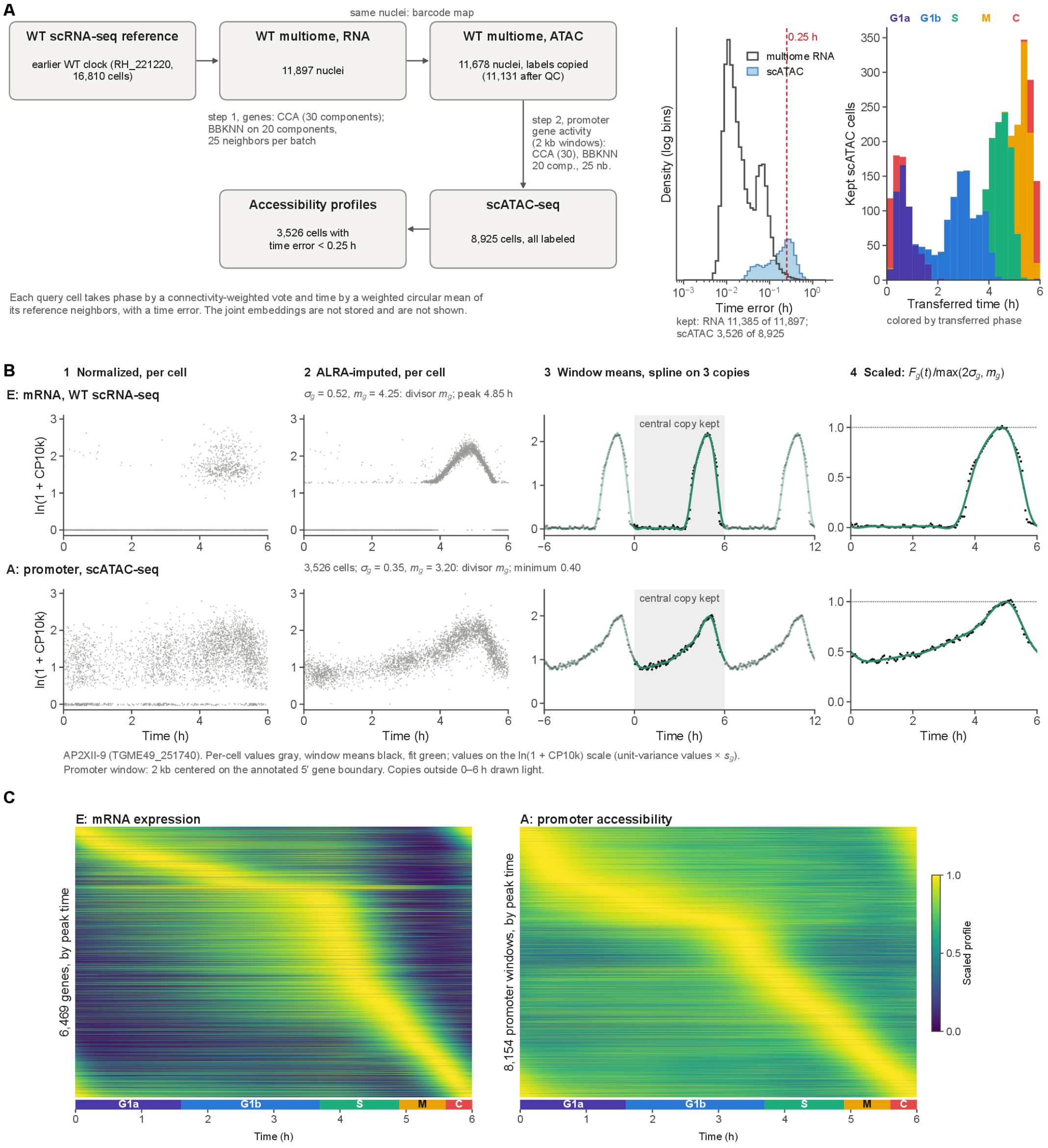
A
same nuclei: barcode map
WT scRNA-seq reference
earlier WT clock (RH_221220, 16,810 cells)
WT multiome, RNA
11,897 nuclei
WT multiome, ATAC
11,678 nuclei, labels copied (11,131 after QC)
step 1, genes: CCA (30 components); BBKNN on 20 components, 25 neighbors per batch
step 2, promoter gene activity (2 kb windows): CCA (30), BBKNN 20 comp., 25 nb.
Accessibility profiles
3,526 cells with time error < 0.25 h
scATAC-seq
8,925 cells, all labeled
Each query cell takes phase by a connectivity-weighted vote and time by a weighted circular mean of its reference neighbors, with a time error. The joint embeddings are not stored and are not shown.
0.25 h
multiome RNA
scATAC
Density (log bins)
Time error (h)
kept: RNA 11,385 of 11,897; scATAC 3,526 of 8,925
G1a G1b S M C
Kept scATAC cells
Transferred time (h)
colored by transferred phase
B
1 Normalized, per cell
2 ALRA-imputed, per cell
3 Window means, spline on 3 copies
4 Scaled: $F_g(t)/\max(2\sigma_g, m_g)$
E: mRNA, WT scRNA-seq
$\sigma_g$ = 0.52, $m_g$ = 4.25: divisor $m_g$; peak 4.85 h
ln(1 + CP10k)
central copy kept
A: promoter, scATAC-seq
3,526 cells; $\sigma_g$ = 0.35, $m_g$ = 3.20: divisor $m_g$; minimum 0.40
Time (h)
AP2XII-9 (TGME49_251740). Per-cell values gray, window means black, fit green; values on the ln(1 + CP10k) scale (unit-variance values × $s_g$).
Promoter window: 2 kb centered on the annotated 5′ gene boundary. Copies outside 0–6 h drawn light.
C
E: mRNA expression
A: promoter accessibility
6,469 genes, by peak time
8,154 promoter windows, by peak time
Scaled profile
G1a G1b S M C
Time (h)

#### Figure S20 (continued)

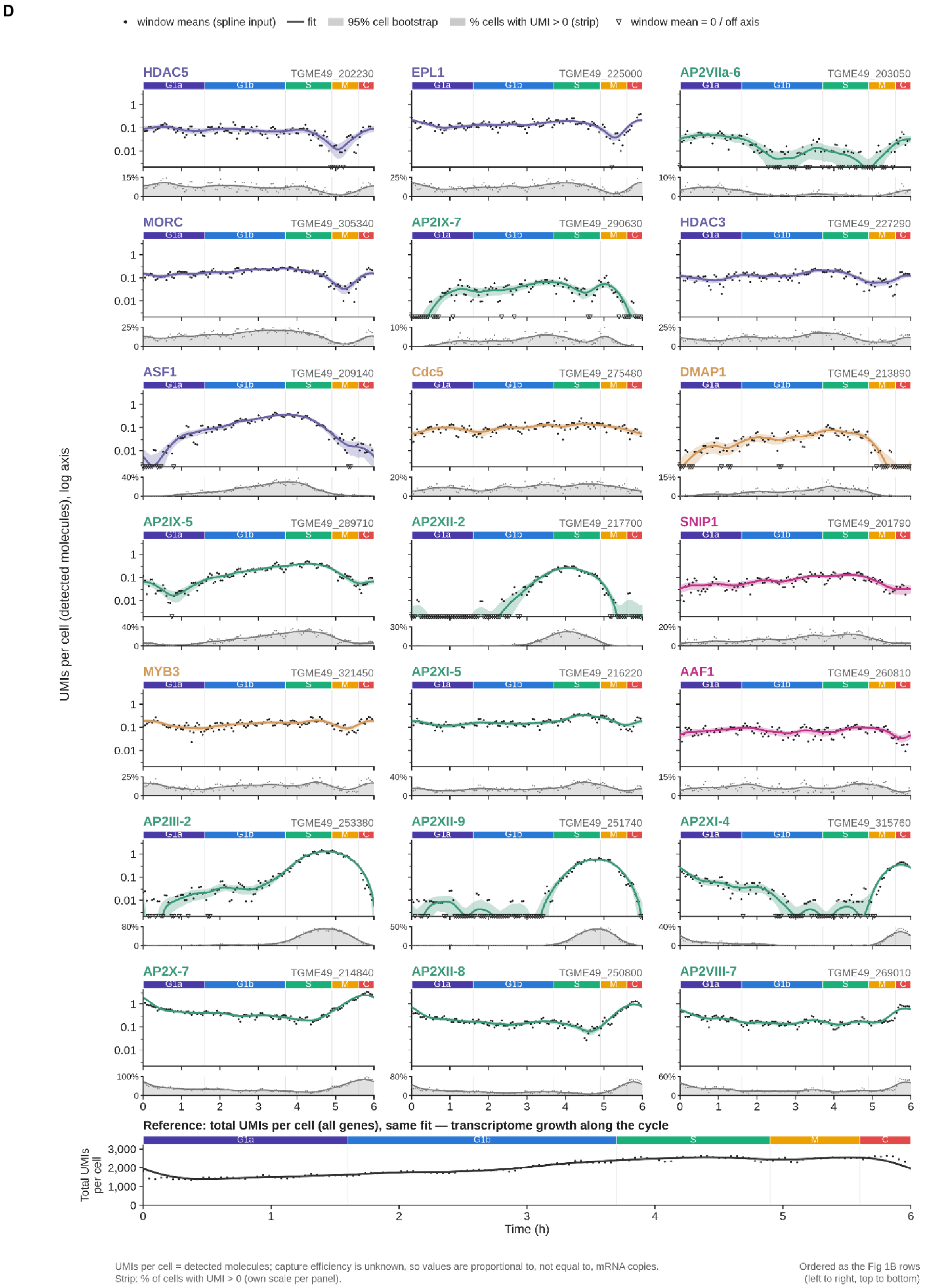


**Figure S20. scATAC processing; expression and accessibility profiles.**

**(A)** Transfer of clock time and phase to the scATAC cells. **Step 1.** WT scRNA-seq reference (earlier WT clock RH_221220, 16,810 cells) to the RNA of the WT multiome (11,897 nuclei), on genes. CCA (30 components), then BBKNN on 20 components with 25 neighbors per batch. A PCA graph of the query cells then propagates the labels. **Step 2.** Labels are copied to the 11,678 multiome ATAC nuclei through the barcode map. These nuclei serve as the reference (11,131 after QC) for the 8,925 scATAC cells, using promoter gene activity: counts in a 2 kb window centered on the annotated 5′ gene boundary. CCA (30 components), then BBKNN on 20 components with 25 neighbors per dataset. **Labels and filter.** Phase comes from a connectivity-weighted vote and time from a weighted circular mean of reference neighbors, with a time error. Cells with time error < 0.25 h (3,526 of 8,925) enter the accessibility profiles. **Plots.** Middle: time-error distributions for the multiome RNA nuclei and the scATAC cells, with the 0.25 h cut; 11,385 of 11,897 RNA nuclei are below it. Right: transferred time of the 3,526 kept cells, stacked by transferred phase. The joint embeddings are not stored and are not shown.

**(B)** One gene from cells to its profiles: AP2XII-9 (TGME49_251740). Top row: mRNA (E), WT scRNA-seq. Bottom row: promoter accessibility (A), scATAC-seq. Columns:

1. normalized values per cell;
2. ALRA-imputed values (rank 16 for E, 29 for A);
3. window means with the periodic spline fitted to three tiled copies of the cycle, keeping the central copy;
4. the scaled profile $F_g(t) / \max(2\sigma_g, m_g)$.

Values are on the $\ln(1 + \text{CP10k})$ scale. For E, $\sigma_g = 0.52$ and $m_g = 4.25$ (divisor $m_g$; peak 4.85 h). For A, $\sigma_g = 0.35$ and $m_g = 3.20$ (divisor $m_g$; minimum 0.40).

**(C)** Scaled profiles of all fitted genes, ordered by peak time. E: 6,469 genes. A: 8,154 promoter windows. The phase strip is below.

**(D)** Absolute mRNA profiles (UMIs per cell, log axis) of the 21 factors, with window means, fit, 95% cell bootstrap and the share of cells with UMI > 0. The bottom row gives total UMIs per cell (all genes) with the same fit. Factors are ordered as the rows of **Fig. 1B**. UMIs per cell are detected molecules, so values are proportional to, not equal to, mRNA copies.

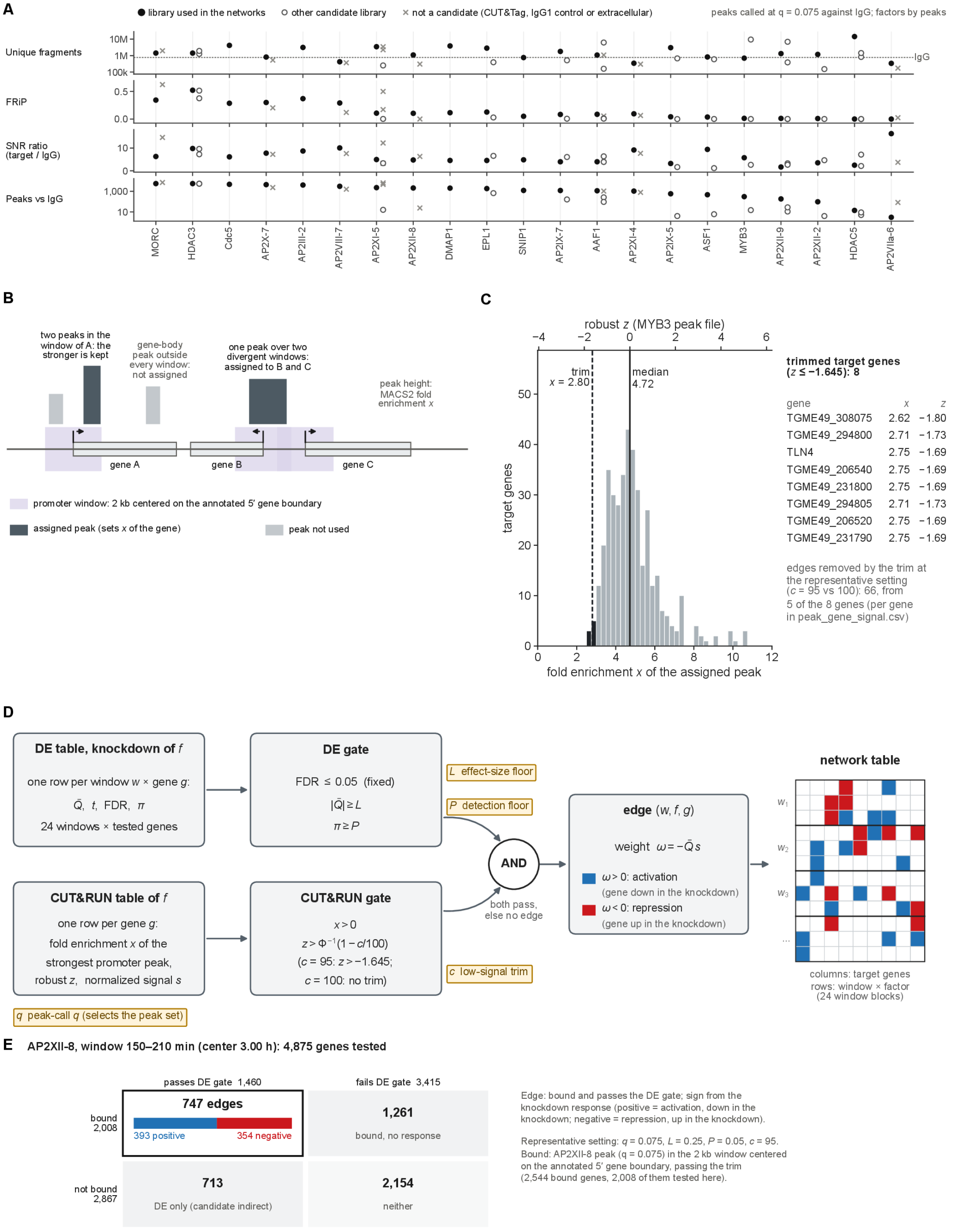


**Figure S21. CUT&RUN processing, peak calling, peak–gene assignment and the edge rule.** **(A)** Peak calling and library selection. For each CUT&RUN library, with peaks called against IgG at q < 0.075: unique fragments, FRiP, SNR ratio (target/IgG density) and the number of peaks. Filled black: the library used in the networks (one per factor). Open: other candidate libraries. Crosses: libraries that are not candidates (CUT&Tag, IgG1 control or extracellular). Factors are ordered by peak count; the dashed line marks the IgG fragment count (772,541).

**(B)** Peak–gene assignment. Each gene has a promoter window of 2 kb centered on its annotated

5′ gene boundary. The strongest peak in the window sets the gene's fold enrichment x. A gene-body peak outside every window is not assigned. A peak over two divergent windows is assigned to both genes.

**(C)** Per-file robust z and the low-signal trim, shown for MYB3. Histogram of the fold enrichment x of the assigned peaks, with robust z on the top axis, the median, and the trim at $z = -1.645$ ($c = 95$). The 8 trimmed target genes are listed, and 66 edges are removed by the trim at the representative setting.

**(D)** Edge rule. The knockdown DE table (per window and gene: $\bar{Q}$, t, FDR, π) passes the DE gate: FDR ≤ 0.05, $|\bar{Q}| \geq L$, $\pi \geq P$. The CUT&RUN table (per gene: x, robust z, normalized signal s) passes the CUT&RUN gate: $x > 0$ and $z > \Phi^{-1}(1 - c/100)$. A gene that passes both gates in window w gets an edge (w, f, g) with weight $\omega = -\bar{Q}\,s$. $\omega > 0$ is activation (gene down in the knockdown); $\omega < 0$ is repression (gene up in the knockdown).

**(E)** The rule in one window: AP2XII-8, 150–210 min, at the representative setting ($q = 0.075$, $L = 0.25$, $P = 0.05$, $c = 95$). Of 4,875 genes tested, 2,008 are bound and 1,460 pass the DE gate. The 747 genes that are both bound and DE become edges (393 positive, 354 negative). Bound but not DE: 1,261. DE but not bound: 713 (candidate indirect targets). Neither: 2,154.

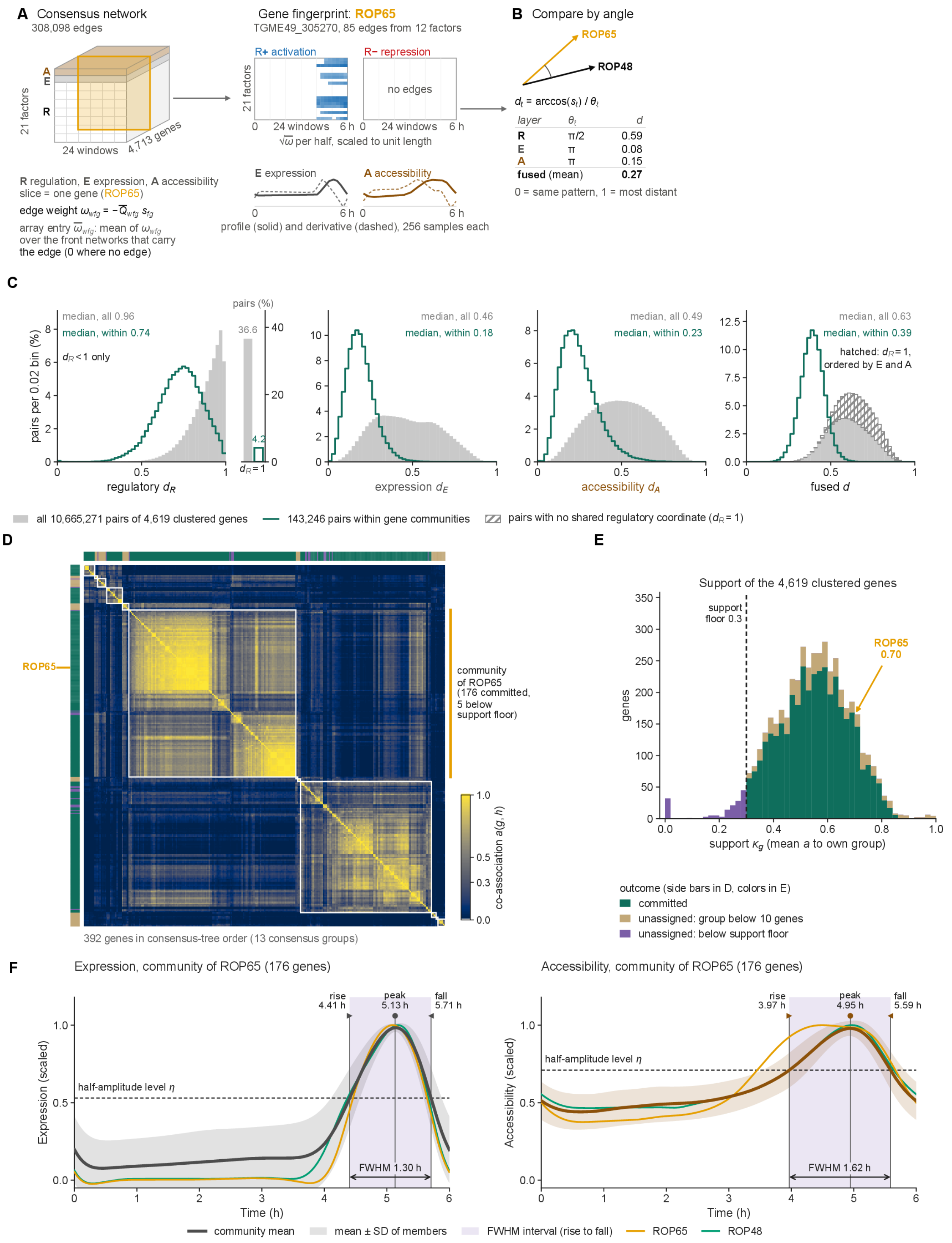


**Figure S22. Ensemble clustering and gene communities.**

**(A)** Gene layers. Left: the consensus network as a factor × window × gene array (308,098 edges; R regulation, E expression, A accessibility). Each edge weight is ω = −Q̄ s, and each array entry is the mean of ω over the front networks that carry the edge (0 where there is no edge). Right: the regulatory fingerprint of ROP65 (TGME49_305270; 85 edges from 12 factors), split into activation (R+) and repression (R−) halves, with √ω per half scaled to unit length. Below are its E and A profiles and their derivatives (256 samples each).

**(B)** Fused distance. The per-layer distance is the angle between two genes' layer vectors over the layer's maximum angle $\theta_\ell$ ($\pi/2$ for R, $\pi$ for E and A). ROP65 against ROP48: R 0.59, E 0.08, A 0.15; fused distance (mean) 0.27.
**(C)** Distance distributions for all 10,665,271 pairs of the 4,619 clustered genes (gray) and the 143,246 pairs within gene communities (green), per layer and fused. A side bar shows the $d_R = 1$ mass: pairs with no shared regulatory coordinate, 36.6% of all pairs and 4.2% of within-community pairs. In the fused panel these pairs are hatched.
**(D)** Ensemble and consensus cut: co-association a(g, h) over 120 replicates for the 392 genes of the 13 consensus groups around the ROP65 community, in consensus-tree order. The ROP65 community has 176 committed genes, and 5 more fall below the support floor. Side bars give each gene's outcome.
**(E)** Commitment: support $\kappa_g$ (mean co-association with the group of the gene) of the 4,619 clustered genes, colored by outcome (committed; unassigned because the group has fewer than 10 genes; unassigned because support is below the 0.3 floor). ROP65 has support 0.70.
**(F)** Communities of the consensus network: expression (left) and accessibility (right) profiles of the ROP65 community (176 genes). The panels show the community mean, the member mean ± SD, ROP65 and ROP48, the half-amplitude level $\eta$, and the rise, peak, fall and FWHM descriptors. Expression: rise 4.41 h, peak 5.13 h, fall 5.71 h, FWHM 1.30 h. Accessibility: rise 3.97 h, peak 4.95 h, fall 5.59 h, FWHM 1.62 h.

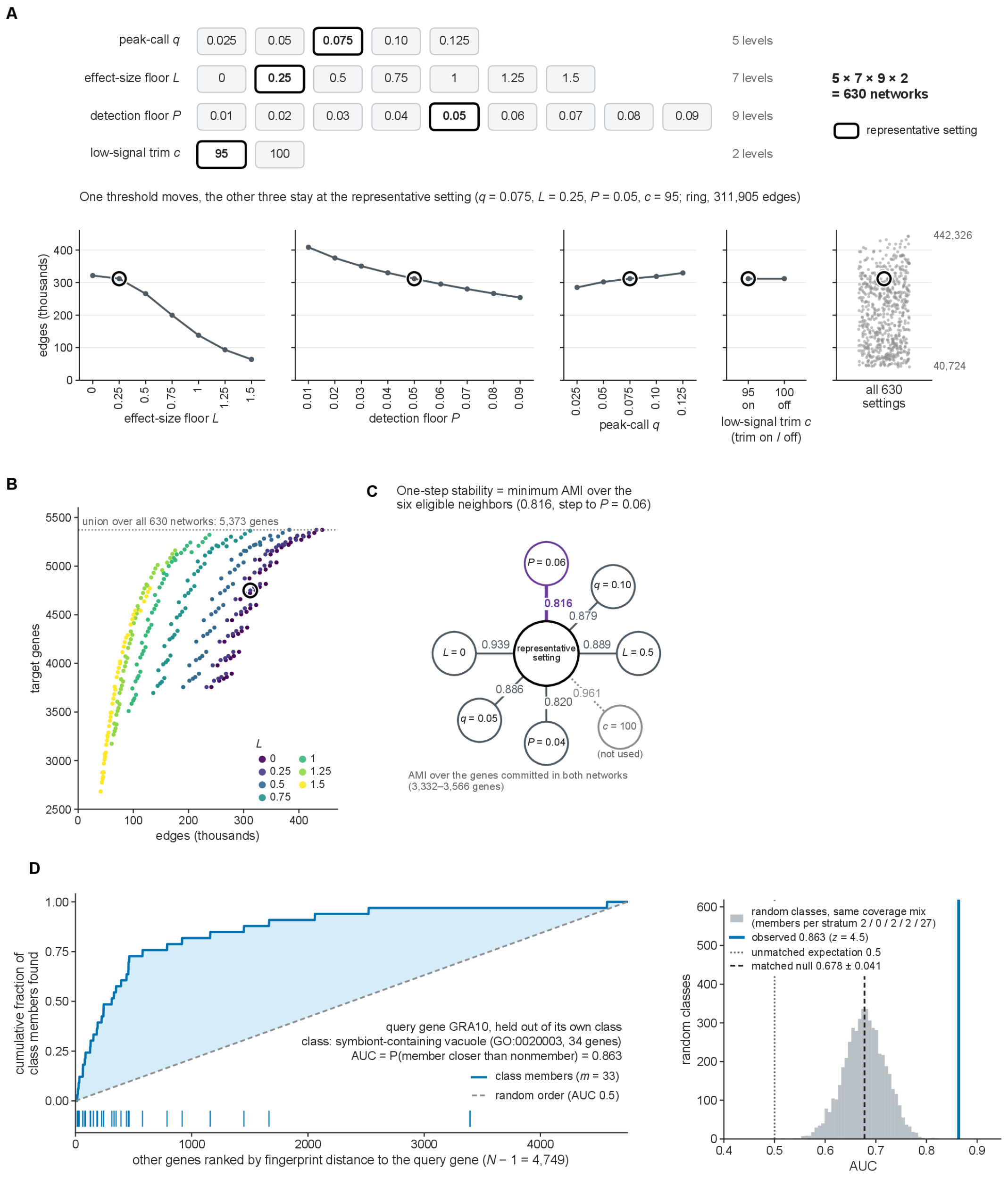
A
peak-call q
0.025
0.05
0.075
0.10
0.125
5 levels
effect-size floor L
0
0.25
0.5
0.75
1
1.25
1.5
7 levels
detection floor P
0.01
0.02
0.03
0.04
0.05
0.06
0.07
0.08
0.09
9 levels
low-signal trim c
95
100
2 levels
5 × 7 × 9 × 2
= 630 networks
representative setting
One threshold moves, the other three stay at the representative setting (q = 0.075, L = 0.25, P = 0.05, c = 95; ring, 311,905 edges)
edges (thousands)
effect-size floor L
detection floor P
peak-call q
95 on
100 off
low-signal trim c (trim on / off)
all 630 settings
442,326
40,724
B
union over all 630 networks: 5,373 genes
target genes
edges (thousands)
L
0
0.25
0.5
0.75
1
1.25
1.5
C
One-step stability = minimum AMI over the six eligible neighbors (0.816, step to P = 0.06)
P = 0.06
0.816
q = 0.10
0.879
L = 0
0.939
representative setting
0.889
L = 0.5
0.886
0.961
0.820
q = 0.05
P = 0.04
c = 100
(not used)
AMI over the genes committed in both networks (3,332–3,566 genes)
D
cumulative fraction of class members found
query gene GRA10, held out of its own class
class: symbiont-containing vacuole (GO:0020003, 34 genes)
AUC = P(member closer than nonmember) = 0.863
class members (m = 33)
random order (AUC 0.5)
other genes ranked by fingerprint distance to the query gene (N − 1 = 4,749)
random classes, same coverage mix (members per stratum 2 / 0 / 2 / 2 / 27)
observed 0.863 (z = 4.5)
unmatched expectation 0.5
matched null 0.678 ± 0.041
random classes
AUC

#### Figure S23 (continued)

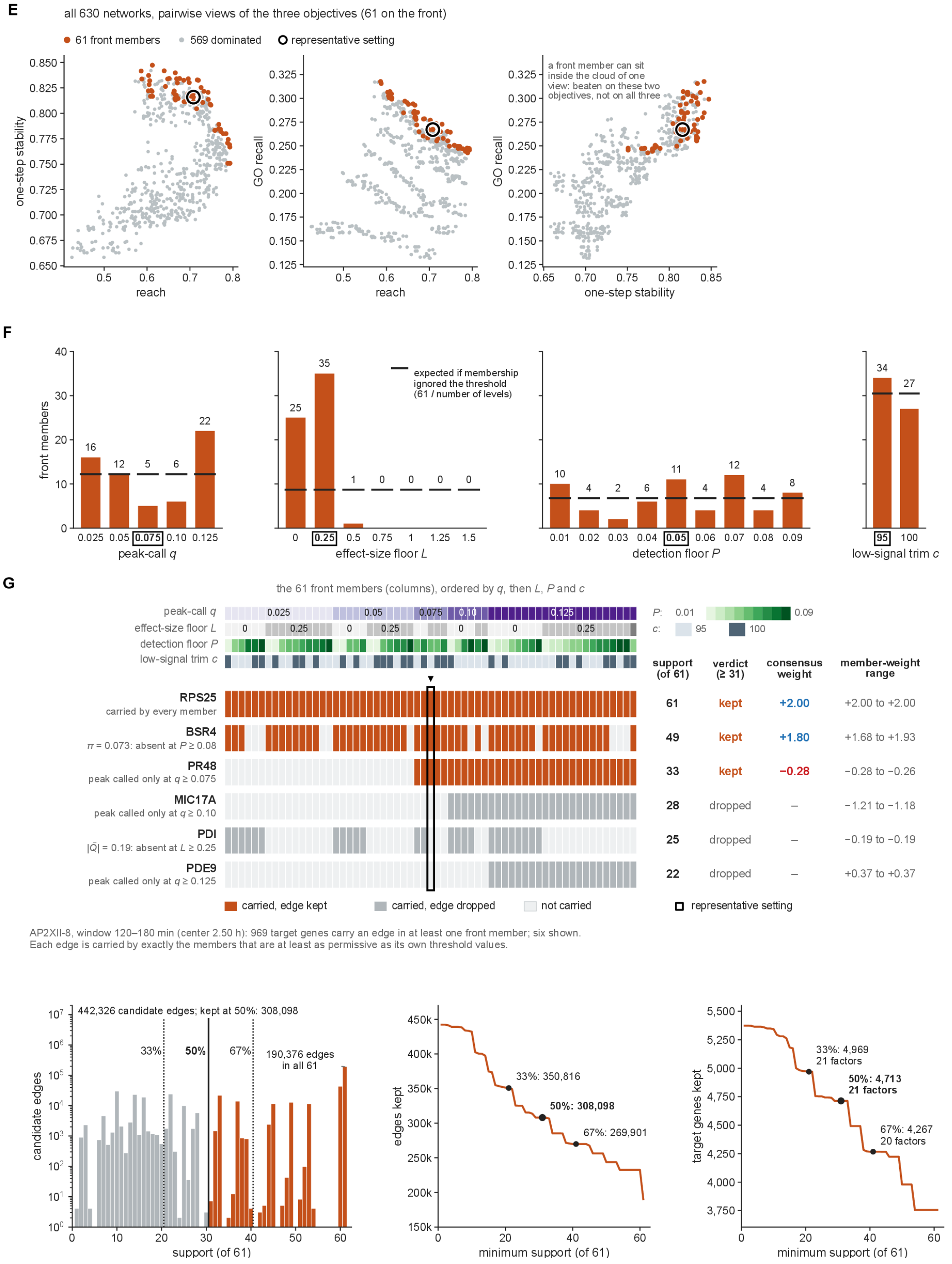


**Figure S23. Parameter sweep and consensus networks.**

**(A)** Parameter grid: peak-call q (5 levels), effect-size floor L (7), detection floor P (9) and low-signal trim c (2), giving 5 × 7 × 9 × 2 = 630 networks. Black outline: the representative setting (q = 0.075, L = 0.25, P = 0.05, c = 95). Below, network size as one threshold moves while the other three stay at the representative setting (ring: 311,905 edges), next to all 630 settings (40,724 to 442,326 edges).

**(B)** Target genes against edges for all 630 networks, colored by L. The dotted line is the union over all networks (5,373 genes); the ring is the representative setting.

**(C)** One-step stability: the minimum adjusted mutual information (AMI) between the representative network and its six eligible one-step neighbors (0.816, the step to $P = 0.06$). AMI is computed over the genes committed in both networks (3,332–3,566 genes).

**(D)** GO recall. Left: query gene GRA10, held out of its class (symbiont-containing vacuole, GO:0020003, 34 genes). Other genes are ranked by fingerprint distance to the query, and the curve gives the cumulative fraction of class members found; AUC = 0.863. Right: AUCs of random classes with the same coverage mix (matched null 0.678 ± 0.041; observed $z = 4.5$).

**(E)** Pareto front. All 630 networks in pairwise views of the three objectives (reach, one-step stability, GO recall): 61 front members (orange), 569 dominated, and the representative setting (black ring).

**(F)** Front members per parameter level. The black lines are the counts expected if membership ignored the threshold (61 / number of levels).

**(G)** Consensus network. Top: vote example for AP2XII-8 in the 120–180 min window. The 61 front members (columns) are ordered by q, L, P and c. Six target genes show how an edge is carried by exactly the members at least as permissive as its threshold values, and kept when at least 31 of 61 carry it. Bottom: edges by support (log scale) and edges and target genes kept as a function of minimum support, with the 33%, 50% and 67% thresholds. The 50% threshold keeps 308,098 edges and 4,713 genes from all 21 factors.

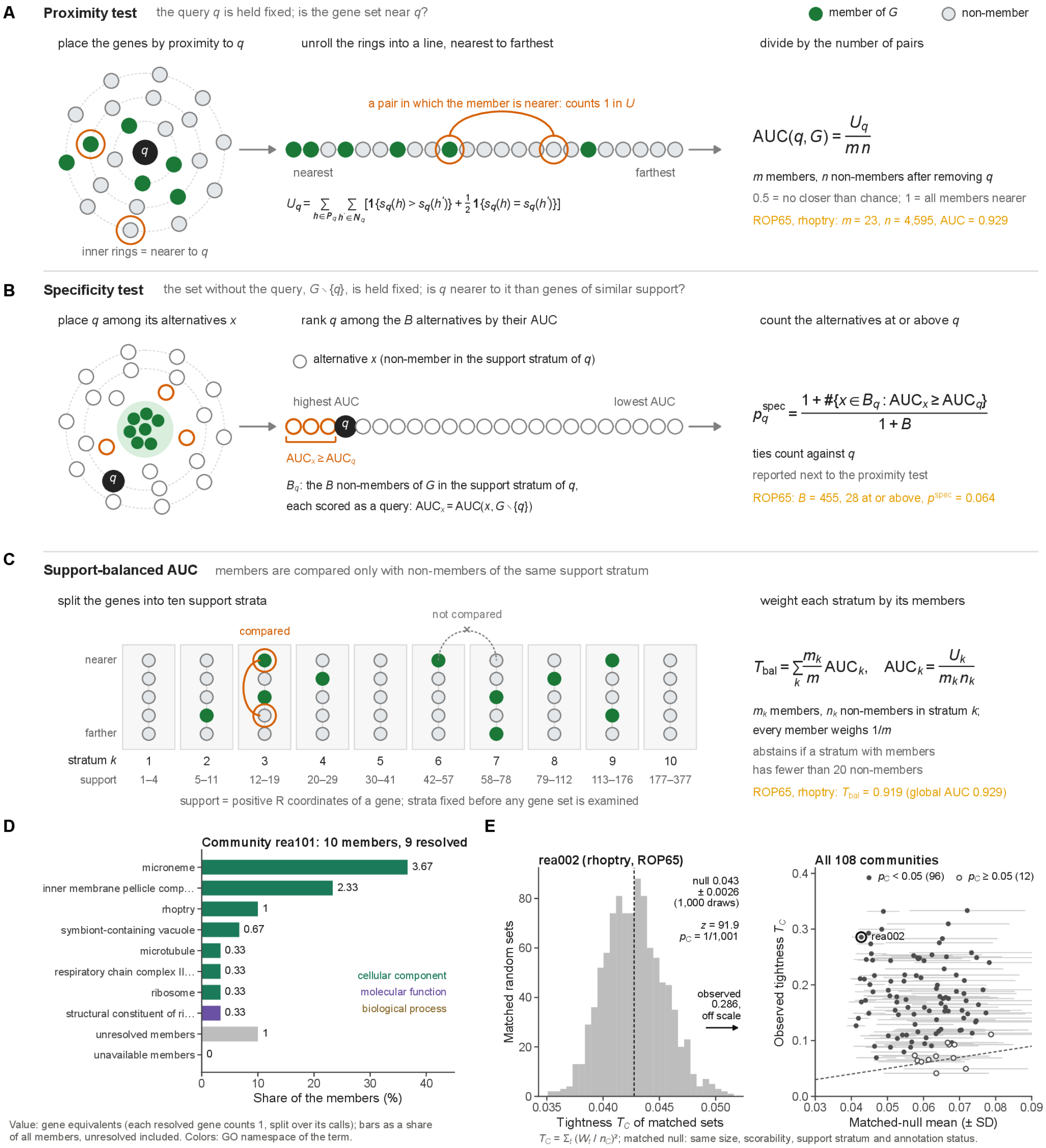


**Figure S24. Community identity (Functional Annotator).**
**(A)** Proximity test. With the query q held fixed, genes are placed by proximity to q and unrolled into a line from nearest to farthest. AUC(q, G) = U_q / (m n) counts the member–non-member pairs in which the member is nearer, over all such pairs. 0.5 means no closer than chance and 1 means all members are nearer. Example: ROP65, rhoptry, m = 23, n = 4,595, AUC = 0.929.
**(B)** Specificity test. With G without the query, G\{q}, held fixed, q is ranked among the B non-members in its support stratum, each scored as a query. p_q^spec = (1 + #{x ∈ B_q : AUC_x ≥ AUC_q}) / (1 + B), with ties counting against q. Example: ROP65, B = 455, 28 at or above, p^spec = 0.064.
**(C)** Support-balanced AUC. Members are compared only with non-members of the same support stratum (ten strata fixed before any gene set is examined). T_bal = Σ_k (m_k / m) AUC_k, and a stratum with members abstains if it has fewer than 20 non-members. Example: ROP65, T_bal = 0.919 (global AUC 0.929).

**(D)** Composition of one community, rea101 (10 members, 9 resolved). Bars give gene equivalents (each resolved gene counts 1, split over its calls) as a share of all members, with unresolved and unavailable members as separate categories. Color marks the GO namespace of each term.
**(E)** Tightness. Left: tightness $T_C$ of the rhoptry community rea002 (ROP65) against 1,000 matched random sets (same size, scorability, support stratum and annotation status). Observed 0.286 lies far outside the null ($0.043 \pm 0.0026$; $z = 91.9$; $p_C = 1/1{,}001$). Right: observed tightness against the matched-null mean (± SD) for all 108 communities. 96 have $p_C < 0.05$; the dashed line is observed = null mean. $T_C = \Sigma_t (W_t / n_C)^2$.

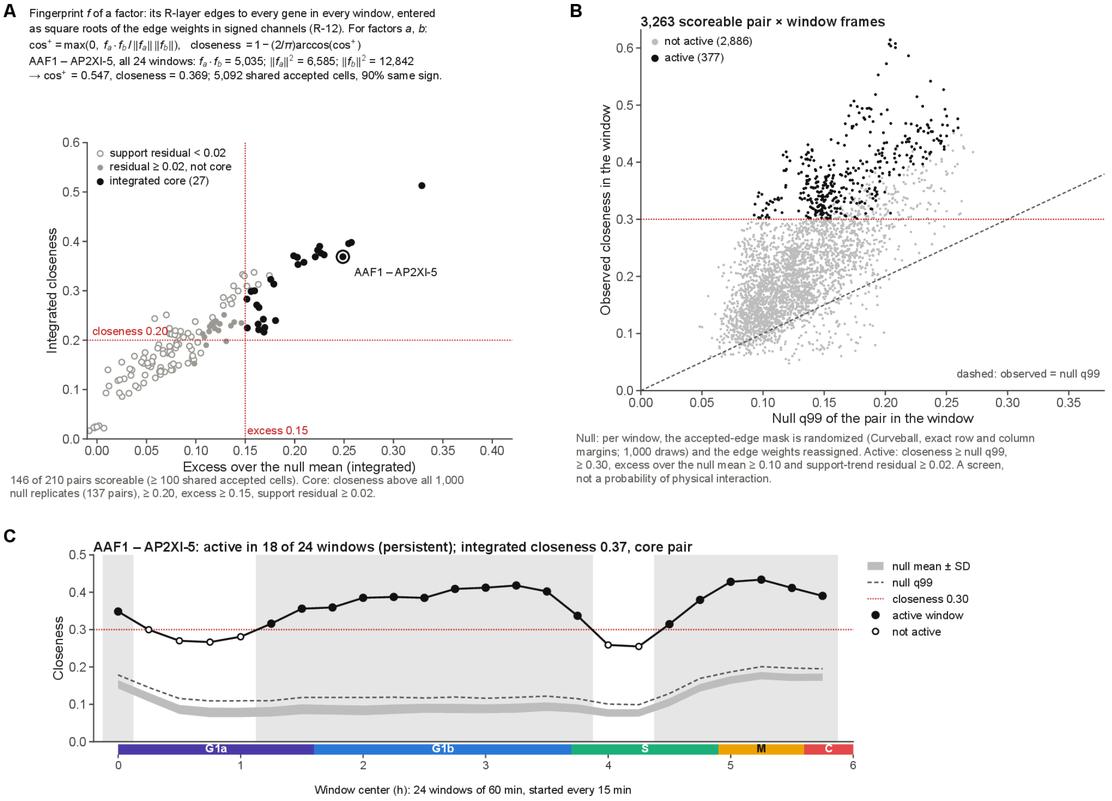


#### Figure S25. Regulator similarity (method for Fig. 3E).

**(A)** Factor fingerprints and closeness. **Fingerprint.** A factor's fingerprint lists its R-layer edges to every gene in every window, entered as square roots of the edge weights in signed channels (square-root R-layer product, R-12). **Closeness.** For two factors, closeness = 1 − (2/π) arccos(cos), where cos is the cosine of their fingerprints (never negative, because both channels are non-negative). **Worked example.** AAF1–AP2XI-5 over all 24 windows, from the stored dot product and squared norms: cos = 0.547, closeness = 0.369. They share 5,092 accepted cells, 90% of the same sign. **Plot.** Integrated closeness against excess over the null mean for the 146 scoreable pairs (at least 100 shared accepted cells). The integrated core (27 pairs, black) requires all of:

- closeness above all 1,000 null replicates;
- closeness ≥ 0.20;
- excess ≥ 0.15;
- support-trend residual ≥ 0.02.

**(B)** Null model and per-window calls. **Null.** In each window, the accepted-edge mask is randomized by Curveball with exact row and column margins (1,000 draws), and the factor's edge weights and signs are shuffled among its randomized targets. **Plot.** Each of the 3,263 scoreable pair × window frames: observed closeness against the null 99th percentile. **Active call** (377 frames) requires all of:

- closeness ≥ null q99;
- closeness ≥ 0.30;
- excess over the null mean ≥ 0.10;
- support-trend residual ≥ 0.02.
- This is a screen, not a probability of physical interaction.

**(C)** Active windows of one pair, AAF1–AP2XI-5, over the 24 windows (60 min, started every 15 min; plotted at the window center). The plot shows observed closeness, null mean ± SD and null q99. Active windows are filled and shaded. The pair is active in 18 of 24 windows (persistent), with integrated closeness 0.37, and is a core pair.

### Supplementary Table Legends

**Table S1. Knockdown phenotype scores.** IMC3 phenotype percentages for the 21 factors after knockdown, with six phenotype categories scored in three replicates and their average.

**Table S2. scRNA-seq quality-control summary.** Single-cell RNA-seq quality-control summary of the 22 libraries, with chemistry, Cell Ranger version, filter thresholds, cell numbers at each filtering stage, median UMIs and genes, BAM-QC counts and Cell Ranger metrics.

**Table S3. Cdc5 and SNIP1 target genes.** CUT&RUN target gene lists of Cdc5 and SNIP1 as three-column overlap lists (Cdc5 only, shared, SNIP1 only) for wild type, and for each factor in the knockdown of the other at 2 h.

**Table S4. Spliceosome donut plot input genes.** Input list of 133 spliceosome-related genes for the donut plot, with descriptions, splicing contributor, conservation complex and interactor sources (Cdc5, SNIP1, GCN5b, BDP1, RES, GSK, AP2IX-7).

**Table S5. PfCdc5-perturbed versus ring DEGs and GO.** Differential expression of PfCdc5-perturbed cells versus rings: the full gene table (2,320 genes), the 23 significant DEGs with transcript and intron counts, and GO enrichment of the upregulated and downregulated genes.

**Table S6. PfCdc5-perturbed versus schizont DEGs and GO.** Differential expression of PfCdc5-perturbed cells versus schizonts: the full gene table (2,571 genes), the 73 significant DEGs with transcript and intron counts, and GO enrichment of the upregulated and downregulated genes.

**Table S7. Daughter-cell budding gene set.** Curated daughter-cell budding set (BUD, 45 genes) with gene names, product descriptions, membership in the 4,619 clustered genes and the gene community of each gene.

**Table S8. AAF1 and AP2XI-5 target genes.** CUT&RUN target gene lists of AAF1 and AP2XI-5 as three-column overlap lists (AAF1 only, shared, AP2XI-5 only) for wild type, and for each factor in the knockdown of the other at 2 h.

**Table S9. AlphaFold 3 AAF1-AP2 pair predictions.** AlphaFold 3 predictions for 156 pairs of AAF1 with AP2 and non-AP2 proteins, with ipTM, ranking score and inter-chain PAE, a group summary and run metadata.

**Table S10. AAF1 pulldown proteomics.** Proteome Discoverer export of the AAF1 pulldown and the RH control pulldown, with a comparison sheet for the scatter plot and a GO term sheet.

**Table S11. Parasite strains.** List of 32 parasite strains (*T. gondii* and *P. falciparum*) with the source of each, as a PMID or this study.

**Table S12. Oligonucleotides.** Oligonucleotides used to generate the knockdown and tagged strains, in five sheets (mAID conditional knockdown, C-terminal tag, N-terminal tag, Tet7Sag4 system and PfCdc5), with gene ID, primer name and sequence.

**Table S13. Antibodies.** Seventeen antibodies with species, dilution or amount for immunofluorescence, CUT&RUN and western blotting, and source and identifier.

**Table S14. Plasmids.** List of 32 plasmids with the source and identifier of each.

**Table S15. Software and versions.** Software, versions and references used in the computational analyses, grouped by computing environment, with 82 rows. References are PMIDs, DOIs or software URLs.

**Table S16. PfCdc5 phenotype titration.** Phenotype titration of PfCdc5 at six concentrations in three replicates over seven time points.

**Table S17. scRNA-seq filter parameters.** Filter parameters and cell numbers of the 22 libraries, including total reads, read minimum, mismatch cap, density features, mixture components, UMI and score floors, and the status of the repeated filter run.

**Table S18. scRNA-seq per-cell tables.** Per-cell tables of the 22 libraries for the 249,877 cells with quality score above 0.9, with a per-library summary sheet. Columns give quality score, doublet calls, phase, pseudotime, time and errors, and UMAP coordinates.

**Table S19. Differential expression time windows.** The 24 time windows of the differential-expression contrasts, forming four sets of six 1 h windows offset by 15 min, with start, end, center label and phase.

**Table S20. Pooled differential expression tables.** Run settings and cell and row numbers of the pooled window and bulk differential-expression tables of the 21 factors, with seeds, partitions and variant. The pooled tables themselves are deposited separately.

**Table S21. Differential expression summary.** Differential-expression summary per factor, with WT and KD cell numbers, genes tested, counts of DEGs by direction in the bulk contrast and along the cycle, and the split of bulk DEGs into direct and indirect targets.

**Table S22. *Plasmodium falciparum* stage markers.** *Plasmodium falciparum* stage-marker genes used for the sc-type stage labels of the PfCdc5 knockdown, from differential expression of each stage against all other cells of the Malaria Cell Atlas (1,448 rows, 1,371 genes). Every listed marker has adjusted P below 0.05, absolute logFC above 1.9 and detection of at least 0.10 in the group where it is higher.

**Table S23. CUT&RUN libraries of the network.** The 21 analyzed (library, control) pairs of the network, one library per factor, all against the IgG control, with metadata, read pairs and length, instrument, QC at q = 0.075 and called peaks over q = 0.025 to 0.125.

**Table S24. CUT&RUN libraries not selected.** The 22 (library, control) pairs not selected for 17 of the 21 network factors, with QC at q = 0.075 compared with the selected library.

**Table S25. CUT&RUN library selection check.** Candidate CUT&RUN libraries per network factor at q = 0.05 and 0.075, and whether the selected library has the most fragments, the highest treatment-to-control signal-to-noise ratio and the most peaks.

**Table S26. Parameter grid and Pareto front.** The 630 networks of the parameter grid with the four thresholds, the three objectives, the network size and a flag for the 61 Pareto-optimal networks.

**Table S27. Consensus network edges.** Consensus network edge list (308,098 edges) with window, factor, target gene, signed weight, effect and support among the 61 front members, plus a per-factor summary. Edges are retained at a support of at least 31 of the 61 front members.

**Table S28. Consensus network sensitivity.** Edges, target genes, factors and communities of the consensus networks built from five Pareto fronts at 33%, 50% and 67% support, plus the support curve of the REA front.

**Table S29. Gene communities.** Gene communities of the consensus network: one sheet per community (108) with size, co-association, silhouette and timing descriptors, and one sheet per clustered gene (4,619) with community, member support and assignment status. 781 clustered genes are unassigned.

**Table S30. Community GO enrichment.** Gene Ontology terms enriched or depleted in the REA communities for the curated and full annotation files, with counts, universe, fold, P value, q values and leaf flag. Only (community, term) pairs with global BH-adjusted q below 0.05 are listed, out of 233,612 tests.

**Table S31. Community hyperLOPIT enrichment.** Enrichment of hyperLOPIT localization classes and compartments in the communities, with counts, fold, Jaccard index, P value and q. Only categories with q below 0.05 within level are listed, out of 4,644 tests.

**Table S32. Binding and knockdown response overlap.** Overlap between the promoter-bound genes of each factor and the genes that decreased or increased in each knockdown, by cell-cycle phase, with enrichment statistics for the same-factor pairs. Significant means Bonferroni-adjusted P below 0.05 over 168 same-factor down and up tests.

**Table S33. Community regulators.** Regulators of each community: the community, factor, time window and mode tests with q below 0.05, and a per-community summary of activating and repressing factors with their significant windows. Tests are selected from 108,864 tests with BH q below 0.05 over all of them.

**Table S34. Functional Annotator calls.** Functional Annotator gene-level calls: retained calls per gene, statistic, catalog and term, and the final top-three calls per gene that feed the community compositions. Only retained calls are listed (AUC above 0.5, q below 0.05, top three), so there is no Significant column.

**Table S35. Community functional composition.** Functional composition of the communities: mass of each called GO term per community and its percentage of all and of resolved members, plus enrichment of the Functional Annotator calls per community and annotation column.

**Table S36. Community tightness summaries.** Summary per community (108) with size, analyzed and resolved members, functional tightness against the matched-null mean, SD, percentile, z and empirical P, the three leading terms, and GO enrichment and cluster-quality summaries. Significant means empirical P below 0.05 (96 communities).

#### Supplementary Movies Legends

**Movie S1. Time-resolved animated regulator-similarity network progression through the *T. gondii* lytic cycle.** Corresponds with **Fig. 3E**.

**Movie S2. Rotation of the AlphaFold-3 ribbon model of the AAF1/AP2XI-5/AP2X-5 complex.** Corresponds with **Fig. 7D**.
